# A third kind of episodic memory: Context familiarity is distinct from item familiarity and recollection

**DOI:** 10.1101/2024.07.15.603640

**Authors:** Richard J. Addante, Evan Clise, Randall Waechter, Jesse Bengson, Daniel L. Drane, Jahdiel Perez-Caban

**Affiliations:** Florida Institute of Technology, Department of Psychology, 150 W. University Dr., Melbourne, FL 32905, USA; Florida Institute of Technology, Department of Biomechanical Engineering, Melbourne, FL 32905, USA; Neurocog Analytics, LLC, Palm Bay, FL, 32905; Windward Islands Research and Education Foundation (WINDREF), Saint George University Medical School, Grenada, West Indies; University of California, Davis, Davis, CA, USA; Emory University, School of Medicine, Atlanta, GA, USA

## Abstract

Episodic memory is accounted for with two processes: ‘familiarity’ when generally recognizing an item and ‘recollection’ when retrieving the full contextual details bound with the item. Paradoxically, people sometimes report contextual information as familiar but without recollecting details, which is not easily accounted for by existing theories. We tested a combination of item recognition confidence and source memory, focusing upon ‘item-only hits with source unknown’ (‘item familiarity’), ‘low-confidence hits with correct source memory’ (‘context familiarity’), and ‘high-confidence hits with correct source memory’ (‘recollection’). Results across multiple within-subjects (trial-wise) and between subjects (individual variability) levels indicated these were behaviorally and physiologically distinct. Behaviorally, a crossover interaction was evident in response times, with context familiarity being slower than each condition during item recognition, but faster during source memory. Electrophysiologically, a Condition x Time x Location triple dissociation was evident in event-related potentials (ERPs), which was then independently replicated. Context familiarity exhibited an independent negative central effect from 800-1200 ms, differentiated from positive ERPs for item-familiarity (400 to 600 ms) and recollection (600 to 900 ms). These three conditions thus reflect mutually exclusive, fundamentally different processes of episodic memory. Context familiarity is a third distinct process of episodic memory.

**Summary:** Memory for past events is widely believed to operate through two different processes: one called ‘recollection’ when retrieving confident, specific details of a memory, and another called ‘familiarity’ when only having an unsure but conscious awareness that an item was experienced before. When people successfully retrieve details such as the source or context of a prior event, it has been assumed to reflect recollection. We demonstrate that familiarity of context is functionally distinct from familiarity of items and recollection and offer a new, tri-component model of memory. The three memory responses were differentiated across multiple behavioral and brain wave measures. What has traditionally been thought to be two kinds of memory processes are actually three, becoming evident when using sensitive enough multi-measures. Results are independently replicated across studies from different labs. These data reveal that context familiarity is a third process of human episodic memory.

**Graphical Abstract:** 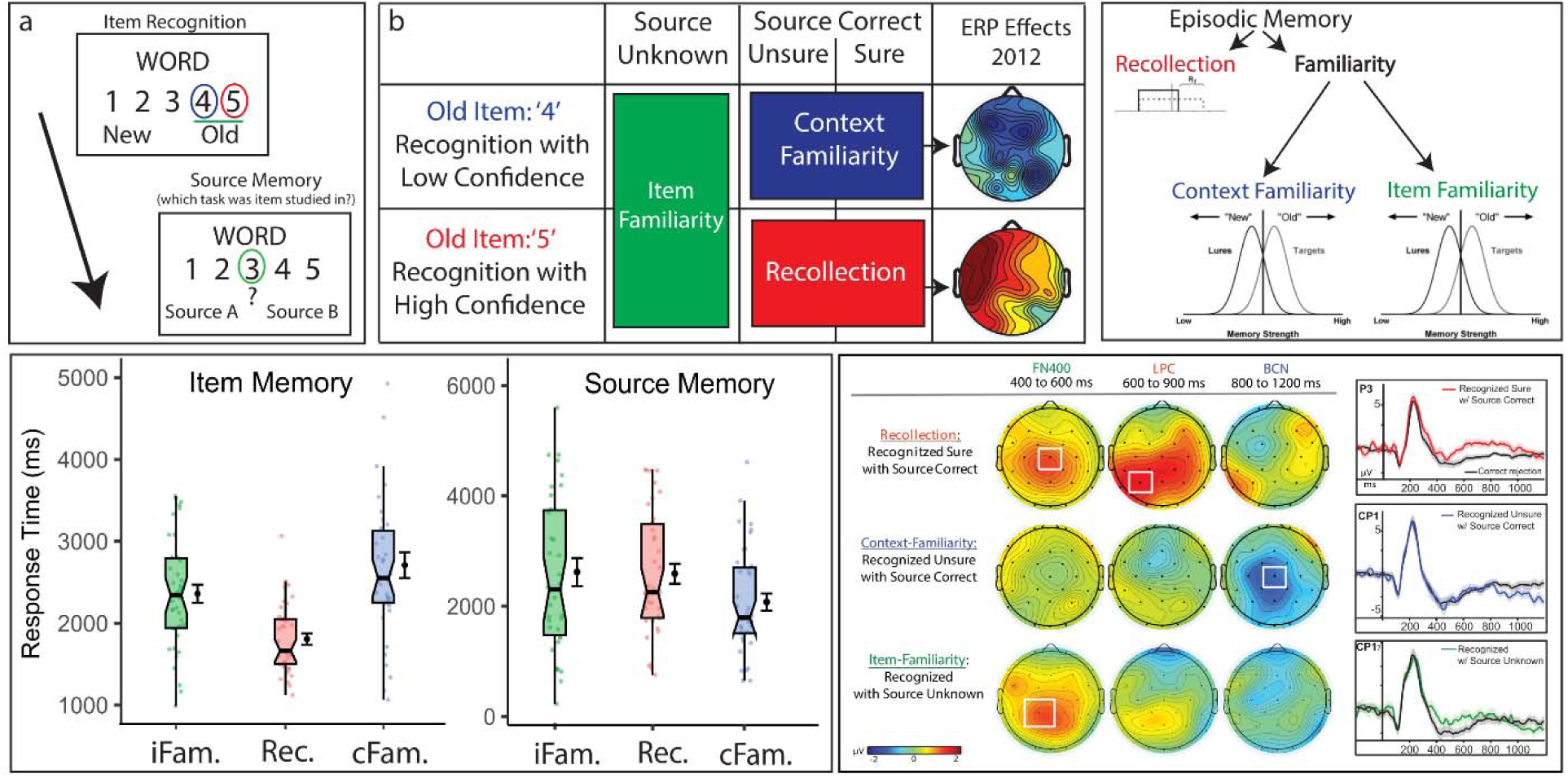

## Introduction

Sometimes people remember information about the source of a prior event correctly (such as background, location, or other kinds of context) without fully recollecting the details of the event ^1–3^. For instance: “this place seems familiar, but I don’t know why or what from” or “I know the general place where I read something [e.g. the bottom left hand part of a page of a book], but I can’t remember what it was [e.g., neither the page nor the specific information]” ^4,5^. It is not clear how this phenomenon occurs in memory. Cognitive models have typically accounted for episodic memory through two processes (for Review, see ^6^): ‘familiarity’ recognizes an item from the past but with varying levels of memory strength and lacking contextual details, whereas recollection retrieves the item bound within the contextual details it was experienced with in the past, such that researchers typically used context retrieval as a proxy for recollection. Familiar recognition of a context in the absence of recollection is thus a puzzle for memory theories because it is difficult to resolve the paradox with just the two processes of recollection and familiarity-but these memories happen nonetheless ^1,3,4,7–14^.

A prior study investigated this conundrum using event-related potentials (ERPs) to focus on the unique memory condition in which people reported low confidence recognition of an item but had accurate source memory for the item’s context ^1^. It was found that familiar context can be retrieved without relying upon recollection, thereby demonstrating that these constructs are dissociable and not dependent (Figure 1).The ERPs associated with this condition were electrophysiologically distinct from those associated with the memory process of recollection. The condition was therefore named ‘context familiarity’, based upon the framework of the Binding-in-Context (BIC) model ^15–17^, which proposes that items and contexts are processed differently in the parahippocampal gyrus of the medial temporal lobe, and that the hippocampus binds the item within the context it was experienced in, e.g. recollection.

**Figure 1.**
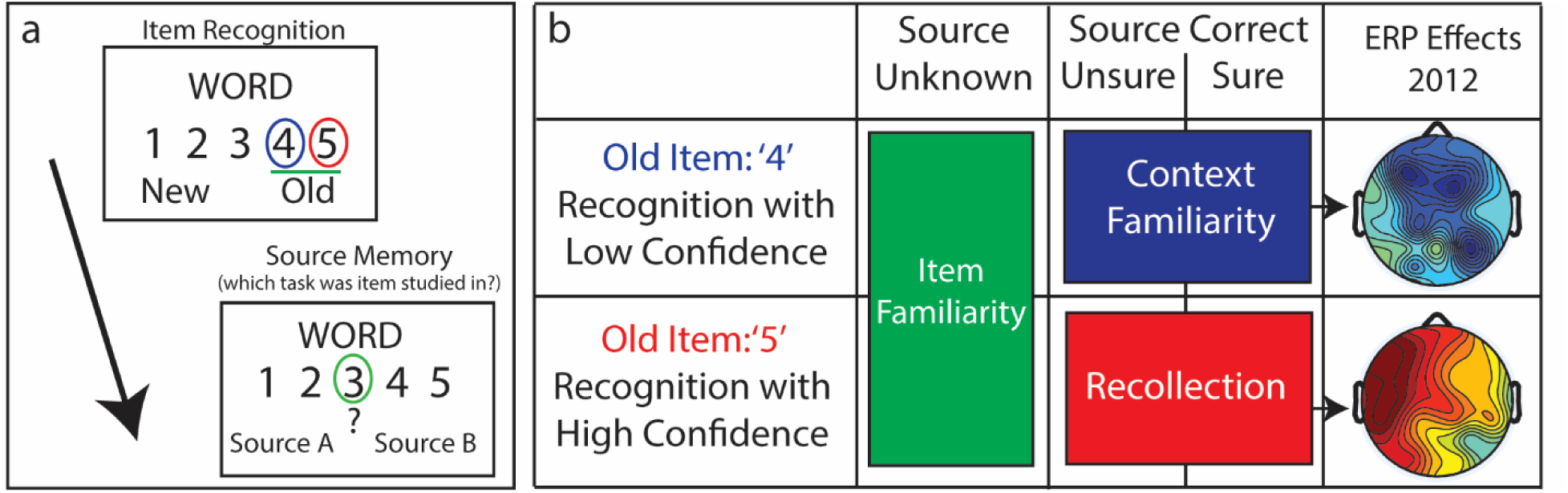
Memory test schematic and combinations of conditions analyzed. Panel A (left): For each of 216 items studied and 108 new item lures, participants responded to a memory test that first asked them to rate their recognition confidence on a scale from 1 to 5 for if they thought the word on the screen was an old item from the study phase or if they thought it was a new item. Immediately following the item recognition judgment, participants were asked to indicate their source memory, e.g. if they remembered which study task the item was encountered in during the study phase. If they could not remember, they were instructed to then press the ‘3’ key to indicate that the source was unknown. Panel B (right): schematic identifying the response profile for the three main conditions of interest analyzed in the current study. Green: item recognition hits collapsed across confidence levels, and which received responses of having no source memory, were operationalized as ‘item familiarity’ (iFam.^66,67^). Blue: low confidence recognition hits (responses of ‘4’) that received correct source memory judgments were operationalized as ‘context familiarity’ (cFam. ^1^). Red: high confidence recognition hits (responses of ‘5’) that received correct source memory judgments were operationalized as ‘recollection’ (Rec. ^1,17,29,32,39^). Right: Representative ERP results of context familiarity and recollection are depicted from the original findings ^1^ (reproduced with permission of the owner), shown as topographic maps of each condition plotted in comparison to correct rejections during the latency of 600 to 800 milliseconds in the memory epoch, with color scale depicting difference amplitudes (from -2 to 2 µV) of each condition as compared to correct rejections.

The ERP effects for this ‘context familiarity’ manifested generally as a widespread fronto-central-parietal negative-going effect. This occurred at approximately 600 milliseconds (ms) after stimulus presentation and extended later in the epoch from 800 to 1200 ms when compared to both correct rejections as well as when compared to incorrect source judgments, too ^1^. This relatively late, broad central negativity (BCN) was dissociated from the late positive component (LPC) effect that is well-associated with recollection and did not show signs of an FN400 effect often related to familiarity. However, that study did not directly address whether the effects of context familiarity were dissociable from those for item familiarity. Therefore, it has remained unknown whether there is one type of overarching familiarity process that includes representations of both item and contexts (e.g. the BIC model), or, if there might be two separable forms of familiarity: one for items and one for context.

As a brief primer, in human long term memory, theoretical models of conscious episodic recognition have been largely governed by the dual cognitive processes of familiarity and recollection ^6,15,17–34^. Recollection is operationalized as the declarative retrieval of episodic information (i.e., an event) specifically with both the item and context of the event bound together into a cohesive retrieval of the episode (for review see ^15^). Recollection is usually associated with the confident retrieval of contextual information surrounding the item of the event (for reviews see ^6,23,33^). The item in the event, however, may be retrieved without recollection, instead relying upon a process of general familiarity of the episodic information, typically conceptualized as declarative, conscious retrieval of an item from a prior episode but without the associated confidence, recollection, or contextual information in which it occurred.

These two memory phenomena are dissociable cognitive processes ^32^, with differential neural substrates in the medial temporal lobes ^35–38^, are neuropsychologically dissociable among patient impairments ^39–41^, and demonstrate spatio-temporally distinct patterns of electrophysiology at the scalp (event-related potentials, ERPs) ^1,42–44^; for Review see ^45,46^. Familiarity has been associated with positive ERP differences in memory trials during a negative-going ERP peak at the mid-frontal scalp sites that occurs at approximately 400 to 600 milliseconds (ms) post stimulus, often called the mid-frontal old-new effect, or FN400 (for frontal-<u>N</u>400 effect). On the other hand, recollection has been associated with positive differences between memory conditions occurring at a peak in the ERP at the parietal region of the scalp from approximately 600 to 900 ms, often referred to as a late parietal component (LPC) or parietal old-new effect ^1,47^ for reviews see ^43,45,46^.

Among studies of episodic recollection, ‘source memory’ has been one of the most reliable methods of assessing recollection for episodic events in both behavioral and cognitive neuroscience paradigms. That is, it has traditionally been assumed that if someone remembers the source of information, then they must have retrieved information about the context of that memory and hence ‘must’ be recollecting that episode. As such, source memory has served as one of the most useful proxies for understanding memory and amnesia deficits ^39^, despite researchers widely acknowledging that source memory is not process-pure as an exclusive measure of recollection ^3,31,48–54^. Prior research has also identified the role that other processes can contribute to source memory ^55^, including item-familiarity ^50,56–58^ and unitization of two items into a new association ^58–63^.

The findings of ERP dissociations for context familiarity and recollection as occurring at different times and different sites on the scalp ^1^ provided direct physiological evidence in support of the BIC model’s theory and related variants ^6,15–17,22,46^; however, there remained limitations. In particular, while those findings had demonstrated that context familiarity was reliably distinct from recollection ^1^, it was unknown if context familiarity is also directly dissociable from item familiarity, or whether it is operating merely as a component element of the same memory process of generic familiarity that governs both items and contextual representations ^15^. The notion that context can be differentially familiar than item information has been suggested in prior studies ^4,5,64^, but was viewed only as context familiarity being merely a variant of the same underlying process of familiarity, and lacked evidence of an independent process until the aforementioned ERP study ^1^. Here, we put forth evidence for a fundamentally different view: that familiarity of context differs qualitatively from item familiarity.

The focus of the present investigation was to assess if the ERP effects for context familiarity can be reliably dissociated from those for item familiarity, and to add novel insights from behavioral measures of response times. When prior investigations observed dissociable ERP activity in time and topography for conditions of recollection (LPC) and familiarity (FN400), it was considered compelling evidence of two independent memory processes ^42,44,45^, that reflect dual-process models of episodic recognition, since a single process would not have (and has not) been able to account for such patterns of neurophysiology without substantial post-hoc modifications ^6,46^. In turn, if the effects of context familiarity are found to be dissociable in ERPs and/or behavior, it could be taken as evidence that context familiarity is a distinct additional process of episodic memory that extends beyond the traditional dual processes of recollection and item-familiarity.

Towards that end, we re-analyzed a published ERP dataset (N = 54) ^65,66^ that utilized the same paradigm as prior studies ^1,39^ and tested a combination of item recognition confidence and source memory judgments. We directly assessed behavioral measures of response times and physiological measures of ERPs for the three conditions of ‘item-only hits with source unknown’ (‘item familiarity’ ^66,67^), ‘low-confidence recognition hits that had correct source memory’ (‘context familiarity’ ^1^), and ‘high-confidence hits that had correct source memory’ (‘recollection’ ^1^) (Figure 1). To preface our findings, we found that context familiarity, item familiarity, and recollection were each dissociable in neural activity patterns occurring at different times and places on the scalp, as well as in behavioral measures of response times for both item and source memory judgments (Experiment 1), and that the electrophysiological findings replicated across several independent studies (Experiment 2).

## Results: Experiment 1

### Behavioral performance

In the current study, when people had low-confidence recognition (item 4 judgments) they went on to provide accurate source discrimination judgments significantly above chance (considered as .50, see Method) (M = .572, SD = .18, SE = .03; t(47) = 2.85, one-tailed p = .003; Cohen’s d = .40, 95% CI [.521, .623]) (Table 1), which was (and replicated) the same level of source memory performance as was previously reported ^1^ (M = .576, SD = .15, SE = .03, 95% CI [.517, .635]). This null difference was quantified by finding no significant differences between the present study and the preceding one ^1^ (t(71) = .081, p = .936, 95% CI [-.080, .087], Bayes Factor_01_ = 3.95). Having established the reproducibility of the prior behavioral performance of memory responses ^1^, we next explored the response speeds and electrophysiological correlates of this paradoxical combination of memory response patterns.

**Table 1.** Proportions of overall response patterns for recognition hits and source judgment combinations for old items during the memory test. Each participant (N = 54) had 216 trials of old items. Left hand column represents item recognition confidence responses, and successive columns denote source memory responses. Top subsection provides the overall proportion of responses of high- and low-confidence recognition hits for each level of source judgment, out of all old trials (214 trials for each of N = 54 participants). Middle subsection indicates the proportion of responses for each source judgment per high- and low-confidence recognition hits; e.g. when subjects did have a recognition hit, it indicated what proportion of times it ended up in each of the source memory categories of interest for the current investigation. Bottom subsection indicates proportion of responses for each source memory judgment out of total recognition hits (collapsing across high and low confidence levels).

| Response Proportions | Source Incorrect |  | Unknown | Source Correct |  | Total |
| --- | --- | --- | --- | --- | --- | --- |
|  | Sure | Unsure |  | Unsure | Sure |  |
| Per total Old Items |  |  |  |  |  |  |
| Item 4: Unsure 'Old' | < .01 | .05 | .02 | .06 | .02 | .16 |
| Item 5: 'Sure Old' | .12 | .07 | .04 | .11 | .30 | .64 |
| Total | .14 | .14 | .22 | .18 | .33 |  |
| Per Confidence Level |  |  |  |  |  |  |
| Item 4: Unsure 'Old' | .05 | .32 | .14 | .38 | .10 | 1.00 |
| Item 5: 'Sure Old' | .19 | .11 | .07 | .17 | .47 | 1.00 |
| Per Total Hits |  |  |  |  |  |  |
| Item 4: Unsure 'Old' | .01 | .07 | .03 | .08 | .02 | .20 |
| Item 5: 'Sure Old' | .15 | .09 | .05 | .13 | .37 | .80 |
| Total | .16 | .15 | .08 | .21 | .39 |  |

### Response Times: Is context familiarity behaviorally dissociable from recollection and item-familiarity?

We analyzed only subjects who had provided pairwise responses in each of the six conditions of interest (N=38). In our paradigm, participants first made a recognition judgment immediately followed by a source memory judgment for each item (Figure 1); we separately assessed the response times for the item and source judgments (Table 2).

**Table 2.** Descriptive Statistics of Response Time (ms) for Item and Source Memory Judgements. Time is reported in milliseconds, and data represents the sample of participants with pairwise observations included in the reported results (N = 38).

| Condition | Item Memory |  |  | Source Memory |  |  |
| --- | --- | --- | --- | --- | --- | --- |
|  | *M* | *SD* | *SE* | *M* | *SD* | *SE* |
| Recollection | 1810 | 422 | 68 | 2596 | 1126 | 182 |
| Context Familiarity | 2710 | 962 | 156 | 2075 | 968 | 157 |
| Item Familiarity | 2361 | 684 | 111 | 2612 | 1577 | 255 |

First, in item recognition judgments, reaction times were assessed across the factor of the three memory conditions (recollection, context familiarity, and item familiarity), using the same low confidence source memory responses in recollection and context familiarity conditions as was done in prior studies ^1^, since it held constant the variable of source memory confidence. Response times for the three memory conditions were subjected to a one-way repeated measures ANOVA with three levels, which revealed a significant main effect of condition (F(2,37) = 25.16, p < .001, *n^2^_p_* =.41). Post hoc pairwise comparisons using the Bonferroni correction revealed that responses significantly differed for item familiarity and context familiarity, such that context familiarity was responded to reliably slower than item familiarity (MD = 348.57 msec, p = .048, 95% CI [2.26, 694.88], Cohen’s d = .48, as well as recollection (MD = 900.15 msec, p < .001, 95% CI [547, 1253], Cohen’s d = .76) (Figure 2). Item familiarity responses were also slower than recollection (MD = 551.58 msec, p < .001, 95% CI [297.97, 805.19], Cohen’s d = 1.24).

**Figure 2.**
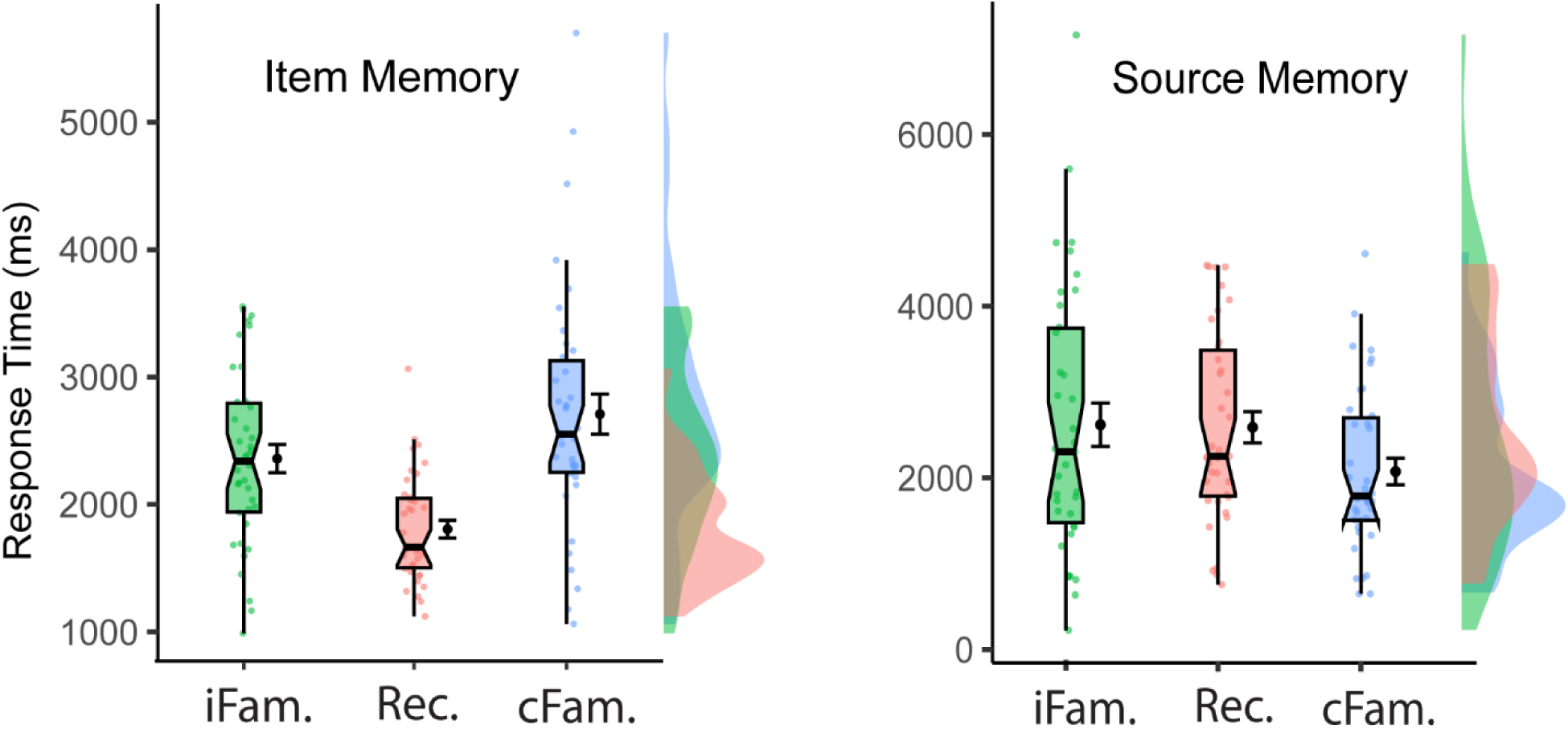
Reaction speeds for item recognition and source memory judgments. Mean times in milliseconds are shown in black with standard error of the mean for responses of item recognition (left) and source memory judgments (right), for three different conditions of memory response patterns. Green: item familiarity (iFam.), operationalized as retrieval of recognition hits that did not have source memory (received responses of ‘source unknown’). Blue: context familiarity (cFam.), operationalized as retrieval of low-confidence item recognition that had correct source memory responses. Red: recollection (Rec.), operationalized as retrieval of high-confidence item recognition that had correct source memory responses. The boxplots demonstrate the median value, notches (95% confidence interval), and interquartile range (IQR) with lower and upper whiskers (Q1-1.5*IQR; Q3+1.5*IQR). The violin plots show density curves centered around the median value. Plots were generated using the scripts created by ^230^.

Within source memory judgments, response times for the three memory conditions revealed a significant main effect of condition (F(1.51,55.81) = 8.28, p = .002, *n^2^_p_* = .18). Further investigation using post hoc tests with Bonferroni correction revealed that source memory judgments for context familiarity responses were faster than those for both recollection and item familiarity (MD = 521.06, p < .001, 95% CI [273.01, 769.10], Cohen’s d = .42; MD = 537.21, *p* = .012, 95% CI [100.20, 974.23], Cohen’s d = .43). (Figure 2). Response times during source memory judgments for recollection and item familiarity conditions did not differ (MD = 16.16, p = 1.00, 95% CI [-399.64, 431.96], Bayes Factor_01_ = 5.70).

Because item- and context-familiarity conditions also differed in recognition confidence (item familiarity included high and low confidence responses, whereas context familiarity included only low confidence recognition responses [Method, Figure 1), it was possible that differences between item- and context-familiarity conditions could have been attributable to the high confidence (‘5’) recognition responses included in the item familiarity condition. To test this directional hypothesis, we conducted a targeted analysis of these conditions while using only ‘low confidence recognition (‘4’) with source unknown’ (item familiarity_Low,_ no source memory responses), and low confidence recognition with source correct (context familiarity_Low_, all correct source memory responses)^1^. While this excluded high-strength item-familiarity responses (high confidence recognition responses of ‘5’), such that a null finding would not be able to falsify the hypothesis, it nevertheless represented a valuable challenge of the alternative hypothesis using only weaker-strength item familiarity matched for item-familiarity strength ^28,29,32,66,68^. Since these control analyses’ conditions were specifically defined by removing response types and thereby inherently weakening its power to detect differences, they nevertheless still produced significant results, and they also went on to be mirrored by similar results in the physiological measurements

The reliable difference originally observed between item- and context-familiarity responses was still preserved (t(36) = 2.09, p = .022, one-tailed, Cohen’s d = .34 95% CI [-inf, - 49.49]; cFam_Low_: M = 2703, SD = 1033, SE = 149; iFam_Low_: M = 2896, SD = 852, SE = 140), and maintained a comparable pattern for source memory responses as were seen in the main analysis above (t(36) = 1.94, p = .030, one-tailed, Cohen’s d = .32, 95% CI [-inf, 57.76]; iFam_Low_: M = 2418, SD = 1697, SE = 279). These findings ensured that differences between familiarity responses (context, item) were not confounded by inclusion of higher confidence levels of memory strength in the item familiarity condition.

### Electrophysiological Results

The EEG analyses began with an assessment of general memory ERPs (hits vs correct rejections) to identify if, consistent with the behavioral findings, there were three different physiological patterns of activation supporting episodic memory retrieval. This adopted the same approach as originally taken to characterize the conventional dual processes of recollection and familiarity ^44,69^ but was applied here to three different locations and latencies selected based upon hypotheses from prior findings ^1,39^. This was then followed by a series of more specific analyses of the particular memory condition contrasts.

For the first analysis, we used a basic contrast of measuring ERPs for hits vs. correct rejections in every subject (N=54) using a priori predictions from the same sites as the previous reports ^1,39^: mid-central site Cz from 400-600 ms for the FN400, left parietal site P3 from 600-900 ms for the LPC, and fronto-central site Fc1 from 800-1200 ms for the BCN. These ERPs were subjected to a 2×3×3 repeated measures ANOVA with factors being memory, time, and site. Results revealed main effects of site (F(1.20, 53) = 46.71, p < .001, *n*^2^ = .47) and time (F(1.6, 53) =13.271, p <.001, *n^2^_p_* =.20) and, importantly, a significant 3-way interaction of memory x site x time (F(2.29, 53) = 20.018, p < .001, *n^2^_p_* =.27) [in addition to interactions of memory x site (F(1.57, 53) = 13.60, p < .001, *n^2^_p_* = .20), memory x time (F(1.61, 53) = 22.89, p < .001, *n^2^_p_* =.30) and site x time (F(1.75, 53) = 14.58, p < .001, *n*^2^ = .22) (Figure 3).

**Figure 3.**
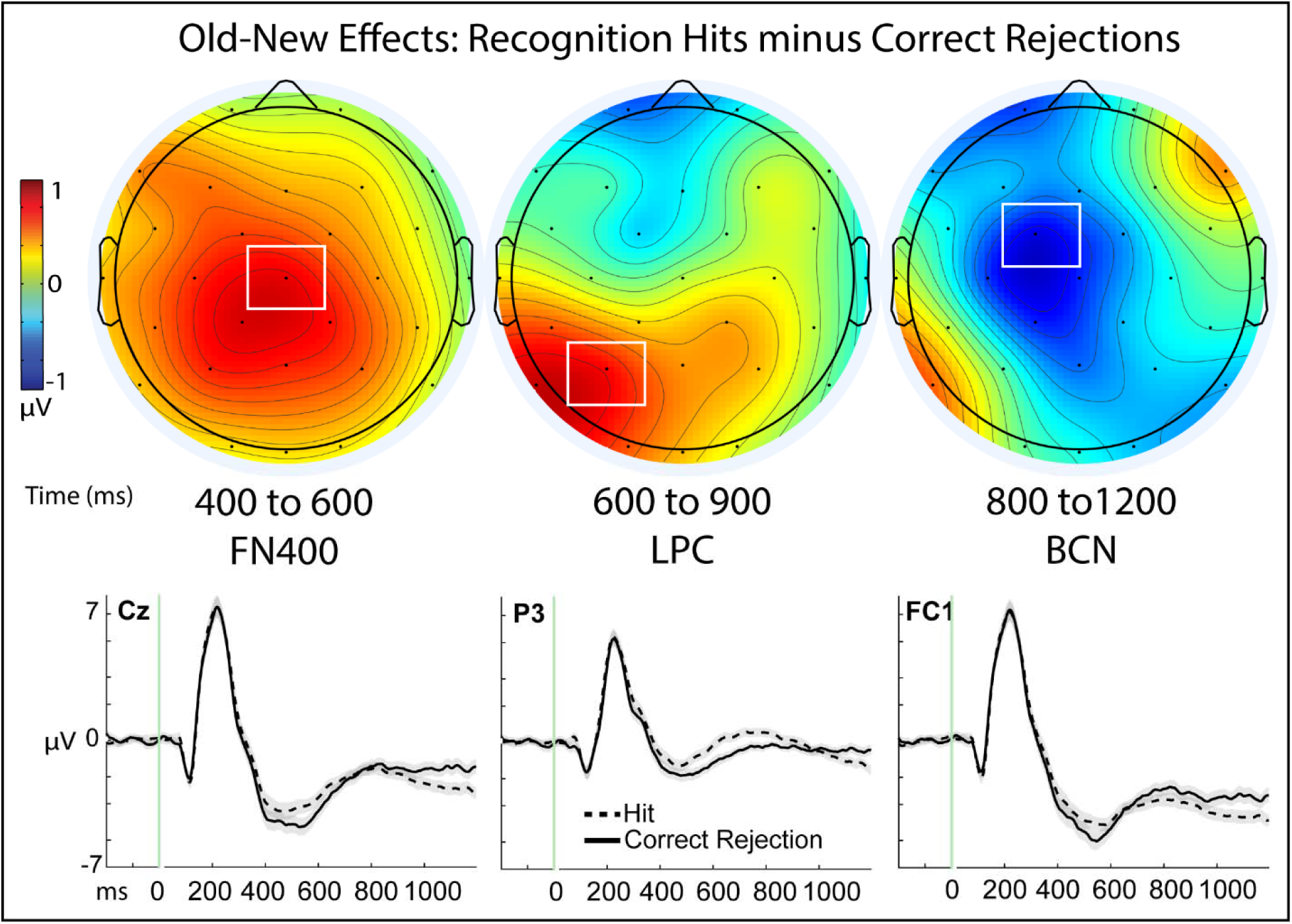
Old-New Effects of Recognition Memory. Top panel: Topographic maps depicting memory effects of the subtraction of hits minus correct rejections; color scale of topographic maps denotes amplitude differences from -2 to 2 mu-volts. Maps are plotted for each of the three time latencies of 400 to 600 milliseconds (ms), 600 to 900 ms, and 800 to 1200 ms that have been associated with the frontal negative old-new effect (FN400), late parietal component (LPC), and later broad central negativity (BCN), respectively. White boxes denote the sites from the topographic maps that are shown for the corresponding ERP figures beneath the maps, as well as where statistical analyses were performed. Bottom panel: Event-related potential (ERP) amplitudes (µV) of the individual conditions of recognition hits (dashed line) and correct rejections (solid line) plotted through the 1200 ms epoch and beginning with a -200 ms baseline period. Shaded areas depict the standard error or the mean for each condition.

Consistent with our a priori directional hypotheses, hits were more positive than correct rejections at Cz from 400-600 ms (i.e. an FN400 effect; t(53) = 3.86, p < .001, one-tailed, Cohen’s d = .53, 95% CI [.47, inf]; Hit: M = -3.66, SD = 3.51, SE = .48; CR: M = -4.50, SD = 3.72, SE = .51) and at P3 from 600-900 ms (an LPC effect; t(53) = 3.10, p = .002, one=tailed, Cohen’s d = .42, 95% CI [.38, inf], Hit: M = 3.46, SD = 2.54, SE = .346; CR: M = -.430, SD = 2.28, SE = .310), but were more negative at fronto-central site Fc1 from 800-1200 ms (t(53) = 2.84, p = 003, one-tailed, Cohen’s d = .39, 95% CI -[inf, -.348]; Hit: M = -3.86, SD = 3.31, SE = .451; CR: M = -3.01, SD = 3.28, SE = .45) with comparable results also evident from 800-1200 ms at adjacent site Cz, t(53) = 2.50, p = .007, one-tailed, Cohen’s d = .34, 95% CI [-inf, -.242]; Hit: M = -2.27, SD = 3.08, SE = .42; CR: M =-1.54, SD = 3.11, SE = .423). These findings provide strong support for three dissociable neural signals underlying basic episodic recognition, operating in different times and places on the scalp ^44,45,57,70–73^. A series of subsequent analyses were then carried out to systematically characterize each condition (context familiarity, recollection, and item-familiarity) as compared to correct rejections ^44,46,69^, described below.

### Experiment 1: Verifying if recollection and context familiarity exhibit different neural correlates

Since the current project hypothesized that context familiarity is distinct from recollection, we first sought to establish that the dataset validly reproduced prior findings ^1^ before endeavoring to explore differences within familiarity of items and context. A sample of N = 27 subjects provided observations in both conditions of interest for this analysis and satisfied inclusion criteria for numbers of valid ERP trials (see Method). A 2×3×2 repeated measures ANOVA for factors of memory condition (recollection, context familiarity), time interval (400-600ms, 600-900ms, and 800 1200 ms), and site (P3, Cz) was performed based upon a priori predictions derived from the preceding analyses and prior reports dissociating context familiarity from recollection-based processing ^1^. Results revealed a significant main effect of memory condition (F(2,26) = 9.418, p = .005, *n^2^_p_* = .27), a main effect of time interval (F(1.138, 26) = 6.262, p = .015, *n^2^_p_* = .19), and a main effect of site (F(1,26) = 15.89, p < .001, *n^2^_p_* = .38), along with a significant time by site interaction (F(1.35, 26) = 18.81, p < .001, *n^2^_p_* =.42) and a memory x time x site interaction (F(1.5), 26 = 7.69, p = .002, *n^2^_p_*= .23) (Figure 4).

**Figure 4.**
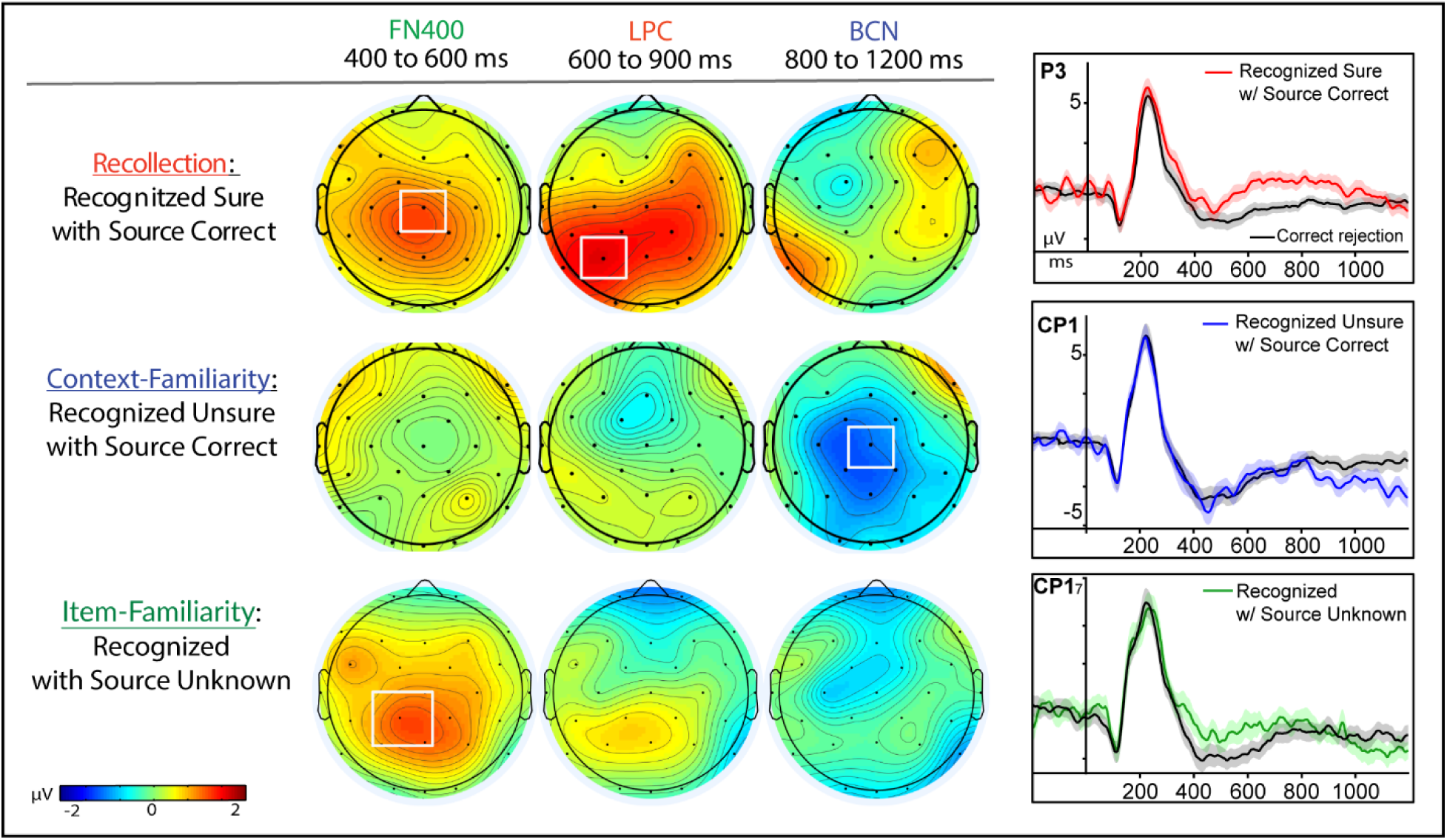
Three physiological effects of episodic memory retrieval. Left panels: Topography maps are shown for each memory condition as compared to correct rejections in Experiment 1: (top) high-confident item recognition judgments that also had correct source memory retrieval; (middle) low-confident item recognition judgments of that also had correct source memory retrieval; and (bottom) recognition hits (collapsed across both high and low confidence) that did not have source memory retrieved (received responses of ‘source unknown’). Color scale of topographic maps denotes amplitude differences for each condition as compared to correct rejections, ranging from -2 to 2 mu-volts. Right panels: Event-related potential (ERP) amplitudes (µV) from representative sites of maximal activation for each of the respective effects during the memory retrieval epoch of -200 to 1200 milliseconds (ms); shaded areas for each plot represent the standard error of the mean.

ERPs for context familiarity compared to the conventional ERP control condition of correct rejections revealed reliable evidence of the same negative-going effect from 800-1200 ms that had been directionally hypothesized from previous reports ^1,39^ (Figure 4), while also reproducing the prior findings of there being no evidence for ERP correlates of recollection or familiarity (LPC, FN400) ^1^. ERPs for context familiarity were more negative-going than correct rejections (CR) from 800-1200 ms at mid central site Cz (t(26) = -2.53, p = .009, one-tailed; Cohen’s d = .49, 95% CI [-inf, -.369]; cFam: M = -2.91, SD = 3.575, SE = .69; CR: M = -1.772, SD = 3.72, SE = .72). Alternatively, there was no LPC effect evident for context familiarity from 600-900 ms (t(26) = 0.161, p = .437, one-tailed; Cohen’s d = .03, 95% CI [-.732, inf, Bayes Factor_01_ = 4.32]; cFam.: M = -.857, SD = 3.00, SE = .58; CR: M = -.932, SD = 2.15, SE = .413), nor any evidence for an FN400 effect for context familiarity from 400-600ms (t(26) = -0.056, p = .522, one-tailed, Cohen’s d = .01, 95% CI [-.753, inf]), Bayes Factor_01_ = 5.12; cFam: M = -5.241, SD = 3.83, SE = .737; CR: M = -5.217, SD = 4.03, SE = .78). ERPs for the recollection condition were also found to be consistent with the prior findings ^1^ when compared to correct rejections (Figure 4): eliciting both an FN400 effect at Cz from 400-600ms, (t(26) = 2.82, p = .004, one-tailed, Cohen’s d = .54, 95% CI [.487, inf]; rec: M = -3.939, SD = 3.54, SE = .68; CR: M = -5.22, SD = 4.03, SE = .78) and an LPC at left parietal site of P3 from 600-900ms (t(26) = 2.98, *p* = .003, one-tailed, Cohen’s d = .57, 95% CI [.665, inf]; rec: M = .627, SD = 2.94, SD = .57; CR: M = -.932, SD = 2.15, SE = .41) but no evident differences during the later time of 800 to 1200 ms (t(26) = .002, p = .499, one-tailed; Cohen’s d= .00, 95% CI = [-.923, inf], Bayes Factor_01_ = 4.91). These results indicated that the neurophysiology for conditions of recollection and context familiarity were indeed distinct in time and place on the scalp. Overall, these results thus reproduced the prior findings that neural processing for the retrieval of familiar contexts was distinct from recollection ^1^, and established the foundation for exploring if the context familiarity may also be directly dissociable from item familiarity.

ERPs for the item familiarity condition also reliably exhibited their predicted effects of an early old-new effect from 400-600 ms, maximal at site Cp1 (N = 19, t(18) = 2.02, p = .029, one-tailed, Cohen’s d = .46, 95% CI [.160, ∞]; iFam: M = -1.32, SD = 3.47, SE = .797; CR: M = -2.47, SD = 2.57, SE = .59), providing concurrent validity with similar other findings of posterior distributions attributed to absolute familiarity ^74–76^. The condition did not exhibit any evidence for an LPC from 600-900 ms at P3 (t(18) = 1.22, p = .119, one-tailed, Cohen’s d = .28, 95% CI [-.238, ∞], Bayes Factor_01_ = 1.27; iFam: M = -5.45, SD = 3.49, SE = .802; CR: M = -1.11, SD = 2.37, SE = .543) nor any later negative effect from 800-1200 ms at Cz (t(18) = .910, p = .187, one-tailed, Cohen’s d = .21, 95% CI [∞, .576], Bayes Factor_01_ = 1.83; iFam: M = -1.93, SD = 3.60, SE = .83; CR: M = -1.29, SD = 2.92, SE = .67). Together, the consistent findings of distinct ERP effects of an FN400, LPC, and BCN among both the established general conditions (Hits vs. CR) and the more specific conditions of response combinations (iFam, cFam, Rec., each vs Cr.) provided convergent validity of similar findings observed from different measures, while also demonstrating concurrent validity of the newer specific ERP measures of memory processing with the more established general ERP measures of memory processes.

### Are the neural correlates of context familiarity different from item familiarity?

To test whether the context familiarity condition reflected the same or different kind of familiarity processing as traditional item-familiarity, we investigated if the two conditions varied during the conventional time window traditionally reported for familiarity-based processing of the FN400 (300-500 ms ^44,45^), performing a within-subjects (N=11) targeted analysis on mid-central site Cz where prior analysis identified peak activation for these conditions in both the present and prior study ^1^ that predicted a one-directional difference of context familiarity being more negative than item familiarity. ERPs for context familiarity were revealed to be significantly more negative than those for item familiarity (t(10) = -2.57, p = .015, one-tailed, Cohen’s d = .76, 95% CI [-inf, - .622]; cFam: M = -4.62, SD = 3.32, SE = .70; iFam: M = -2.42, SD = 3.22, SE = 1.33; Figure 5A), and topographies indicated a broad negative-going distribution of this effect across central scalp sites and throughout the epoch. To assess if these effects varied throughout the epoch a 2×3 ANOVA was performed using factors of memory condition (item familiarity, context familiarity) and time interval (300-500, 600-900, 800-1200 ms). Results revealed a marginal main effect of memory (F(1, 10) = 4.251, p = .066, *n^2^_p_* = .298, Cohen’s d = .65), a significant main effect of time (F(1.18, 10) = 5.13, p = .039, *n^2^_p_* = .339), and no memory x time interaction (F(1.34, 10) = 1.48, p = .255).

**Figure 5.**
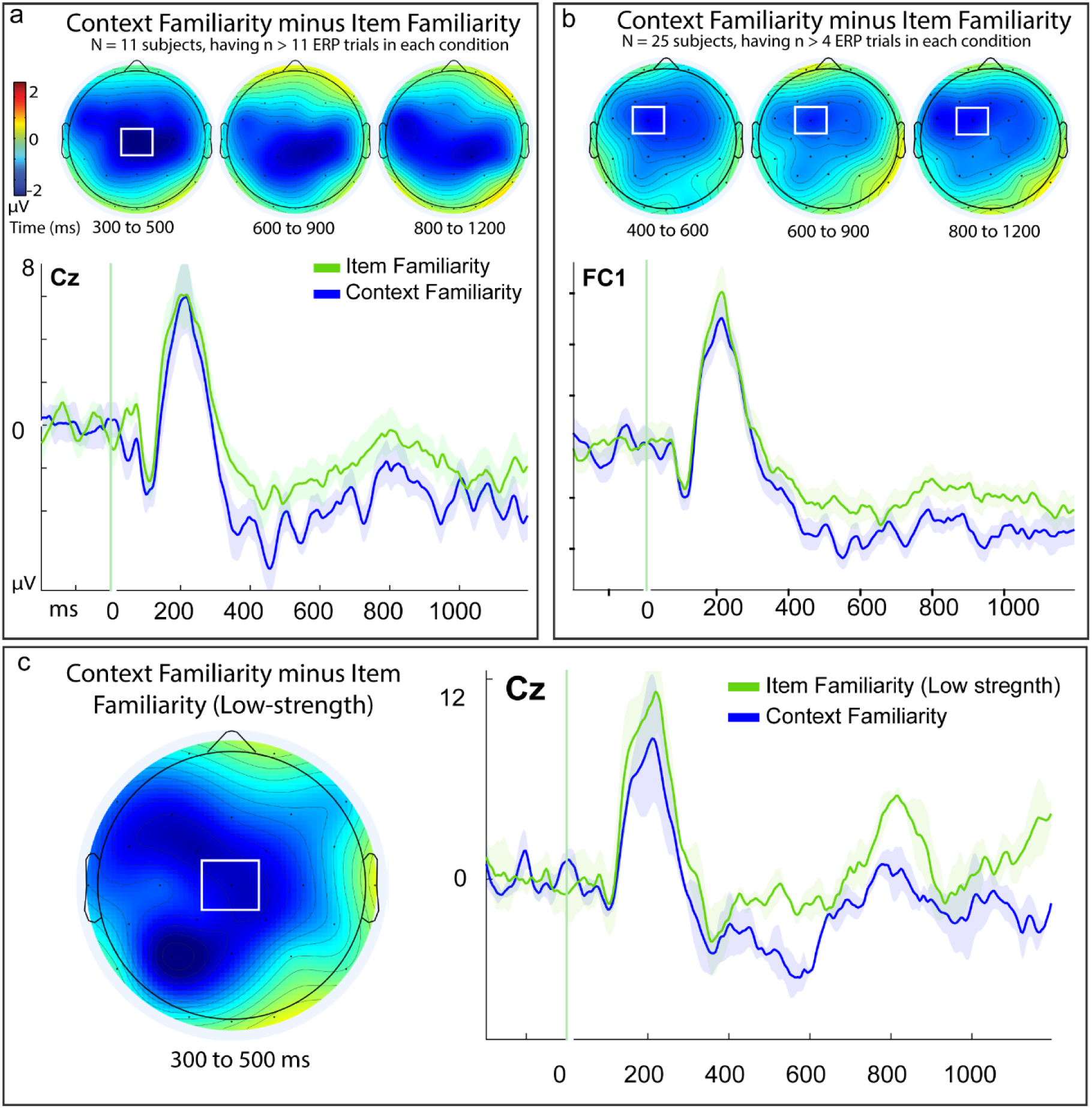
Within-subjects ERP differences of context familiarity and item familiarity. Results depict findings from Experiment 1. Panel A: within-subject effects for participants who shared pairwise response trials in both conditions of item familiarity and context familiarity (N = 11; note: N = subjects, n = trials, see Methods section 2.3.2). Effects were evident at 300-500 ms and distributed broadly across the scalp while being maximal over central midline site Cz. Topographic maps of difference waves for effects results from the subtraction of item familiarity from context familiarity conditions, respectively, with color scale depicting difference amplitudes from -2 to 2 mu-volts; white boxes denote regions of peak activity and targeted analyses; time intervals for each topography map is noted beneath it in milliseconds (ms). Shaded areas of event-related potentials (ERPs) depict the standard error or the mean for each condition. Panel B: Results from a sample of participants (N = 25) meeting a lower inclusion criterion for signal-to-noise ratio (n = 5 artifact-free ERP trials pairwise). Panel C: Control analysis of the same conditions but while holding constant the variable of item recognition confidence (depicting results from only the low confidence ratings of ‘4’ for each condition, and excluding the high confidence ‘5’ responses from item familiarity, N = 3, n > 11).

To test if these results could be attributable to the potential confound of high confidence recognition responses included in the item familiarity condition, we sought a similar control test as was described for the behavioral data of response times: by removing the high-confidence responses from the item-familiarity condition so that it was matched to the low-strength responses of the context familiarity condition^2^. ERPs for this contrast were available from only a limited sub-sample of the primary dataset due to there being only N = 3 participants available meeting our strict a priori inclusion criteria of the ‘item4+source unknown’ item familiarity condition ^65^ (see Methods section 2.3.1) (though note that the present paradigm has repeatedly been shown in prior studies to be effective and sensitive to detecting reliable differences in similar small sample sizes of specialized clinical patients on ERP measures of familiarity, confidence, and implicit memory repetition effects ^39,66,77^, in addition to conditions of metacognition ^65^, and that subsequent analyses conducted in Experiment 2 obtained the same results using a larger sample size of N =12 in an independent data set).

We used a directional t-test based upon the hypothesis that context familiarity exhibits more negative-going ERPs than item familiarity at the same mid-central size (Cz) and time (300 to 500 ms) that was found in the primary analysis (Figure 5C). Despite the limited sample, ERPs for context familiarity_Low_ remained more negative than those for item familiarity_Low_: while this difference was not initially significant from 300-500 ms (t(2) = 1.67, p = .119, one-tailed, Cohen’s d = .962, 95% CI [-∞, -1.37], Bayes Factor_01_ of .610; iFam: M = -1.67, SD = 3.99, SE = 2.30; cFam: M = -3.49, SD = 2.19, SE = 1.27) there was a significant effect maintained in the nearly-identical time interval of 400-600 ms that has also been commonly used for measuring the FN400 in prior studies using this specific dataset ^65,66^: (t(2) = 5.57, p_bonf_ = .015, one-tailed, corrected, Cohen’s d = 3.21, 95% CI [-∞, -1.5]; iFam: M = -1.36, SD = 2.38, SE = 1.38, values: - 3.29, -2.03, 1.31; cFam: M = -4.51, SD = 1.54, SE = .89, values: -5.40, -5.404, -2.73) (Figure 5C), indicating that the main effect was preserved in the more stringent control analysis.

Since we used a relatively conservative criteria for inclusion of participants in the within-subjects analysis (n = 12 artifact-free ERP trials pairwise, see Method), it provided a rigorous threshold to foster better signal-to-noise ratio of ERP effects but also resulted in a fairly small sample (N = 11). Therefore, we also performed a parallel analysis with a more liberal inclusion criteria for subjects^3^, which could increase the sample size and challenge the findings via testing them with reduced signal-to-noise ratio ^78,79^. The same results persisted when lowering the trial inclusion criteria to n = 5, which provided a larger sample of N = 25 subjects: ERPs for context familiarity still exhibited a negative difference from ERPs of item familiarity that was maximal at adjacent site Fc1 and extending broadly across bilateral mid-central regions (Figure 5B). We used a similar 2×3 ANOVA as employed in the primary analysis, using factors of memory condition (item familiarity, context familiarity) and adjacent time interval (400-600, 600-900, 800-1200 ms), and like the primary results using the more conservative inclusion criteria with a smaller sample, there was a main effect of memory (F(1, 24) = 4.37, p = .047, *n^2^_p_* = .15, Cohen’s d = .42) with no significant evidence for effects of time nor a memory x time interaction (both F’s < 1), indicating that ERPs for context familiarity were again significantly different from item familiarity beginning early in the epoch and persisting throughout despite lower signal-to-noise ratio ^78,79^. Overall, the convergence of findings indicated that ERPs for item familiarity and context familiarity differed reliably during the traditional time window of familiarity processing, and thus that these two memory conditions represented reliably different kinds of familiarity processing on a trial-wise level within subjects ^44,45,57,70–73^.

As the preceding analysis demonstrated moment to moment differences of modular cognitive processes within the same individuals over time, we next sought to determine if these patterns would persist as individual characteristics across individuals ^65^. The within-subjects approach used above required paired observations in both conditions of infrequent memory response patterns; however, the two conditions of item-familiarity and context-familiarity tend not to co-occur often within subjects. That is, those who tend to retrieve source memory well (i.e. better performers) also rarely report lacking source information (lesser memory), and those who tend to fail retrieving source memory do not also frequently report retrieving it. We thus utilized a between-subjects approach that treated those with sufficient trials in one condition but not the other as separate groups, in a mutually independent way (total sample of N = 26; item familiarity group: N = 9, context familiarity group = 17). This resulted in the exclusion of those from the within-subject analysis, but nevertheless converged to provide the same findings with an entirely different subset of participants, despite the inherent challenges to the hypothesis from a smaller sample.

Using a between-groups t-test, the same pattern of ERPs persisted between subjects from 300-500 ms such that ERPs for context familiarity were significantly more negative-going than from item familiarity (t(24) = 2.82, p = .005, one-tailed, Cohen’s d = 1.16, 95% CI [1.93, inf], Figure 6A), (cFam: N = 17, M = -4.62, SD = 4.63, SE = 1.23; iFam: N = 9, M = .250, SD = 3.34, SE = 1.12), and was not statistically significant during the later time intervals (600-900 ms: t(24) = 1.21, p = .119, one-tailed; 95% CI [-.698, inf], Bayes Factor_01_ = 1.57; 800-1200 ms: t(24) = .664, p = .257, 95% CI [-1.72, inf], Bayes Factor_01_ = 2.27). Overall, this mirrored the results from the preceding within-subjects analysis, independently demonstrating that these are distinct physiological patterns for familiar memory responses both at the single trial level (within-subjects) and at the level of individual variability (between-subjects), and that the context familiarity condition cannot be attributed to merely a form of the same kind of familiarity process supporting familiarity of item recognition.

**Figure 6.**
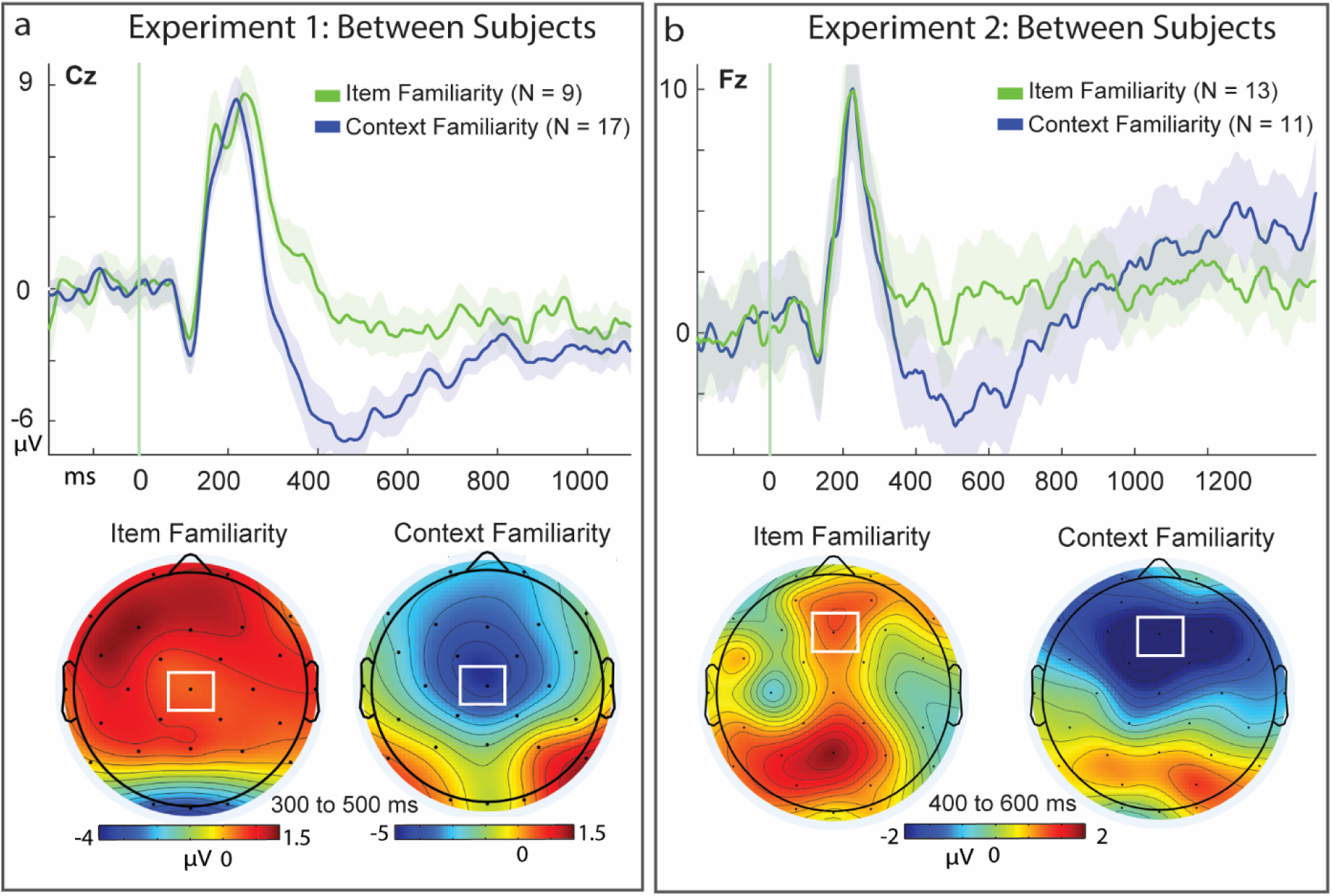
Between-subjects ERP differences of context familiarity and item familiarity. Panel A, top: Between-subject effects in independent sub-groups of participants who had response trials in one condition but not the other (i.e.: mutually exclusive N = 26 total; item familiarity: N = 9, context familiarity: N = 17), from Experiment 1 ^65,66^. Event-related potentials (ERPs) are shown from representative midline central site Cz; Shaded areas of event-related potentials (ERPs) depict the standard error or the mean for each condition. Bottom: range-normalized topographic maps of individual conditions from 300 to 500 ms, with color scale depicting voltage on the numerical scales indicated. Panel B: Between-subjects effects from participants of Experiment 2 ^1,67^, (N = 24 total; item familiarity: N = 13, context familiarity: N = 11).

Control analyses of range-normalized values revealed that the group differences between the memory conditions were not attributable to scaling artifacts or cohort effects that might be operating independent of the memory conditions. Since it is possible that the results of the between-subjects analyses could be driven in part by group-level differences in overall magnitudes of the ERP amplitudes between the ‘cohorts’ (e.g. scaling artifacts; ^80–83^, we conducted follow-up analyses to control for this possibility. We range-normalized the ERP amplitudes of the individual conditions within each group for item-familiarity and context-familiarity, respectively (Figure 6A, bottom). This approach serves to overcome potential scaling artifacts of differential amplitudes, as it normalizes the ERPs in the dimension of amplitude magnitude and allows for investigation of where they are spatially located ^44,72,83^ as has been done in prior work ^44,66,84^.

Results from this control analysis revealed that range-normalized values for item- and context-familiarity ERPs remained significantly different from 300-500 ms at mid-central site Cz (t(24) = 2.36, p = .027, Cohen’s d = .97, 95% CI [-1.24, -.084]; iFam: N = 9, M = .75, SD = .64, SE = .21; cFam: N = 17, M = .096, SD = .69, SE = .17) (Figure 6A), and thus that the differences between the memory conditions were not attributable to scaling artifacts or cohort effects of groups that might be operating independent of the memory conditions. Collectively, these findings indicated that the observed differences in familiarity response types reflected differential neural processing at the individual subject level of variability, in addition to the trial-wise nature of the memory processing that was identified in the preceding within-subjects analyses. These results thus converge to reveal that the two conditions represent fundamentally different kinds of familiarity operating in service of episodic recognition.

## Results Experiment 2

If there are differential neural correlates of context familiarity and item familiarity operating as distinct processes of episodic memory, then the differences should be evident in other, independent experiments. Therefore, we assessed questions of reproducibility and sought validity of the findings through replication ^85–87^. We investigated if the differences between ERPs for context familiarity and item familiarity (Figures 3, 4, 5, & 6) could be reproduced among independent datasets that were aggregated from three previously-published studies using a nearly-identical paradigm and which had previously reported both traditional ERP correlates of item familiarity as well as ERP effects of context familiarity being separately dissociated from those of recollection, but had not directly contrasted conditions of item-familiarity to context-familiarity ^1,39,67^.

The analyses of Experiment 2 were conducted using the same sites and latencies that had been reported as representative of effects in the present and previous studies (mid-central Cz; bilateral frontal sites F3, F4 ^1,39^). The null hypothesis for these analyses is that ERPs for the two conditions of context- and item-familiarity would exhibit no differences, if in fact the two memory responses were being supported by the same kind of underlying memory processing, and/or if the main results of Experiment were representing a form of type I error. The alternate hypothesis was that context familiarity would exhibit more negative-going ERPs than item-familiarity, and that if there is any such difference in ERP activity for the two conditions then it would be indicative that the two memory response types could not be measuring the same kind of memory process, and thus reflecting different kinds of memory processing people were using to making these different memory judgments ^6,32,40,44–46^.

### Experiment 2: Within-subjects Familiarity

First, we assessed if the within-subjects results observed between item familiarity and context familiarity in Exp. 1 (Figure 5) could be seen at the mid-central site of Cz, using a 2×3 ANOVA with factors of memory (context familiarity, item familiarity) and time (300 - 500 ms, 600 - 900 ms, 800 – 1200 ms; N = 20). This revealed that there was a significant main effect of memory (F(1,19) = 4.55, p = .046, *n^2^_p_* = .193, Cohen’s d = .48), a main effect of time (F(1.26, 19) = 19.72, p < .001, *n^2^_p_*= .509), but no significant memory x time interaction (F(1.68, 19) = .437, p = .616, *n^2^_p_*= .022) (Figure 7). These findings indicated that the two conditions differed reliably at the mid central site throughout the epochs from 300 to 1200 ms, and represented a faithful reproduction of the differences originally observed between ERPs for context- and item-familiarity in the present study’s main findings (Figure 5).

**Figure 7.**
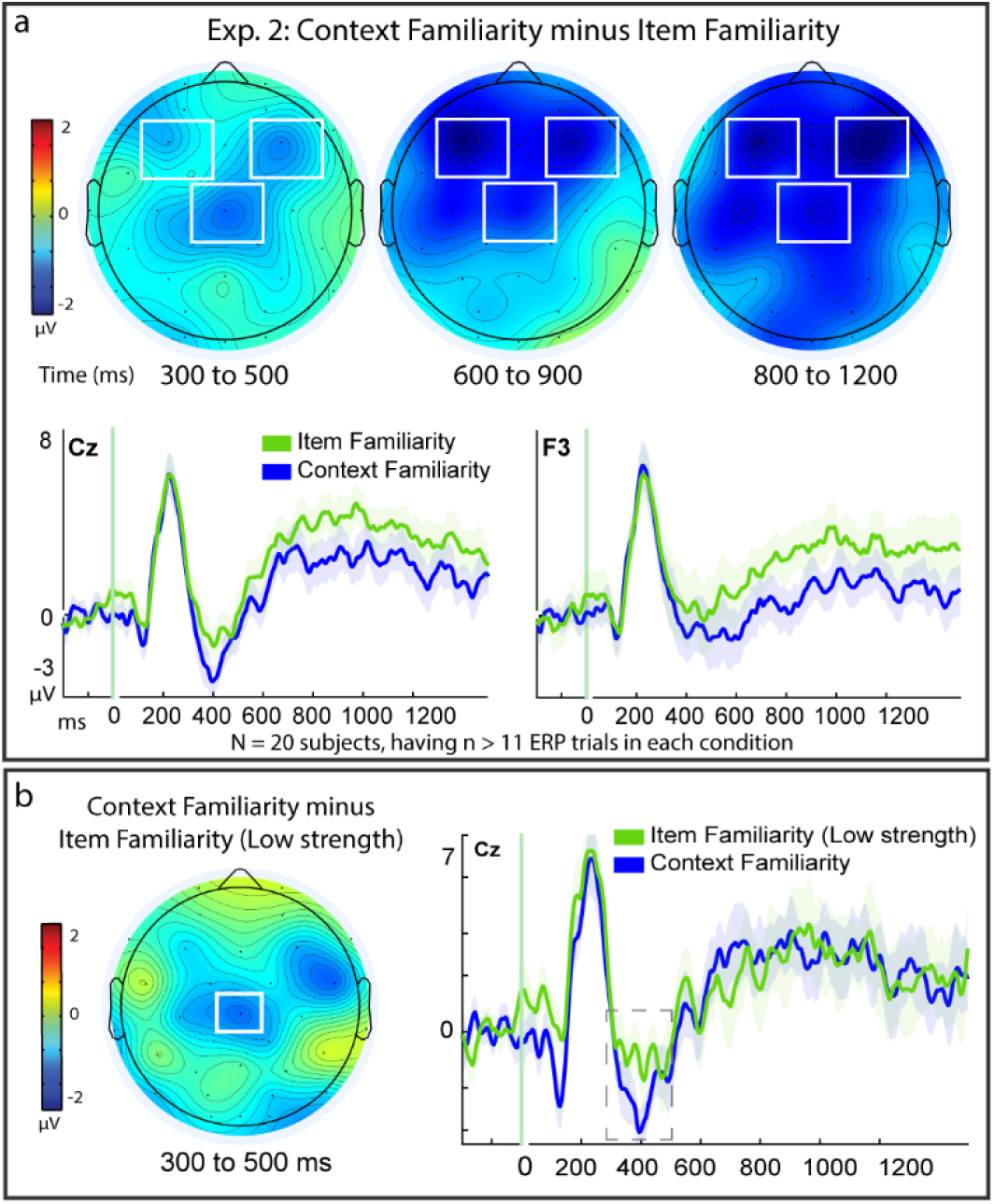
Independent Replication of ERP Differences for Context- and Item-Familiarity. Within-subject effects for participants sharing pairwise response trials in both conditions of item familiarity and context familiarity (N = 20) in Experiment 2 data aggregated from previously published studies ^1,39,67^ (see Method). Panel A, top row: topographic difference maps of the resulting within-subjects difference between context familiarity minus item familiarity, in time intervals shown in milliseconds (ms). Color scale depicts difference amplitudes from -2 to 2 mu-volts, white boxes denote regions of targeted analyses revealing statistically significant effects. ERPs from representative sites Cz (mid-central), F3 (left frontal), and F4 (right frontal); shaded areas depict the standard error or the mean for each condition. Panel B: Same contrast as depicted as in Panel A but excluding high confidence recognition responses from the item familiarity conditions (compare to Figure 5C), to match conditions on item recognition confidence levels (N = 12).

Because the previous datasets ^1,39^ had reported the ERP effects for context familiarity also manifesting at bilateral fronto-central sites, we expanded the aforementioned analyses to also include the frontal sites F3 and F4, using a 2×3×3 ANOVA. This analysis continued to provide a faithful replication of the original findings of the present work, as there was a main effect of memory (F(1,19) = 6.69, p = .018, *n^2^_p_* = .261, Cohen’s d = .49), an effect of time (F(1.29, 19) = 9.69, p = .003, *n^2^_p_* = .338), and a site x time interaction (F(2.24, 19) = 15.20, p <.001, *n^2^_p_*= .444) as the previously-reported frontal sites became more pronounced later in the epoch (Figure 7). Importantly, these results provide an independent replication of the critical findings reported in the present study, comprised of entirely different participants from a different laboratory (the UC Davis Dynamic Memory Lab vs. California State University – San Bernardino, using different EEG hardware. Thus, the results of Experiment 1 in the main text of the present study are bolstered as valid, reliable findings observed across diverse participants from four different independent experiments ^1,39,65,67^.

### Experiment 2: Controlling for confounds of confidence levels in item familiarity

One possible alternative interpretation of the ERP differences we observed between the conditions of item familiarity and context familiarity (Figure 5) is that they could have been confounded by inclusion of high confidence recognition trials in the item familiarity condition. That is, the conditions of item- and context familiarity also vary on the factor of item recognition confidence, since the item familiarity condition includes both high and low confidence responses of strong and weak item familiarity levels, whereas the context familiarity condition contains only the low confidence recognition judgments that had correct source memory. One approach to address this issue is to exclude the high confidence responses from the item familiarity condition^4^, even though, as noted in Experiment 1, the context familiarity condition could still differ from strong forms of item familiarity that were excluded from the analysis ^24,27,28^. This control analysis was performed in the behavioral results reported in Experiment 1 and was found to successfully preserve the findings that the conditions for item- and context-familiarity were reliably distinct in response times.

For the ERP version of this control analysis in Experiment 1, there was only a small sample (N =3) meeting our inclusion criteria for ERPs of pairwise comparisons of the item familiarity condition with context familiarity (Figure 5C) that still provided a significant effect, but would benefit from an independent replication in a larger sample. In Experiment 2 there were more participants available (N = 12). We thus assessed ERPs for ‘item4+source unknown’ (iFam_Low_.) vs. ‘item4+source correct’ (cFam.), using a directional t-test based upon the a priori hypothesis derived from the prior findings in Experiment 1 that context familiarity exhibits more negative-going ERPs than item familiarity at the same mid-central size (Cz) and time (300 to 500 ms) as was found in Experiment 1. This revealed that there were still significant effects (t(11) = 1.97, p = .037, one-tailed, Cohen’s d = .60, 95% CI [-∞, -.101]; iFam: M = -.849, SD = 2.74, SE = .79; cFam: M = -1.99, SD = 2.06, SE = .593), and thus the main finding was replicated again and with a larger sample than original finding in Experiment 1. Thus, the inclusion of the high confidence recognition responses in the condition of item familiarity is not contaminating nor confounding the results we observed of reliable differences between item- and context-familiarity (Figure 5A, Figure 7A).

### Experiment 2: Between Subjects Familiarity

The between-subjects ERP effects from Exp. 1 (Figure 6A) were also reproduced when aggregating across participants from prior studies ^1,67^ (Figure 6B). As was done in the main results, participants were included in conditions of either item familiarity or context familiarity if they met the trial inclusion criterion of sufficient trial responses (n = 12) in exclusively one condition but not the other, and these were entirely different participants than those included in the within-subjects analysis described above. This resulted in a subgroup of N = 13 for the item familiarity condition, and N = 11 for context familiarity. As familiarity effects for these prior studies were reported for 400 to 600 ms ^1,39^, a targeted analysis was conducted during this latency to assess if context familiarity ERPs exhibited the same negative-going difference from item-familiarity ERPS at the mid-frontal site characteristic of previous familiarity effects reported in these participants ^1,67^ (Fz). A between-group t-test revealed the ERPs to reliably differ (t(22) = 1.923, p = .034, one-tailed, Cohen’s d = .79, 95% CI [-∞, -.426]), with ERPs for context familiarity again being more negative-going than those for item-familiarity (N = 11, M= -2.69, SD = 4.98, SE = 1.50; N = 13, M = 1.25, SD = 5.00, SE = 1.39, respectively).

### Experiment 2: Within-subjects Memory Condition Effects

Next, the results of each individual condition’s memory effect were compared to correct rejections, as was done in Figure 4 of the main findings of Exp. 1. ERP correlates of context familiarity again exhibited a reliable effect from 800 – 1200 ms (N = 33) that was distributed broadly across the central-posterior scalp and maximal at site Cp1 (t(32) = -3.22, p = .003, Cohen’s d = .56, 95% CI [-3.17, 7.12]; cFam: M = 1.84, SD = 4.35, SE = 0.76; CR: M = 3.78, SD = 3.94, SE = 0.69), but for which there was no significant FN400 effects evident from 400-600 ms (t(32) = .163, p = .113, 95% CI [-.195, 1.75], Bayes Factor_01_ = 1.25; cFam: M = -.006, SD = 4.24, SE = .74; CR: M = -.782, SD = 3.84, SE = .67), nor any evident LPC at P3 from 600-900 ms (t(32) = 1.07, p = .293, 95% CI [-1.66, .515], Bayes Factor_01_ = 3.17); cFam: M = 2.19, SD = 4.42, SE = 0.77; CR: M = 2.76, SD = 3.66, SE = 0.64) (Figure 8). This finding reproduced and confirmed the Exp. 1 finding (Figure 4) that context familiarity was associated with a broad central negative effect, that it held no evidence of association with the FN400 and provided an evidence for absence of the LPC effect.

**Figure 8.**
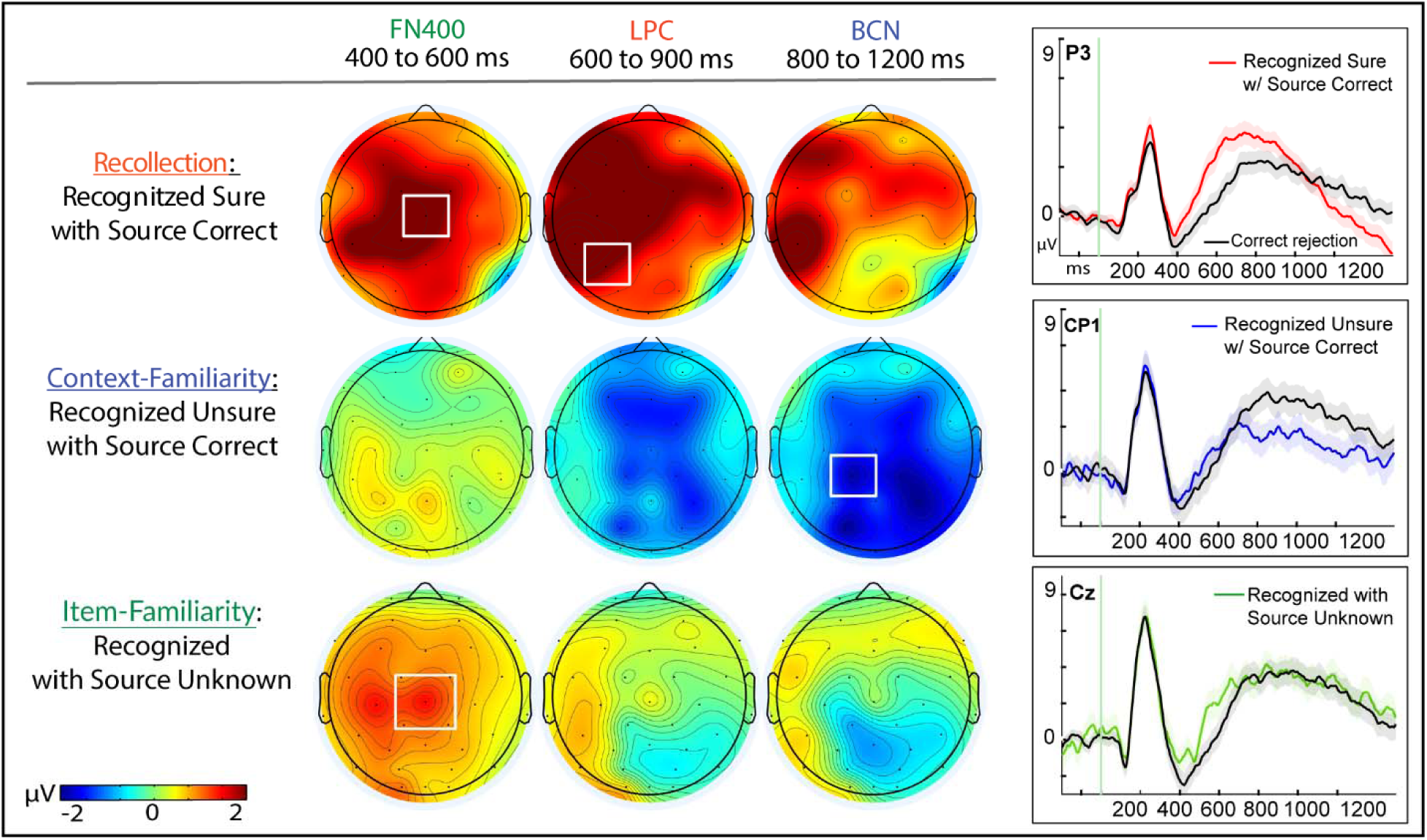
Physiological effects of episodic memory retrieval conditions. Left panels: Topography maps are shown for each memory condition as compared to correct rejections in Experiment 2, comprising data from three previously published studies utilizing the same paradigm as the present study ^1,39,67^. (top) High-confident item recognition judgments that also had correct source memory retrieval; (middle) low-confident item recognition judgments of that also had correct source memory retrieval; and (bottom) recognition hits (collapsed across both high and low confidence) that did not have source memory retrieved (received responses of ‘source unknown’). Color scale of topographic maps denotes amplitude differences for each condition as compared to correct rejections, ranging from -2 to 2 mu-volts. Right panels: Event-related potential (ERP) amplitudes (µV) from representative sites of maximal activation for each of the respective effects during the memory retrieval epoch of -200 to 1200 milliseconds (ms); shaded areas for each plot represent the standard error of the mean.

On the other hand, item familiarity was associated with significantly more positive ERPs than correct rejections at mid-central Cz from 400 to 600 ms (N = 38) (Figure 8), reproducing the FN400 effect reported in Figure 4 of the main results section (t(37) = 3.77, p < .001, Cohen’s d = .61, 95% CI [-2.38, -.715]; iFam: M = .362, SD = 3.78, SE = .61; CR: M = -1.18, SD = 3.79, SE = .62). During the later latencies of 600 to 900 ms and then 800 to 1200 ms associated with recollection and context familiarity, respectively, there were no reliable differences evident between the two conditions (t(37) = 0.95, p = .348, 95% CI [-1.44, .519], Bayes Factor_01_ = 3.76; iFam: M = 3.27, SD = 4.47, SE = .725; CR: M = 2.81, SD = 4.34, SE = .703); t(37) = .160, p = .877, 95% CI [-1.08, .923], Bayes Factor_01_ = 5.66; iFam: M = 3.49, SD = 4.39, SE = .71; CR: M = 3.41, SD = 4.19, SE = .68) (Figure 8), reproducing the same pattern of results reported in the present study’s main results (Figure 4). These findings indicated that the neurophysiological correlates of the item familiarity condition were linked with a FN400 effect but no comparable evidence for any LPC or BCN effects (and with Bayes Factor evidence of absence for each other effect, respectively), consistent with the well-established profile of item-familiarity ERPs observed within and across the two present Experiments and among the broader literature of other studies ^1,43–45,57,84^.

Furthermore, similar to the findings of the present study, the recollection response condition exhibited both a reliable FN400 and LPC effect when compared to correct rejections (N = 54), (t(53) = 6.14, p < .001, Cohen’s d = .84, 95% CI [1.43, 2.81]; Rec.: M = .207, SD = 4.92, SE = .668; CR: M = -1.915, SD = 4.53, SE = .61; t(53) = 4.73, p < .001, Cohen’s d = .64, 95% CI [.942, 2.33]; Rec. M = 3.92, SD = 3.96, SE = .54), but provided no evidence for any BCN (t(53) = .585, p = .561, 95% CI [-1.55, .848], Bayes Factor_01_ of 5.72) (Figure 8, 9). Collectively, these analyses of previously published datasets using the same memory paradigm as the present study provide a replication of the results observed in the current investigation. These patterns of effects reported here across both behavior and neurophysiology, both within and between subjects, and their replications among several other independent studies cannot be accounted for by a priori predictions from existing models of only a single- or dual processes of episodic memory.

**Figure 9.**
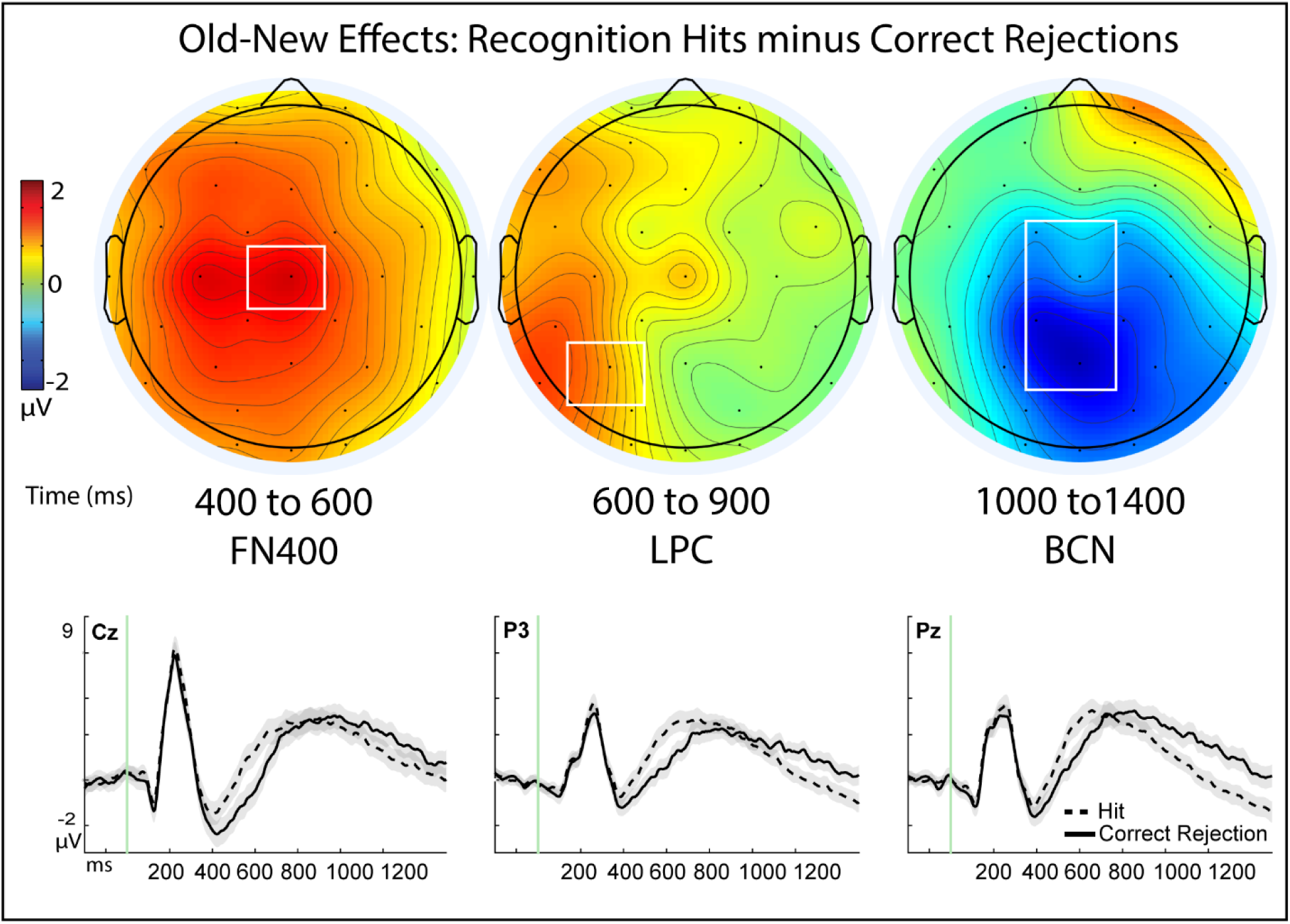
Old-New Effects for Recognition Memory in Experiment 2. Data for Experiment 2 was aggregated from three previously published studies ^1,39,67^ (N = 56). Top row: topographic difference maps of the resulting difference between hits minus correct rejections, plotted for each of the three time latencies of 400 to 600 milliseconds (ms), 600 to 900 ms, and 1000 to 1400 ms that have been associated with the FN400, late parietal component (LPC), and late broad central negativity (BCN), respectively. Color scale depicting difference amplitudes from -2 to 2 mu-volts, white boxes denote regions of targeted analyses revealing statistically significant effects. Bottom panel: Event-related potential (ERP) amplitudes (µV) of individual conditions of recognition hits (dashed line) and correct rejections (solid line) plotted through the 1500 ms epoch and beginning with a -200 ms baseline period. Shaded areas depict the standard error or the mean for each condition.

### Experiment 2: Within-subjects Old-New Memory Effects

Using the same aggregated data also reproduced the other findings reported in the present study of a triple dissociation of ERP effects for item-familiarity, recollection, and context familiarity among the general memory contrast of hits minus corrections (Figure 3), whereby a 3-way interaction was originally found among an FN400, LPC, and BCN. For this analysis, we focused upon the latencies where effects had been observed in the prior studies for the FN400, LPC, and BCN (400 to 600 ms, 600 to 900 ms, 1000 to 1400 ms, respectively^5^), and focusing on sites Cz, P3, and Fc1 reported for effects in the present study (N=56). Results from a 2×3×3 ANOVA revealed that, similar to the main findings from Experiment 1 of the present study, there was no initial main effect of memory (F(1,55) = 2.19, p = .145, *n^2^_p_*= .038), but there were significant main effects of site (F(1.68, 55) = 5.82, p = .007, *n^2^_p_* = .096), and significant interactions of memory x time (F(1.52, 55) = 32.77, p < .001, *n^2^_p_* = .373), site x time (F(1.57, 55) = 23.59, p < .001, *n^2^_p_*= .300), plus a significant memory x site x time interaction (F(3.20, 55) = 4.77, p = .003, *n*^2^ = .080) (Figure 9).

Consistent with the a priori hypotheses, results from directional t-tests based upon our a priori hypotheses revealed that there were significant effects of the FN400 (t(55) = 6.16, p < .001, one-tailed, Cohen’s d = .82, 95% CI [1.27, inf]; Hit: M = -.208, SD = 4.39, SE = .588; CR: M = -1.95, SD = 4.48, SE = .599), LPC (t(55) = 3.06, p = .002, one-tailed, Cohen’s d = .41, 95% CI [.402, inf]; Hit: M = 3.13, SD = 3.63, SE = .49; CR: M = 2.24, SD = 3.99, SE = .534), and BCN (t(55) = 1.85, p = .035, one-tailed, Cohen’s d = .25, 95% CI [-∞, -.083]; Hit: M = .916, SD = 3.68, SE = .49; CR: M = 1.80, SD = 4.60, SE = .62). The BCN effect identified at the a priori selected location of Fc1 was observed to have extended broadly and been maximal at site Pz: t(55) = 4.33, p < .001, one-tailed, Cohens d = .58, 95% CI [-inf, -1.14], Together, these results revealed the same finding of a significant memory x site x time interaction in an independent data set, effectively reproducing the original findings reported in the present study (Figure 3, Figure 9). Specifically, these results confirm that the three different memory effects are present within the most general measures of memory historically used in the field, in addition to the specific conditions described herein.

### Are the neural correlates of context familiarity reflecting non-conscious memory processing**?**

While the responses given for context familiarity were defined as being explicit declarative responses from people for item recognition hits and also had overtly-declarative correct source memory judgments, it nevertheless remained theoretically possible that they reflected a form of non-conscious processing such as that related to guessing from 100-300ms ^88^ or fluency that has been known to occur characteristically in earlier latencies of 200-400ms ^84,89,90^ for Review see ^75^. Thus, to rule this possibility out, we sought to verify if the memory processing associated with the context familiarity condition’s effects in these participants was different from the hallmark signs of non-conscious, implicit memory signals that have been extensively characterized in the prior literature (i.e.: recognition misses ^44,66,75,84^, old items endorsed as 1 or 2 responses during the memory retrieval test). The null hypothesis of this analysis was that if the context familiarity condition was actually representing an implicit memory process, then it should show the same, or similar, activity as other implicit memory processing, such as the misses in the time windows when each effect is known to operate, respectively (200-400 ms for implicit guessing, 800-1200 ms for context familiarity), and thus predict no significant differences in ERPs.

### Experiment 1: Misses

We assessed ERPs for misses and context familiarity at the centro-parietal site of peak activity identified earlier for context familiarity (Cp1), using a within-subjects two-factor ANOVA with repeated measures on both factors (N = 27). This analysis revealed no significant main effect of condition (F(1,26) = 1.03, p = .321, *n^2^_p_* = .038), identified a significant main effect of latency (F(1.91, 26) = 19.95, p < .001, *n*^2^ = .434), and importantly revealed a significant interaction of memory condition and latency (F(2.16, 26) = 3.78, p = .026, *n^2^_p_* = .127), which was further investigated with follow-up analyses (Figure 10A). This revealed that ERPs for context familiarity were significantly more negative-going than misses at centro-parietal sites (Cp1) from 800 – 1200ms (t(26) = 2.31, p_holm_ = .029, corrected, Cohen’s d = .44, 95% CI -2.40, -.140]; Context Familiarity M = -2.69, SD= 3.46, SE = .66; Miss M = .-1.43, SD = 2.34, SE = .45) though not in the earlier latency of 200-400ms (t(26) = .443, p = .660; Bayes Factor_01_ = 4.37 indicating moderate evidence for the null). This demonstrated that the context familiarity condition was exhibiting substantively different physiology from recognition misses at centro-parietal sites during 800 to 1200 ms, refuting the null hypothesis that they were both reflecting the same implicit process outside of conscious awareness (e.g. guessing driven by implicit factors).

**Figure 10.**
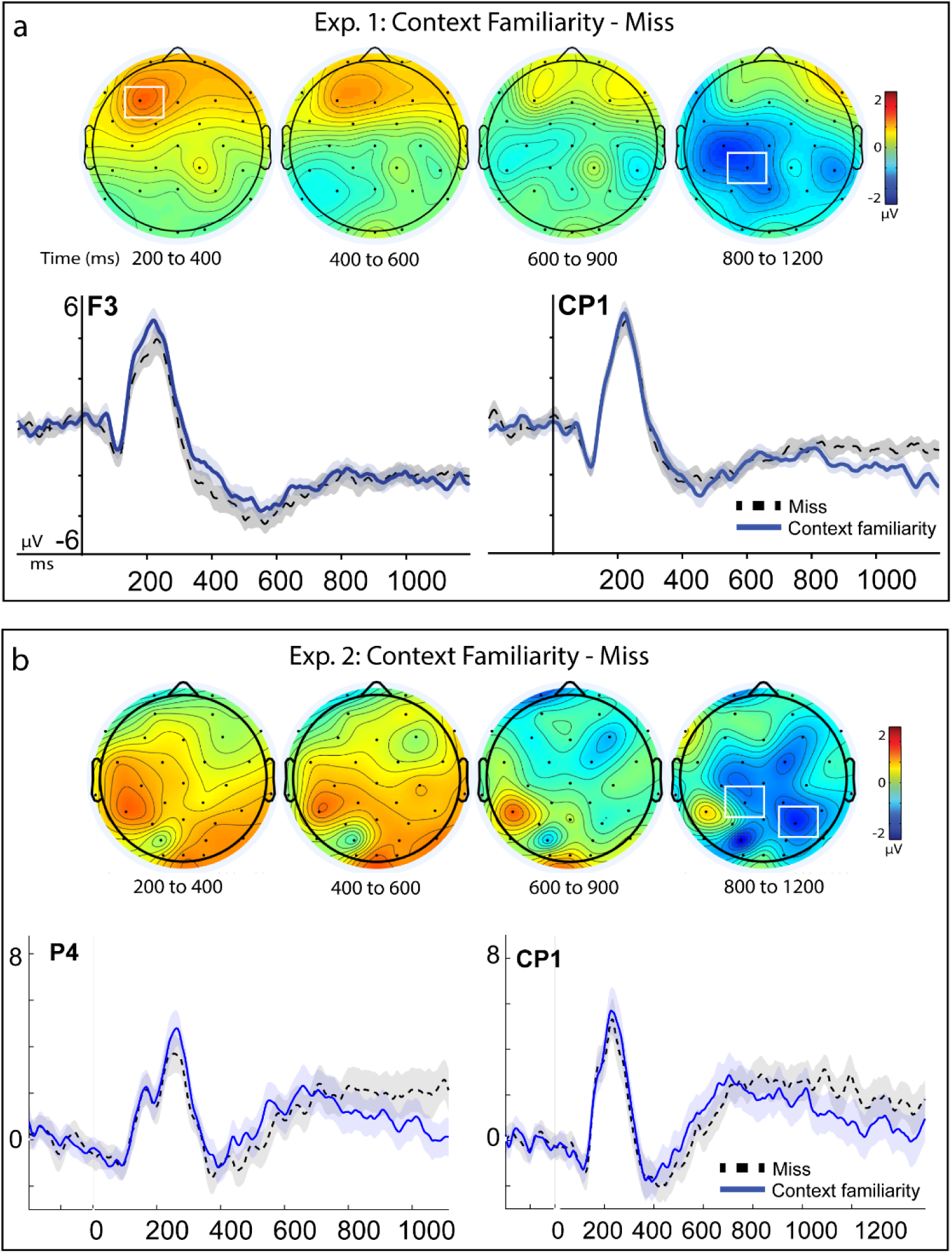
Physiological effects of context familiarity versus recognition misses. Panel A: Top: topographic maps of the difference map for activity of context familiarity minus activity for recognition misses (N = 27). Maps are plotted for each time interval of 200-400 milliseconds (ms), 400 to 600, 600 to 900 ms, and 800 to 1200 ms; color scale of topographic maps denotes amplitude differences from -2 to 2 mu-volts. Event-related potential (ERP) amplitudes (µV) for each condition plotted for representative sites of maximal activation (left frontal, F3, and centro-parietal Cp1), through an epoch of -200 to 1200 ms of a memory retrieval test. Shaded areas depict the standard error or the mean for each condition. Panel B: Replication data from Experiment 2. Data from three previously published studies (N = 27). ERPs for each condition plotted for representative sites from original analysis (Cp1) and of maximal activation (right parietal Cp5, 200-400 ms), through an epoch of -200 to 1200 ms of a memory retrieval test.

Since topography maps of Experiment 1 also indicated an early frontal activity pattern for this contrast, we conducted an exploratory analysis of the evident frontal region at its maximal left frontal site of F3. Inspection of these findings revealed that ERPs for context familiarity were more positive-going than misses in the early latency of 200-400ms (F3: t(26) = 2.55, p = .034_holm_, corrected, Cohens d’ = .49, 95% CI [-2.07, -.221]; cFam: M = 1.57, SD = 2.89, SE = .55; Miss: M = .427, SD = 3.01, SE = .57), continuing marginally 400-600 ms (t(26) = 1.90, p_holm_ = .068, corrected, Cohen’s d = .37, 95% CI [-1.85, .07]; cFam: M = -4.36, SD = 3.62, SE = .697; Miss: M = -5.26, SD = 4.11, SE = .792) (Figure 10A). Overall, these analyses revealed reliable, significant differences between conditions of context familiarity and implicit memory misses in Experiment 1 but were also assessed for the replicability in Experiment 2.

### Experiment 2: Misses

The same pattern of differences between context familiarity and misses was also observed as being independently reproduced by targeted analyses in Experiment 2’s data sets (N = 27) based upon directional hypotheses derived from the preceding findings for the same contrasts in Exp. 1, during the 800 to 1200 ms time interval of context familiarity effects, occurring at the same site as the primary results (Cp1: (t(26) = 1.79, p = .043, one-tailed, Cohen’s d = .34, 95% CI [.046, ∞], cFam: M = 1.68, SD = 3.68, SE = .707; Miss: M = 2.68, SD = 3.35, SE = .644; maximal activation site of P4 (t(26) = 2.35, p = .013, one-tailed, Cohen’s d = .45, 95% CI [.396, ∞], cFam: M = .689, SD = 3.33, SE = .64; Miss : M = 2.14, SD = 3.34, SE = .681) (Figure 10B). Similar results to the primary findings in the earlier latencies of 200-400ms and 400-600 ms were also observed in Experiment 2’s datasets, revealing more positive ERP activity for context familiarity, albeit in more central-parietal sites of these participants (maximal at Cp5) (t(26) = 2.62, p = .007, one-tailed, Cohen’s d = .50, 95% CI [∞, -.400], cFam: M = 1.93, SD = 2.71, SE = .52; Miss: M = .786, SD = 2.84, SE = .55; t(26) = 1.81, p = .041, one-tailed, Cohen’s d = .35, 95% CI [∞, -.062], cFam: M = .019, SD = 3.08, SE = .594; Miss: M = -1.08, SD = 3.48, SE = .67, respectively).

Together, these results from both Experiments 1 & 2 revealed that neural activity for explicitly declared context familiarity judgments was reliably differentiated from that of implicit misses at the human scalp, during early latencies (200-400 ms) and later latencies (800 – 1200 ms), and across several different independent studies. This line of converging evidence refuted the null hypothesis that both conditions of context familiarity and implicit misses had the same functional significance or were representing the same cognitive process ^44,69–73,91^. More broadly, recent work in the same experimental paradigm has also identified that other, more pure, kinds of implicit recognition processes can operate distinctly from misses^66^, but that pattern of implicit processing (positive-going right parietal activity from 600-1000 ms) was entirely opposite of the negative-going central site patterns of ERPs characterized here for context familiarity, and thus not possible as a viable account of the present findings either. In sum, explicit context familiarity judgments were supposed to be equivalent to misses if representing similar implicit processing, but instead were significantly more positive than implicit misses in early epochs at frontal sites and then reversed to have significantly more negative voltage than at centro-parietal sites at later times in the epoch, thus providing clear evidence of divergent validity from the measures of implicit memory processing. This pattern for context familiarity cannot easily be attributable to accounts such as guessing or signals outside of episodic awareness and lending further weight against the possibility that context familiarity judgments were driven by implicit processes.

## Discussion

### General Summary of Findings

We began the investigation to understand how (by which process?) people make accurate source judgments for items they recognize with low confidence. Emphasizing reproducibility in science ^85–87,92–96^ the current study first replicated the original behavioral performance and ERP effects of context familiarity as previously reported ^1^. Extending those findings, the main finding here was that the effects of item familiarity, context familiarity, and recollection were reliably dissociable in multiple measures of both behavior and neural activity; these ERP effects were then reproduced in independent datasets. These patterns of effects cannot be accounted for by a priori predictions made by models of only a single- or dual processes of episodic memory without making post-hoc modifications. Item- and context-familiarity thus appear to be operating as fundamentally different kinds of memory processes, based upon established norms for identifying cognitive processes via neuroimaging and behavioral data ^40,44,70–73,97–101^, such that dissociable physiological activity occurring at different times, at different locations, and for different conditions, cannot be taken as evidence that they are reflecting the same cognitive process and must therefore reflect at least partially non-overlapping cognitive operations. Together, this provides strong evidence that these three observations of ERP effects of memory are reflecting mutually exclusive, distinct episodic memory processes, and therefore advancing an updated model of the organization of episodic memory as discussed in further detail across each finding throughout the sections below.

### Behavioral findings

Item familiarity and context familiarity were found to be behaviorally distinct in measures of response time, both in item judgments and in source judgments. Control analyses that held memory strength constant among conditions still preserved these differences and demonstrated that they were not confounded by recognition confidence levels. Item familiarity responses were faster during item judgments, whereas context familiarity responses were faster during source memory. It is difficult to account for these patterns of dissociations if attributing the responses to the same underlying cognitive processes of a single familiarity signal ^20,28,29,32^. Context familiarity responses were also responded to significantly different than were responses of recollection, with recollection being faster than both item- and context-familiarity conditions during recognition and slower during source memory (Figure 2). There are two different ways that one can interpret these patterns of behavioral results: (1) They are consistent with existing findings of the relative speeds of recollection and familiarity (recollection being slower and familiarity being faster ^32,53^; (2) they are contrary to traditionally-held patterns of recollection being slower than familiarity. Each view will be discussed in more detail below, but importantly, regardless of interpretation, the memory conditions were reliably distinct in behavioral measures, indicating that mnemonic processing for context familiarity is distinct from item familiarity and recollection.

Traditionally, recollection has been conceptualized as a slower, deliberative process, whereas familiarity was modeled as a faster, automatic process ^32^. As reviewed in the Introduction, source memory judgments are typically considered a general (though not exclusive) measure of recollection, which in the present study was reliably slower during source memory than the conditions of both context- and item-familiarity responses. Those results from source memory RTs are thus consistent with traditional characterizations of recollection as a slower, deliberative process ^32^. Accordingly, item recognition tasks are usually considered to be a relatively better measure of familiarity-based responding, and faster response times observed here for item-familiarity than for context-familiarity in the present findings can thus be seen as it being the item (not the context) that is familiar during this early stage of the memory task. In this view, that recollection responses were also faster in item recognition would be seen merely that their inclusion of the highest confidence responses was driving a concurrent higher item familiarity and thus faster recognition time, consistent with traditional patterns. As the context familiarity responses then go on to become faster than item familiarity responses during the source judgments, it preserves the traditional notion that familiarity is also a faster automatic process-this time it is familiarity for context that is differentially faster (not items). That is, in a traditional view of RTs results are seen as showing that the two different types of familiarity (item, context) are faster than each other in each of the respective domains in which they are tested for representing differentially (item recognition, source memory), and ultimately consistent with the common findings that familiarity operates faster, while recollection is a slower process.

Alternatively, the pattern of faster recollection during item recognition responses (and slower than context familiarity in the source judgments) could also be seen as departing from common findings that recollection is generally a slower process than item familiarity. While some studies have found that recollection can sometimes be faster than familiarity in some instances ^102,103^, that discrepancy was suggested to likely be due to instructions to participants that they respond after both memory processes are complete-thereby delaying the responses times ^32^. The current paradigm also requires subjects respond to a combination of both item and source memory confidence ratings on each trial, which could similarly impose complex demands on their memory judgements. Moreover, recent studies noted below have reported recollection to have faster responses than familiarity during recognition, suggesting that the traditionally assumed speeds of recollection and familiarity should not necessarily always be taken as simply 1-directional (i.e.: that familiarity must always be faster than recollection) and that is likely more complex in directionality.

For instance, a review of pupillometry studies of recognition memory concluded that, contrary to common observation of behavioral studies, recollection is not always the characteristic ‘slow and effortful’ process typically assumed, but can also be quite ‘fast and easy’ in its manifestation as a memory process ^104^. When more advanced modeling methods were used for measuring the speeds of memory processes across six different experiments, recollection was again found to operate as both fast and slow ^105^, similar to findings in the present data. Such patterns are also consistent with proposals that recollection is a two-stage process that includes both an initial rapid assessment of memory, plus a later second stage that is slower and more deliberative ^106^, and with behavioral studies suggesting a third recognition process of familiarity ^107^. Together, the response time profiles observed in the present work (Figure 2) are consistent with other studies ^108^, while also adding insight from specific memory response conditions that have not previously measured response times, to reveal that regardless of chosen interpretation views, the conditions remain nevertheless as neurocognitively distinct.

### Physiological findings

In physiological measures, first, a triple dissociation of activity patterns was evident in general ERP effects observed during the basic memory measure of hits vs. correct rejections, qualified by a significant condition x time x location interaction that revealed a mid-central FN400 effect from 400-600 ms, a left parietal LPC Effect from 600-900 ms, and a broad central negativity effect (BCN) from 800-1200 ms, and replicated in a second independent data set. Second, when investigating individual ERP effects of the specific memory conditions of item-familiarity and context-familiarity, each was found to be reliably different within-subjects and across subjects and was dissociated across the different latencies and locations on the scalp as noted above, and each was also dissociated from recollection. The same results were observed for each independent condition when contrasted against the standard ERP baseline condition of correct rejections; all of these findings were also reproduced in archival data from independent studies. This suggests that differential neurocognitive processes underlie each of the memory judgments, respectively ^44,70–73,101^.

The neural correlates for context familiarity did not exhibit the LPC nor the FN400 effects (the putative correlates of recollection and item-familiarity, respectively) in any analysis. The absence of such standard ERP effects was further bolstered by the Bayes Factor results, which provided moderate evidence that both the LPC and FN400 were not present for these memory judgments instead of being merely undetectable. This conclusion is further bolstered by the finding that context familiarity exhibited significant effects present elsewhere in time and scalp location, so the absence of LPC and FN400 correlates could not be merely a case of insufficient statistical power to detect such differences.

### Why might context familiarity effects occur later than item familiarity?

In our main ERP contrasts of each memory condition (vs. correct rejections, Figure 4, Supplementary Figure 1), context familiarity manifested effects later than item familiarity (800-1200 ms vs. 400-600 ms, respectively), and behavioral response times for context familiarity judgments were similarly later than those for item familiarity (2710 ms, 2361 ms, Figure 2, Table 2) [though we note that during the source memory judgment context familiarity responses were faster than item familiarity (2075 ms, 2612 ms, respectively)]. To a certain extent, the present work is not designed to directly answer *why* the times differ, but importantly is reporting the original finding that *they are, in fact, different at all*. One speculative reason is that context information may be broader than item information: encompassing additional elements/features that require neural pattern completion ^109–113^– and that this could take the extra few hundred milliseconds that we observe here. We note here that the later ERP time interval (800-1200 ms) was not the only time that context familiarity was found to exert an effect: it also differed reliably from item familiarity ERPs during the early epoch of 300-500 ms (Figures 5, 6, 7), too. We underscore that this early finding was important to the present work, because it demonstrated that the two memory conditions are occurring through differential patterns of neural activity early in the epoch (300-500 ms)-not just later.

There should also be caution in over-interpreting the response time patterns of ERP effects earlier than the behavioral effects. The timings of ERP effects are occurring substantially before the later behavioral memory judgments, which might seem odd to researchers unfamiliar with ERP research on memory. However, this pattern of faster ERP times than behavioral times for the same memory processes has been generally true for most studies throughout the literature. This is because ERP effects have generally never been thought to represent a direct 1:1 mapping of memory process (as discussed in ^39^) but instead are widely regarded as “the putative neural correlates” of the memory processes ^45^. That is, the ERPs are *covert* measures of memory-related activity^79^, but the *overt* memory behavior is generally thought to occur later in time (when people indicate their memory response) after the initial neural processing has set the stage for integrating the ensuing cognition and then deploying the actual motor response. So, there should be caution exerted in assuming too much about the timing of ERPs-what matters most, in our view here, is that they differ at all, and thus must be representing at least partially non-overlapping cognitive processing ^44,45,71–73,101^.

### Possible Alternative Interpretations

How are study participants making accurate source memory retrieval for items recognized with low confidence? There are several different accounts which could potentially be used to interpret them for future explorations. We will address four possible alternatives and limitations here for what other accounts might explain the pattern of findings for context familiarity: a) generic item familiarity, b) delayed recollection, c) the late parietal negativity (LPN) observed in other studies for controlled search processes, and d) implicit memory or guessing. Each is reviewed below.

### A single signal of generic familiarity?

One possibility to account for the present findings is that results for context familiarity could perhaps ‘simply’ reflect a variant of generic familiarity-based processing for which item and context are merely differences in levels but not kinds, such as the distinction between absolute (pre-experimental) and relative familiarity (in-the-experiment) of the stimuli ^74,75^. This potential account would propose that the ERP measures of each condition of item- and context-familiarity would vary incrementally rather than fundamentally, similar to how ERP measures have been found to vary incrementally to incremental gradations of item familiarity strength ^1,114,115^. Yet, that pattern was not found in the present study, which instead revealed the two conditions to vary in fundamental, not incremental, ways. The present effects of context familiarity were significantly dissociable from the effects of item familiarity-in measures of both behavior and physiology, so cannot be the same ^44,45,70–73^. Thus, a shared, single generic process of familiarity cannot account for this pattern of memory responses that have been observed here and in several independent data sets.

The ERPs for conditions of item- and context-familiarity were found to significantly differ in N400 potentials from 300-500 ms, and the N400 component has also been known to be influenced by factors such as context ^91,116–121^. This association with context is consistent with the present interpretations of the psychological constructs we describe as ‘context familiarity’ and was differentiated from the opposing patterns observed for item-familiarity. The posterior distribution of the FN400 effect for the item-familiarity condition is consistent with other findings ascribing it to a sub-type of absolute familiarity (as opposed to relative familiarity, which is reported to have a more anterior scalp distribution ^75,91,122,123^. The FN400 effect observed in the present study had the same mid-central topography as was reported originally for this paradigm ^1^, which is between the frontal and posterior distributions observed in other studies that manipulated factors of relative and absolute familiarity ^74,75,118–120,122,123^, respectively. This mid-central distribution may likely reflect a combination of both relative and absolute familiarity with the stimuli in this particular paradigm since the study was not designed to differentiate those factors.

Finally, accurate source memory has also been shown to be sometimes achieved through reliance on cognitive processes such as item familiarity ^50,56^ and unitization ^58–60,62,63,124,125^, and the Source Monitoring Framework ^53,54^ has directly proposed that familiarity can support source judgments in some contexts. However, weighing against such possibilities of the source judgments being contributed to by those forms of unitized item-familiarity is that prior ERP findings for item familiarity and unitization contributing to source judgments are found as positive-going ERP difference effects ^50,75,124^, instead of the significant negative-going ERP effects observed in the current data.

### Does the BCN reflect a delayed recollection process?

Another possible account for the effects of the context familiarity condition is that perhaps recollection is a bit delayed and happening later in the epoch^6^. There are several broad reasons why it is likely that the ERP effects of the context familiarity condition could not plausibly be reflecting a “Delayed recollection” process, which we will follow with several additional specific reasons. For instance, first there is scant evidence to support a contribution of recollection to the condition we measured as context familiarity: there has been no ERP evidence of the putative neural correlates of recollection for this condition across several different studies, and this absence of evidence has been quantified by Bayes Factor analyses providing consistent evidence of absence. Alternatively, the evidence that has accrued across various studies and measures continues to unambiguously indicate that the condition of context familiarity differs from recollection trials in substantive ways that include behavioral response times for both recognition and for source memory judgements, and several physiological measures across several experiments.

The context familiarity condition also tends to be low confidence source judgments, which argues against the premise that a later process of slow recollection might be driving their trials following a delayed memory search. This is because if recollection occurred later then it could be assumed to have more high confidence correct source judgments (and yet the response data doesn’t have that profile). That is, if recollection is “delayed”, then the common features of recollection should also be evident in at least a reasonable proportion of those source memory response conditions for when that response is ‘eventually’ happening, and those features of recollection commonly include high confidence responses for source memory ^29–32,52,114,126–128^. Since that, too, runs contrary to the pattern of responses evident in the present data, it weighs against the theory of ‘delayed recollection’. This alternative account of delayed recollection would also de facto hold that a delayed recollection arrives later in time, even though the source judgments for this particular condition are provided faster than the source judgments in the actual recollection condition. As such, it is difficult for that to be reconciled in light of the pattern of behavioral responses. The ‘delayed recollection’ account also would have trouble explaining why the context familiarity ERPs differed reliably from the item familiarity ERPs during the early latency of 300-500 ms if it was only representing a delayed process happening later after 1200 ms. For these reasons, it is improbable (and unsupported by evidence) to speculate that the effects of context familiarity could plausibly be reflecting a delayed recollection; however, more specific reasons to reject this theory also exist and are detailed below.

The term of ‘delayed recollection’ appears only sparsely in the memory literature. One of the few extant studies of “delayed recollection” using ERPs ^129^ found that ‘delayed recollection’ occurred under conditions of “changed viewing conditions”-which were not a part of, nor directly applicable, to the present study. Notably, that study did not find any negative-going ERP effects to be associated with their condition of ‘delayed recollection’, which was instead associated with the traditional, positive LPC measures (the opposite of the present findings). Since recollection typically produces a specific hallmark pattern of positive ERP effects ^42,43,45,46^, and we instead observed a distinct opposite polarity of negative-going ERP effects, the logic to support a ‘delayed recollection’ account remains lacking. That is, there is not a reasonable explanation for why the polarity of the canonical recollection-related physiology would have been reversed among our other control analyses demonstrating the standard positive-polarity of recollection effects in all other standard conditions of memory. This provides an additional line of evidence against a ‘delayed recollection’ account and it is thus difficult to support as a viable account of our present findings.

Another related ERP study ^130^ claiming ‘delayed recollection’ did report a later negative-going difference for hits vs correct rejections that they attributed as late parietal negativity (LPN, see section 5.4.3 below), but in their paradigm their reported LPN effect actually became more reduced in their ‘delayed memory condition’, and was instead larger in their condition of immediate memory retrieval. This, too, is the opposite of what would be predicted from a ‘delayed recollection’ account of the present findings, and thus it, too, ends up providing evidence against the ‘delayed recollection’ account. Could the BCN effects we attribute to context familiarity instead be reflecting processes of non-criterial recollection ^131^? If so, one could predict that then there would still be an LPC associated with recollecting the non-criterial information-a pattern of results that we did not see in the present series of studies, and for which there was Bayes Factor evidence of absence.

The alternative theory of ‘delayed recollection’ is further complicated by other problematic aspects: it would predict no differences in ERPs between the condition we call context familiarity and that of correct rejections because if there wasn’t memory/processing yet (i.e. it occurred via a ’delayed recollection’) then there would be no reason for the condition to differ from a non-memory condition baseline of correct rejections. But that is not what was found in the data: there is in fact a significant difference between the condition of context familiarity and correct rejections, and we need to then know why- or, at least, be able to offer a plausible account to explain why that difference exists. Our account provides at least one viable and parsimonious explanation (it is the dissociable memory process of context familiarity), whereas the delayed recollection account does not provide an explanation.

The theory of delayed recollection is additionally difficult to sustain when one thinks about it more deeply to its natural consequent end. It is, in effect, saying that recollection might be present later in time if only we were to look (a potentially unfalsifiable hypothesis). One problem with that is that it is improperly assuming a direct 1:1 mapping of cognitive processes to ERP effects, de facto assuming that the timing of one effect (ERPs) must coincide with the timing of another effect (the cognitive process). However, it is widely understood that ERP effects of memory are considered to be covert measures that occur separate to the overt subjective experience of conscious memory reported by the subject ^45,66,71,72,79,132^. Thus, it requires acknowledging that both item-familiarity and recollection are both normatively occurring in a delayed manner evident in their reported response times (in the present data: 2.4 seconds, 1.8 seconds, respectively) that is preceded in time by their respective ERP correlates (e.g. ∼400 msec, ∼800 msec, respectively). That is, all recollection is delayed, at least in the sense that the traditional recollection condition’s responses are normatively delayed from the earlier ERPs effect correlates that are well-established as the LPC. Why then would it now be something different and special for this one condition of accurately retrieving the source-especially when considering that the context-familiarity source memory responses are notably *faster* (e.g. happening less delayed) than both recollection and item-familiarity? The post-hoc modifications needed to support the delayed recollection account’s uniqueness to the context familiarity condition while leaving the canonical ERP effects of item-familiarity and recollection unaccounted for are thus deemed untenable.

### Is the BCN a case of the Late Parietal Negativity (LPN)?

Another common suggestion about the broad central negative (BCN) ERP effects in the present study is that they could represent the Late Parietal Negativity (LPN)^7^, which is a heterogeneous effect sometimes seen in studies of source memory and other tasks (for Reviews see ^133,134^). The LPN has been linked with a controlled search process for episodic memory and is coarsely thought to reflect the mental processes of re-constructing the prior episode. As a brief primer, the LPN was proposed to reflect a reconstructive or evaluative process in memory search when memory features are not fully recoverable ^133–135^. Prior work studying the LPN has identified early and late variants of the LPN, with the earlier variant occurring approximately 600 to 1200 ms post stimulus onset and being variously interpreted as a) a mnemonic search for context ^136^, b) re-activating context-specifying information from an encoded episode ^135,137^, or c) during later epochs that we did not measure (1200-1900 ms), associated with the evaluation of fluency ^76,136,138^ or d) reflective of the absence of information (1200 to 2400 ms) ^139^.

The results here may appear at first glance to be similar to the late parietal negativity (LPN), in that they are negative-going ERP differences that occur relatively late in the epoch, and extend to some parietal regions (though observed here & previously to emanate from foci in more central regions ^1^). As such, the present findings could potentially be seen to fit one/some of those 4 different accounts of the LPN’s functional significance. However, upon closer inspection there are several reasons, described fully below, of why the LPN interpretation does not fit the observed patterns in the present data sets. Thus, the LPN interpretation remains problematic and difficult to reconcile.

First in determining if the present study’s broad central negativity (BCN) might instead be the same effect as the late parietal negativity (LPN) is a direct appeal to what the authors of the primary Review of the LPN have said about the matter ^134^, where it was specifically noted that the LPN was *not* the same effect: *“… in the studies by Addante et al. (2012a,b)… these components are unlikely the same ERP component as the LPN.”* (page 631), and *“Addante et al. (2012a) examined ERP correlates of source memory for items that were recognized low confidence… the rather broad scalp distribution and the early onset of the negativity do, however, not necessarily support an LPN interpretation of this effect”.* (page 634 ^134^). The original report of the negative ERP effects of context familiarity ^1^ also similarly provided why the two were not the same effects, concluding that: “…*these factors make it difficult to attribute the observed effects as an LPN” (page 448).* So, neither of the authors of either of the two effects thought they were measuring the same things, and this bears some weight towards a measured interpretation of the findings.

Another possible account of the present findings as an LPN effect is that they may reflect a “controlled search process” that is one of the core (albeit vague) functional significances ascribed to the LPN ^134^. There are several lines of reasoning weighing against this possibility. In theory, every trial of the study should be inducing participants to engage a ‘controlled memory search’ simply by virtue of the memory question being asked of them. That is, the participants are expected to be searching their memory on every trial for if they remember the item and the source of its encoding condition, then reporting it. In that sense, the controlled memory search would be a variable that is held constant across all conditions, thus canceling out in the contrasts, and leaving open the question of why this particular memory condition would show the negative ERP effect when compared to other conditions that also involve the searching of memory do not show the effect. Furthermore, if the present effect was representing a controlled search process, then it would have also presumably been seen in the recollection condition when compared to correct rejections, if participants searched memory for controlled recollection of the information. However, the same negative ERP effects weren’t seen in the recollection condition (which had far more trials included for statistical power, too), nor were the same effects seen for the item-familiarity condition in which participants were searching for the unknown source information as well, as described further below.

Expanding upon this line of reasoning, consider the item familiarity condition (i.e. successful recognition memory hits, but lacking source memory): participants report that they do not have source memory and spend significantly longer response times on the source memory judgments. If participants had made a correct item judgment on the basis of item familiarity, and could not recollect the source of the event, then presumably they would certainly be engaging in a controlled search process of their memory to see if they could find that source information. This provides a solid premise to infer that the condition would also include a “controlled search of memory” for the source information that still comes up empty for them, and makes a clear hypothesis that we should see a major negative-going ERP effect on these trials if it reflected the same thing as the LPN ^134^. Yet, that is not what the present data shows, nor in several of the independent replications of the finding, and instead we see that the item familiarity condition differs reliably and consistently from the context familiarity condition, and that they each reliably differ from a baseline condition of correct rejections. Hence, the present differences must be attributable to another variable than just a controlled search of memory, and this creates a challenge for the LPN interpretation.

An additional point of differentiation of the present results from the LPN is the different patterns of topographic and temporal activities. Differences in topographical and temporal profiles of neural correlates of behavioral responses has been widely taken to reflect dissociable cognitive processes ^44–46,66,69–72,114^. The LPN is generally found by other studies to be maximal in posterior parietal areas, whereas the current effect was observed in superior central regions of the scalp and distributed broadly to both frontal and central-parietal areas, and had been previously differentiated from posterior sites ^1^. In regard to timing, the LPN has been previously characterized as beginning during periods after the epoch ended in the present study (1200 ms)^134^. Among those irreconcilable factors was sensitivity to memory conditions: the LPN has been reported to either be invariant to source accuracy (i.e.: it was present for both correct and incorrect conditions ^136^) or even to be larger for incorrect source judgments ^140^. However, the current conditions’ effects have been found to be specific for correct source memory when compared directly to incorrect source memory ^1^. Furthermore, other lines of evidence weighing against the LPN account of our findings comes from testing a direct prediction made that the LPN should be present “*not necessarily tied to successful memory retrieval but also observed during misses…”* (pg. 635 ^134^). We did not find that prediction supported in the present data, as the present findings directly dissociated ERP effects of misses from the ERP correlates of context familiarity, which was also replicated, and thus adds to the considerable line of evidence weighing against an interpretation of the present findings for context familiarity as an LPN.

A related problematic issue for an LPN account of the present findings is that the LPN is well-researched but poorly defined for its functional significance. The LPN is operationalized broadly as either a “reconstructing of the past” or a “controlled search process”, which, colloquially, only means “trying to remember”, which should occur on every memory retrieval trial to one extent or another and is difficult to reconcile with the present pattern of results. Due to the poor operational definitions available to the LPN, it is possible that the LPN findings from at least some other studies could instead be reflecting context familiarity ^137,141^. That is, there is also the other possibility that some instances of reported results interpreted as an LPN might have actually been misinterpreted and reflecting the effects of context familiarity, instead of an LPN. In such cases, misinterpretation of effects could be due to lacking paradigmatic sensitivity (i.e.: being able to dissociate influence of item familiarity from context familiarity, or recollection) or having lacking a theoretical framework to understand results as reflecting context familiarity (i.e.: if one was only considering the existence of two episodic memory processes of recollection and generic familiarity).

For instance, another laboratory’s study of contextual memory retrieval confidence of source information, their findings of a broad, Cz-centric ERP negativity at approximately 1000 milliseconds occurring for source memory retrieval were interpreted as an LPN in that study ^137^ but their results’ characteristics could also be interpreted as reflecting context familiarity instead, given the successful retrieval of the context of source memory attributes in that study. Leynes et al (in press) also report broad central negativity effects occurring from 500-800 ms post-stimulus in their Experiments 2 and 3 for recognition hits being more negative than correct rejections for the conditions of testing context in a masked word priming study, which could be seen context familiarity influencing mnemonic processing, much as was reported in an ERP study of cued recall with semantic primes ^84^.

In sum, the main, non-specific feature shared among the present findings and the LPN is that each is a negative-going ERP effect-but that in itself is not a very helpful descriptor, just as many other positive-ERP effects are not assumed to reflect the same thing just by virtue of their being a positive difference. The present findings are clear that the broad central negativity observed in the present studies is evident to the specific condition of context familiarity and not reasonably ascribed to other negative-going ERP effects such as the LPN, nor cognitive operations associated with the LPN such as a controlled search process or reconstruction of the past, concurring with the prior observations of their differentiation ^1,134^, respectively.

### Are BCN effects driven by fluency or guessing?

It is also possible that the current results might reflect a form of guessing. In this view, effects may be reflecting conceptual implicit memory contributions to source memory, as has been seen in the form of guessing observed in prior studies of item recognition ^88^. However, ERP findings of guesses indicated that guess processing was occurring much earlier in time (200-400 ms and 600-700 ms) than what we found in the current study (800-1200 ms) ^88^, so the current findings do not bear those hallmarks of guessing. The fact that two explicit declarative responses (item, source) were observed for judgments in the condition is also inconsistent with guessing, too, as is the above-chance performance of the responses. While it’s possible people could have been successfully guessing in these item + source judgments, this possibility becomes increasingly less likely when considering that participants were given the option of saying they ‘don’t know’ the source in our task’s paradigm (i.e.: ‘source unknown’, Figure 1), and which are thus excluded from the condition.

Furthermore, if results were influenced by guesses, then it is notable that the purportedly accurate ‘guesses’ would actually be driven by an implicit memory process (i.e.: ‘recognition without awareness’). We conducted several direct analyses to assess if context familiarity ERP activity resembled the known hallmark signs of other implicit processing (i.e., misses) that would have been presumably supporting the guess behavior, but instead found that it was significantly different in both time and scalp location from implicit misses. That finding is further bolstered by another, different implicit activity recently observed in the same dataset for a purer form of implicit memory (IMAP) ^66^ that was found instead to manifest in right parietal regions in positive-going ERP effects from approximately 400 to 1000 ms (not the fronto-central sites observed here from 800 to 1200 ms in negative-going ERP effects). Prior work discussing the requirements for a condition being ‘recognition without awareness has established the criteria that information about context should be absent from the memory ^142^ – which also rules out the condition of interest for context familiarity in the present study. Taken together, the evidence weighs against interpretations of context familiarity being primarily the result of processes associated with guessing from underlying implicit memory signals.

### Are the negative ERP effects reflecting implicit fluency?

It is probable that, much like many explicit memory processes, context familiarity is also supported by forms of implicit processes (i.e., repetition priming, perceptual priming, contextual fluency) that then give rise to the explicit memory judgments, as has been found in other studies of item familiarity and cued recall ^84,143,144^. Familiarity of context may naturally represent various forms of fluency in processing information which has linked negative-going ERP effects for old items to repetition and perceptual fluency ^50,89,90,119,145^. As such, it is possible that these explicit response trials for context familiarity could be supported by an underlying fluency heuristic that leads one to accurate recognition and source judgments.

According to this explanation, the negative ERP effect likely reflects the fluency heuristic. This effect might purportedly be obscured during high-confidence trials by the predominance of positive LPC effects associated with recollection, which could override the presumed baseline negative ERP activity postulated to occur in recollection conditions as well. While such an explanation is hypothetically conceivable in this familiar context, where various possibilities exist, it remains speculative and lacks parsimony with the data. This is because it fails to explain why conditions of item familiarity did not display similar negative ERPs in the apparent absence of recollection. Additionally, the effects of context familiarity manifest later in time (800-1200 ms) than the very early latencies (200-400 ms) where fluency is often found exerting its effects ^75,90^. Through the series of systematic analyses here we demonstrate that there are still core distinctions between the kinds of familiarity represented by items and context, both of which surely are multiply determined by earlier sensory and contributory processes. We believe that based upon the combination of converging data and the divergent findings for studies measuring fluency at much earlier latencies, locations, and conditions, the present findings cannot be explained by mere fluency.

### Future Directions

There are several next steps which can be envisioned for future research based upon the present findings. For instance, intracranial measures of subsections of the medial temporal lobe regions such as the parahippocampal cortex and perirhinal cortex ^146–155^ as hypothesis-driven tests of model predictions from the present work (see Section 5.5 below, Fig. 8). Such kinds of studies can include both invasive and non-invasive stimulation approaches to both enhance ^156–166^ and disrupt ^156^ memory operations of different episodic processes, and assessing their relevance in real-world environments/context ^167–170^. Other potential future approaches include exploring for neuropsychological dissociations among various types of MTL patients ^171^ or those with specified focal lesions of MTL subregions ^39,77,172^, or electrophysiological searching for differential oscillatory bandwidths that may possibly be selective for each process ^67,146–149,161,173–190^.

In pursuing future studies, note that the specialized condition used in the present study to measure context familiarity (low-confidence recognition hits accompanied with accurate source memory) does not necessarily mean that it is the only potential way to measure context familiarity. Rather, it is likely just one of many possible approaches to capturing an elusive cognitive process. The findings here suggest the importance of including multiple measures of memory into experimental protocols, as it permitted the leverage needed to identify the combined conditions needed to measure context familiarity in the present experiments. We would expect future studies to find context familiarity to accordingly be measurable in other ways pending properly sensitive experimental designs that do not conflate it with recollection or item-familiarity or guessing.

For example, we surmise that context familiarity could be found in future studies using different paradigms that systematically vary repetition of context while holding items constant and vice versa, similar to repetition paradigms used in prior fMRI studies ^2,35,37,68,191^ that found results similar to the present data. Our measure here was a combination of multiple responses (memory confidence scales) across multiple measures (item and source memory judgments), and thus reflects the sophisticated paradigmatic sensitivity that will likely be needed for follow-up explorations. Towards this end, studies should use an unequal confidence scale for source memory judgments to permit a ‘source unknown’ response in order to avoid the guesses that are inherently integrated in balanced confidence scales ^88,192–194^; that is, be sure to move from a commonly-used 6-point confidence scale ^35^ to a 5-point confidence scale ^1,65–67^. That is, including a ‘source unknown’ option into source memory tests should be essential for future researchers to avoid contaminating source memory responses with instances of guessing or the source unknown.

Another direction for future studies could be to assess the possibility of delayed recollection supporting the condition identified here as context familiarity. One way to do that could be by studying response locked analyses of ERP or oscillatory analyses ^174–178,195^, as opposed to the stimulus-locked events studied here for the purposes of consistency to the literature of known ERP effects of episodic memory ^42,43,45,46,57,70,84^. But it is important to also note that such studies of response-locked activity in this paradigm risk being confounded or conflated in their baseline period due to the preceding activity of the item recognition prompt/decision in in the present paradigm-so future studies would need to be very carefully designed to ensure those and other factors are properly disambiguated.

### Implications for models of memory

The medial temporal lobe region of the parahippocampus has been modeled by several frameworks to be a neural substrate supporting contextual representations of episodic memory ^6,15,16,196,197^ (Figure 11).This is based upon neuroimaging studies identifying activations of that region for contextual processing ^35,37,38^, while ascribing item familiarity to the adjacent perirhinal cortex ^37,68,191,198^. This has been paralleled by findings from studies using intracranial recordings showing temporal dissociations of familiarity and recollection processing amid the parahippocampal gyrus and hippocampus, respectively ^108,151,152,199,200^, which has been similarly observed using functional neuroimaging methods ^150^.

**Figure 11.**
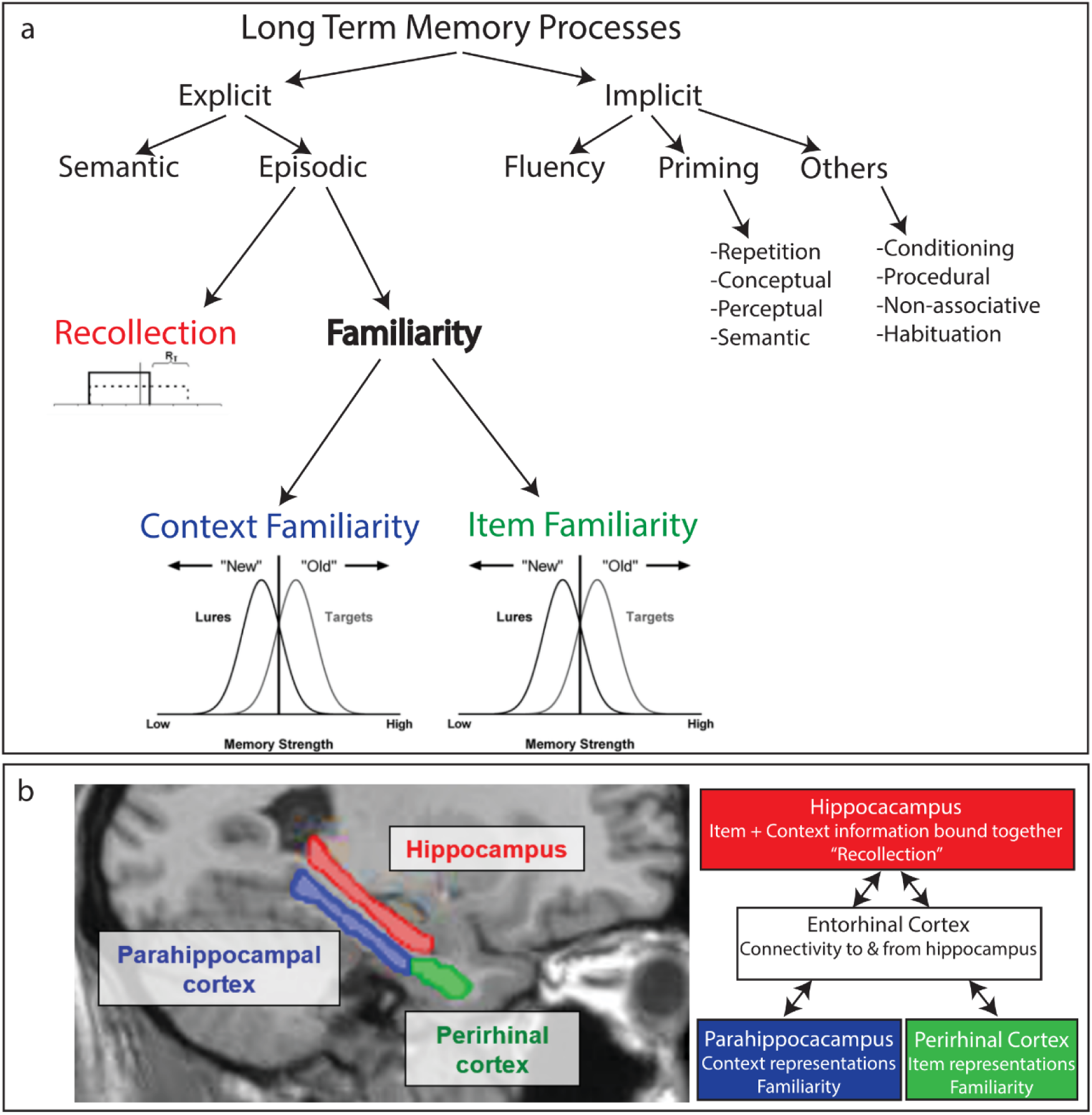
Physiologically informed model of long-term memory processes. Top, A: The tri-component model presents an updated representation of the canonical declarative memory framework that is inclusive of physiological findings from physiological data indicating that recollection and familiarity are distinct processes of episodic memory ^39,44^, and that context- and item-familiarity are also dissociable processes in episodic memory (Figures 4, 5, 6, 7). Model depicts the emerging views that differential forms of implicit fluency contribute to driving explicit familiarity ^75^. A sample schematic of recollection shown models it as a threshold process based upon physiological findings in the current and former data indicating that is limited to high-confidence responses and does not extend to low-confidence responses even if source memory is retrieved ^1,31,51^. Context familiarity and item familiarity are each depicted as continuous processes of signal detection, with representative distributions of old and new information each. Bottom, B: Anatomical and conceptual representation of medial temporal lobe processing of mnemonic information, based upon the ‘binding-items-in-context (BIC)’ model ^6,15^, updated here to include differential familiarity processes for items and context in the perirhinal and parahippocampal cortices, respectively. Note that the model presents a simplified representation of the breadth of complex memory operations and neuroanatomical substrates.

Neuropsychologically, converging reports of the perirhinal cortex role in item familiarity come from a rare clinical patient, NB, with an epileptic resection lesion specific to the perirhinal cortex that left the parahippocampal cortex intact ^201^. This patient was deficient in familiarity for items (words) but showed intact abilities of familiarity for pictures and faces ^172,202–204^, which we consider here as broadly representing kinds of context in episodic events and this could be seen as performance consistent with the patient’s intact parahippocampus service of context familiarity (though it is possible that faces and/or picture stimuli might have potentially instead be treated by that patient as an item in certain studies with other contexts, or alternatively the face/picture memory performance might have potentially been supported by the other hemisphere’s intact perirhinal cortex). Similar reports of familiarity for contexts in the absence of object familiarity or recollection have emerged from neuropsychological ^205^ and normative studies using visual scenes of mazes, whereby it is the spatial layout of the scene that is experienced as familiarity and not the items or objects ^9,10,206,207^.

Together with the present studies, these findings converge to suggest an updated framework for the organization of long-term episodic memory from the canonical declarative memory model used ubiquitously throughout textbooks, following from the BIC model that had originally motivated the hypotheses tested in the present work ^15^. We propose an updated model to conceptualize the organization of specific episodic memory processes among a broader framework of long term memory (Figure 11), which is primarily based upon being specifically physiologically-informed, whereas most prior models of long term memory have instead been either informed primarily by cognitive psychology, neuropsychology, or neuroanatomical/neuroimaging studies ^6,15,17,25,26,208–212^ (though see ^75^ for an example of a physiologically informed model of just familiarity-specific memory processing).

Our new tri-component model provides an updated representation of the canonical declarative memory framework and BIC model, that is now inclusive of findings from physiological data indicating that recollection and familiarity are distinct processes of episodic memory ^1,39,44,45^, and that context- and item-familiarity are also dissociable processes in episodic memory (Figures 4, 5, 6, 7, ^1,3^). This physiologically-informed model also encompasses the emerging views that differential forms of implicit fluency contribute to explicitly-aware familiarity ^75,89,118–120,123,145,213^ and that other forms of implicit memory depend upon the hippocampus 66,77,214-220.

In our proposed framework (Figure 11), the episodic process of recollection is modeled as a threshold process based upon physiological findings in the current and former data, since the LPC effects were found to be limited to high-confidence responses and did not extend to low-confidence responses even if source memory is retrieved ^1^; for other related cognitive frameworks see ^30,51,56^. Context familiarity and item familiarity are each depicted as distinct continuous processes of signal detection for relatively old and new information distributions ^1,114,115,221^, based upon the current and prior data, and as opposed to the traditional model of them both merged within a single continuous process of general familiarity. The model also updates the understanding of the parahippocampal and perirhinal cortices as processing differential familiarity signals (Figure 11B), as opposed to previous models depicting them to operate amid a single familiarity signal along both regions of the larger parahippocampal gyrus 6,15,17,20,21,197.

How might the subjective experience of context familiarity occur during cognition? We draw upon our proposed additions to the BIC model/variants ^6,15,17^ to suggest a cognitive framework that can be built to understand how the condition of context familiarity may arise or manifest (note: this is a proposal of how it might occur in some instances but should not be taken as suggesting that it is how recognition works in every case). This account both echoes and extends models provided by prior work investigating the role of context in episodic memory ^1,16,22,191^, while integrating fluency heuristics ^75,89,119^. It has been proposed that people use a fluency heuristic to make recognition judgments ^75,89^, and the current findings extend that by suggesting the possibility that a fluency heuristic can also be used to support accurate source memory judgments, too. This would be based upon an underlying fluency for context, as opposed to fluency for items or conceptual/perceptual dimensions.

By our account, as a retrieval probe is first shown on a screen by itself, the item doesn’t at first seem familiar (i.e., no FN400 for item familiarity from ∼300-600ms that we observed). The context, however, may seem familiar, even if they do not recollect the details of either the item or how the context was bound with other details from the prior episode. Akin to how item familiarity can be derived from underlying fluency of the stimuli, so too would context familiarity be facilitated by a separate underlying fluency for the context. If the context does not initially seem familiar from ∼300-600 ms, then in the ensuing time (600-900 ms, LPC) the participant would continuing to mentally look back in time for any context in which to place the item being shown on the screen (e.g. ‘where might I have seen that before?’), seeking the pattern completion process to autocomplete a recollection of the relevant details from the prior event. During that time (600-900 ms), that autocompletion process of binding any existing neural signals of context together with the presented item fails (i.e.: our observations of an absence of evidence for the LPC correlate of recollection), consistent with the slower response times observed for these item judgments.

As time continues (800-1200 ms) the person continues to assess their memory for any connections of details, something about the stimuli remains diagnostic for prior exposure (the broad central negativity effect emerges to reliably differentiate from correct rejections and item familiarity), which could be either the explicit familiarity of a contextual detail such as the encoding source or the implicit sense of its contextual fluency for associated information about the item. The successful retrieval of some familiar contextual information would be enough to logically deduce that the item must have been old ^4,5^. That is, one can then draw upon that awareness that the context seems familiar to derive the logical inference that the item is likely old, akin to how other studies have found item familiarity can lead to false alarms in associative recognition by similar logical deductions ^12,222–225^. Ultimately, this basis of a retrieved prior context is sufficient to support an accurate hit in item recognition, albeit with low confidence because the item itself is not remembered directly, while the correct source is identified in the ensuing judgment based upon that retrieved familiarity of the context of its prior exposure ^5^. The item is thus accurately recognized due to the context seeming familiar: context familiarity.

Importantly, our tri-component account provides an improved precision in characterizing human episodic memory, and as such could help to better understand disparate findings from memory studies and neuropsychological patient impairments that occasionally arise in the literature. That is, if prior studies had been conflating distinct familiarity processes for items and context, and conflating context familiarity with instances of recollection, when occurring in differential proportions and under differential experimental conditions, it could easily produce disparate clinical findings that have been reported about source memory, context, recollection, and neuropsychological impairments thereof ^6,17,24,26,29,39,171,172,203,204,211,226–229^. If we are now able to delineate between these three distinct but overlapping processes of memory, it could help illuminate a better understanding of the nature of memory impairments and lead to better precision in types of amnesia diagnoses.

The idea that there can be different kinds of familiarity for items and for contexts has been previously mentioned by several others ^1,4,16,64^. Whereas prior work established the premise that there can be separable familiarity of items and familiarity of contexts ^4,16^, they nevertheless proposed that each was stemming from the same singular memory process of underlying familiarity. For instance, the Discrepancy Attribution Hypothesis offered that people can produce either piece of mnemonic information separately ^4^, stating:

> “contextual information can also cause the person to experience a feeling of familiarity: encountering a stimulus in a context that is unusual for that stimulus (e.g., a clerk on the bus) can cause this.” However, and importantly, they go on to propose that “**the use of contextual information to perform judgments about the past is not qualitatively different from the use of identity** [emphasis added]” (page 561) ^4^.

Here, we put forth a fundamentally different view: that production of familiar context *does* differ qualitatively from the production of familiar item information and provide converging lines of support that the familiarity of items and familiarity of contexts are evidently operating as separable independent neurocognitive processes.

### Limitations

ERP studies are inherently limited in spatial resolution of their topographic distribution of effects and are generally unable to provide source localization of the subdural cortical and subcortical source generators of the effects recorded at the scalp ^79^. Therefore, spatial distribution of the physiological effects reported here should be expected to have a reasonable amount of variability in future studies, and may vary upon different experimental factors, protocols, and paradigms. While the physiological effects of the present study’s main Experiment 1 ^65,66^ were found to be reproducible in independent data sets collected from different demographics in different universities (Experiment 2) ^1,39,67,77^, Experiment 2’s archival datasets were unable to collect meaningful response times to memory judgments due to a protocol that asked participants to withhold their response until after viewing the memory probe for 1.5 seconds, thus rendering the response time uninformative and unable to asses for reproducibility of the results from Experiment 1. Therefore, while the physiological effects that formed the backbone of the present study were independently replicated, the behavioral results remain to be assessed for the extent of their potential replication among future studies of larger samples.

### Conclusions

The field of memory research is replete with complexities in memory behaviors in normative and clinical patient populations that are difficult for existing theories to fully account for. We show here that the common measure of source memory can be supported by a disparate process that is independent of both recollection and item-familiarity. The memory condition of context familiarity was found to be distinct from item familiarity, in addition to previously identified dissociations from recollection. This conclusion was evident from triple dissociations observed in both behavioral and physiological measures. Moreover, the differences in item-familiarity and context familiarity reflected differential neural processing at both the individual subject level as well as trial-wise differences in memory processing that was identified within-subjects and were replicated across several independent studies. These findings cannot be accounted for by relying upon using only two processes of episodic memory for interpreting the results ^33,52^. Item familiarity and context familiarity are therefore seen as fundamentally different kinds of familiarity processes independently operating in service of episodic recognition ^44,69–73^: one process for items and another for context, and each distinct from recollection. This extends traditional dual process models of recognition to reflect that context familiarity constitutes a third process of episodic memory.

## Supporting information

4th Response to two (new) Reviewers

Third Response to Reviewers at NatureCommsPsych

Second Response to Reviewers at NatureCommsPsych

First Response to Reviewers at NatureCommsPsych

## Acknowledgments

Funding support of this work came from the National Institutes of Health (NIH), which included grants 2 L30 NS112849-03, 2 L30 NS112849-02, and 1 L30 NS112849-01 awarded to RJA from the National Institute of Neurological Disorders and Stroke (NINDS), from Florida Institute of Technology (FIT) grants to RJA from the Institutional Research Incentive (IRI) program. Authors thank the Florida Tech Provost Office of Dr. Marco Carvalho for generously providing direct funds to support the project. DLD received support from grants R01 NS124585 from the NIH/NINDS and A1321808 from Medtronic, Inc. JPC and RJA received support from a NASA/FL Space Consortium Grant. Authors thank Alana Muller, Lindsey Sirianni, Anderson Wilder, and Cristo Rey for research assistance, and are grateful for the constructive contributions of six anonymous- and one non-anonymous Reviewers. Publication of this article as Open Access was funded by the Open Access Subvention Fund and the John H. Evans Library at Florida Institute of Technology and the Office of the Vice Provost for Research.

## Author Contributions

RJA designed the study, supervised the data collection, analyzed the data, created the figures, interpreted the findings, wrote the manuscript, and responded to Reviews. EC and JPC contributed to data analysis. RW, JB, and DLD contributed to interpretation of findings and manuscript preparation.

## STAR Method

### EXPERIMENTAL MODEL AND STUDY PARTICIPANT DETAILS

#### Experiment 1

The current study (Experiment 1) consisted of a re-analysis of previously published ERP data ^65,66^, so the sample size was determined by the number of participants’ data that was available in the archival dataset; the study was not pre-registered. The participants consisted of 61 right-handed students free from neurological disorder and memory problems. Data was not used for four participants due to noncompliance issues (i.e., pressed only one button throughout the task or ignored experimenter’s instructions), data lost due to experimenter error (N=1) or EEG data deemed unusable (N=2) due to excess motion artifacts/noise that resulted in an exclusion of the majority of EEG trials. This presented a working data set of N = 54 for the current study, which was more than double the size of the original study reporting physiological correlates of context familiarity (N = 25) ^1^, and triple the sample size standards of many of the foundational ERP studies of recognition memory ^40,44,101,114,231–235^.

As described by the data’s original publication ^65^: Participants were recruited through a combination of methods including advertisements placed around CSUSB or through the schoolwide research pool SONA. Participants recruited through advertisements were paid $10 an hour for sessions that lasted approximately 2 hr. The majority of participants were women (N = 48); 57% were Hispanic, 23% Caucasian, 11% Asian, and 10% identified as more than one ethnicity. The average age was 23.5 years old (*SD* = 4.82). Findings do not apply to only one particular sex, gender, or demographic; the study was a secondary analysis of archival data ^1,39,65–67^, and thus was constructed to be neutral in consideration of any particular sex or gender or other demographic factor. Inclusion of participants was based upon the nature of the archival data that was available (we included all data sets available), which had included anyone who volunteered to participate for the research from broad advertisement efforts on the campus of California State University – San Bernardino, and determined based upon self-reporting. Since the pursuit of the present study was one that sought to assess the reproducibility of prior findings, analyses were performed agnostic to sex and gender or other demographic factor in order to faithfully reproduce prior findings of that had adopted the same approach. There are no known reasons for why an absence of demographic-specific analyses would limit the study’s generalization, and the findings were observed to be reproducible across several independent studies that comprised different demographic proportions and produced results in line with traditional findings of ERP effects of memory in the field across many labs, countries, and demographic distributions since the early 1990s.

None of the participants reported any visual, medical, or physical issues that would interfere with the experiment. Most participants spoke English as their first language (N = 47) and those who had indicated speaking a different first-language had been speaking English for an average of 16.73 years (*SD* = 4.74, SE = 1.22, Median = 17.0, Minimum = 7, Maximum = 25). Written informed consent was obtained for participation in the experiment and protocol approved via the Institutional Review Board of California State University – San Bernardino, and data analyses observed the privacy rights of human subjects, consistent with the Declaration of Helsinki and as approved by the Institutional Review Board of Florida Institute of Technology.

#### Experiment 2

The participants included in Experiment 2 were aggregated across three previously published studies ^1,39,67,77^ that used nearly identical experimental protocols of item recognition confidence followed by source memory judgments as were used in the present Experiment 1 ^66^, for a total sample of N = 56. These studies had only directly contrasted conditions of recollection and context familiarity, but had never directly assessed if the ERPs for item- and context-familiarity differed ^64^. Each study’s cohort of participants have been previously reported for having intact levels of familiarity-based recognition on both behavioral and electrophysiological measures that did not significantly differ as a function of younger/middle aged adults ^39,64,77^. They have therefore been previously treated as equivalent cohorts of familiarity-based recognition, which was the subject of the present investigation ^39,77^, and thus we also treated them accordingly here as well.

The participants were reported in prior studies ^1^ as consisting of twenty-five healthy right-handed undergraduate students (Mean age = 20.4 years [SD = 2.9, SE = .60, Min = 18, Max = 30], mean education level = 14.4 years [SD = 1.8, SE = .4, Min = 12, Max = 19] demographics of 52% Caucasian, 20% Hispanic, 20% Asian, 8% more than once race; seventeen female) and twenty-two ^1,67^ right-handed undergraduate students (Mean age = 20.9 years [SD = 3.1, SE = .70, Min = 18, Max = 29], Mean education 14.5 years [SD = 1.4, SE = .30, Min = 13, Max = 17], demographics of 55% Caucasian, 9% Hispanic, 5% African-American, 32% Asian; ten females), all recruited from the University of California–Davis Psychology Department subject pool. They received credit for participation and were free from neurological, visual, motor, or other medical disorders. An additional nine middle-aged participants (six female; Mean = 45.4 years, SD = 12.5, SE = 4.2, Min = 27, Max = 57) were included whom were tested on the same memory protocol (six male) ^39,77^ and had a comparable profile of education (M = 16.8 years, SD = 5.7, SE = 1.9, Min = 12, Max = 31) and majority demographics (78% Caucasian, 22% more than once race). The experiments had been conducted as approved by the University of California– Davis Institutional Review Board protocol for research on human subjects.

## METHOD DETAILS

The paradigm used was the same item- and source-memory confidence paradigm that has been routinely used in our prior reports to characterize various facets of memory ^1,39,65–67,77,165^, with slight modifications as noted below ^65^. This paradigm consisted of an encoding phase containing four sequential study sessions during which participants studied 54 words per session, followed by a retrieval phase that contained six test sessions in which the participant’s memory was tested for 54 words in each session. Item recognition was tested on a five-point confidence scale with ‘1’ indicating being sure the item was new, and ‘5’ indicating being sure the item was ‘old’ (Figure 1).

During the encoding phase, participants were given instructions to make a simple decision about the word presented. The participants were either asked to judge if the item was man-made or if the item was alive. The instructions were presented in one of two counterbalanced orders: ABBA or BAAB. The participants viewed four lists of 54 words during the encoding phase. After the encoding phase was complete, the EEG cap was applied to the scalp of participants. During the memory test, participants viewed a total of 324 words, 216 of which were presented in the encoding phase and 108 of which were unstudied items (new items). All words presented during study and test were presented in white font on black background screen. Word stimuli were selected from the same source as stimuli used in the original study in 2012 and preserved the same characteristics. Word stimuli were selected from the Medical Research Council Psycholinguistics Database (http://www.psych.rl.ac.uk/MRC Psych Db.html). Word stimuli were all nouns, had an average rating of concreteness of 589 (min=400, max=670), imageability of 580 (min=424, max=667), Kucera–Francis Frequency of 30 (min=3, max=198), and an average number of 4.9 letters in each word (min=3, max=8).

During the memory retrieval phase, the participants were read instructions asking them to judge if the stimulus word presented was old (studied during the encoding phase) or new (not studied before in the encoding phase) (Figure 1). To begin a trial, a screen with a small white cross at the center was presented for one of three randomly chosen times: 1 second, 2.5 seconds, or 3 seconds. Then the participants were presented with a word in the middle of the screen, the numbers 1, 2, 3, 4, and 5 evenly spaced beneath the word, the word “New” on the left by the number 1, and the word “Old” on the right under the number 5. Participants pressed any number between 1 and 5 to indicate if they confidently believed the word was old (5), believe the word was old but was not confident (4), did not know if the word was old or new (3), believe the word was new but was not confident (2), or confidently believed the word was new (1). This prompt was subject-paced, and participants were told to choose the response that gave the most accurate reflection of their memory, and to respond as quickly and accurately as possible.

Immediately after each decision on item memory confidence, participants were asked to answer a source memory question about if the word came from the animacy decision task or the man-made decision task. The word and numbers remained on the screen but this time, word “Alive” was presented on the left by the number 1, and the word “Manmade” was presented on the right under the number 5. Participants were told to choose the response that gave us the most accurate reflection of their memory and could respond that they confidently believed the word was from the animacy task (1), believe the word was from the animacy task but was not confident (2), did not know the source of the word or had replied in the question directly before that the word was new (3), believe the word was from the manmade task but was not confident (4), or confidently believed the word was from the manmade task (5). Trials were terminated by subject response. Each session consisted of a list of 54 words. Six lists of 54 words were presented during the retrieval phase. The retrieval phase of the paradigm also included a simple metacognitive question to participants for estimating their performance, which occurred once every ten trials and for which the data has been previously reported elsewhere for metacognition effects ^65^. The current investigation focused instead upon the memory related responses, as the metacognitive response data has been reported previously. Each session consisted of a list of 54 words; overall, six lists of 54 words were presented during the retrieval phase.

### Conditions of interest

Following common convention in the field, we operationalized these processes based on the combination of responses given for item recognition confidence and source-memory judgments, consistent with the way prior research has defined these conditions. For recollection, we followed the convention of prior findings that it represented instances of retrieving high-confidence item recognition followed by correct source information: item responses of ‘5’ accompanied by source judgments that were correct (Figure 1; ^1,39,65^. For context familiarity, we followed the same approach used to originally identify this process ^1^: instances of low-confidence item recognition accompanied by correct source judgments (Figure 1). Item familiarity was defined based upon previous work ^66,67^ as instances in which participants retrieved successful item recognition (hits, combining 4 and 5 responses) but reported having no source memory information (source unknown, 3 responses) (Figure 1). These conditions of item- and context-familiarity are ones which have never been directly compared against each other in the literature while using behavioral or ERP measures, and so it remains to be determined if and how they may differ, and the extent to which the process of familiarity may be organized among these variables as a continuous or distinct process(es).

Response times for item recognition and source memory responses were measured as follows: item recognition times were measured from the onset of the memory probe (i.e. the word) to the button press of the recognition decision; source memory response time was measured from the onset of the source memory prompt (which happened immediately upon the end of the item recognition judgment) to the button press of the source memory decision.

### Electrophysiological Acquisition

Each subject was tested individually inside a private chamber. Stimulus presentation and behavioral response monitoring were controlled using Presentation software on a Windows PC. EEG was recorded using the actiCHamp EEG Recording System with a 32-channel electrode cap conforming to the standard International 10– 20 System of electrode locations and was acquired at a rate of 1024 Hz. The EEG cap was sized while the participant’s face was wiped free of skin oil and/or makeup in preparation for attaching ocular electrodes. Five ocular electrodes were applied to the face to record electrooculogram (EOG): two above and below the left eye in line with the pupil to record electrical activity from vertical eye movements, two on each temple to record electrical activity from horizontal eye movements, and one electrode in the middle of the forehead in line horizontally with the electrode above the left eye as the ground electrode. EOG was monitored in the horizontal and vertical directions, and this data was used to eliminate trials contaminated by blinks, eye-movements, or other related artifacts. The EEG cap was placed on the participant’s head and prepared for electrical recording. Gel was applied to each cap site and impedances were lowered below 15 KOhms via gentle abrasion to allow the electrodes to obtain a clear electrical signal. Subjects were instructed to minimize jaw and muscle tension, eye movements, and blinking.

### Electrophysiological Analysis

Physiological measurements of brain activity were recorded using EEG equipment from Brain Vision LLC. All EEG data was processed using the EEGLAB and ERPLAB toolboxes in MATLAB ^236,237^. The EEG data was first re-referenced to the average of the mastoid electrodes, passed through a high-pass filter at 0.1 hertz as a linear de-trend of drift components, and then down sampled to 256 hertz (Hz). The EEG data was epoched from 200 milliseconds prior to the onset of the item recognition stimulus to 1200 milliseconds after the stimulus was presented ^65^. Independent components analysis (ICA) was performed using Infomax techniques in EEGLab ^238^ for artifact correction and the resulting data was individually inspected for artifacts, rejecting trials for eye blinks and other aberrant electrode activity. During ERP averaging, trials exceeding ERP amplitudes of +/-250 mV were excluded, as described in the previous report of this data ^65,66^. A 30 Hz low pass filter was applied to each subject’s ERPs as a non-causal, infinite impulse response (IIR, Butterworth) filter, implemented through ERPLAB toolbox ^237^. In order to maintain sufficient signal-to-noise ratio (SNR), all pairwise comparisons relied upon including only those subjects who met a pre-determined criterion of having a minimum number of 12 artifact-free ERP trials per condition being contrasted ^78,84,239–241^. As was noted in the Introduction, the present work is not claiming any specifically process-pure ERP measurements of memory, but instead we adopt what others ^42,45,46,57^ as well as our own previously published positions ^1,39,47,65,66,77,84^ described as the ‘putative neural correlates’ of recognition memory processes.

## QUANTIFICATION AND STATISTICAL ANALYSIS

### Measuring source memory performance

The current study’s interest focused specifically upon combinations of item + source memory responses described below. Like most studies of source memory, we wanted to know how accurate people were in discriminating between the two sources of information in the experiment, and so excluded the source unknown responses from the performance calculation of source discrimination. This procedure collapses high- and low-confidence source judgments into general ‘correct’ and ‘incorrect’ conditions. We thus calculated this source memory performance in the same way as had been done in prior studies ^1^: the total number of source correct trials divided by the total number of all correct & incorrect source memory judgments (excluding the source unknown responses) [# correct source trials / (sum of # of correct + # of incorrect source trials)]. The source memory calculation we used is akin to the standard recognition measure of memory performance (%Hits-%FA) but applied to source memory discrimination, which would require excluding the ’source unknown’ responses, since a false alarm in source memory would be a source misattribution (not the lack of source memory). When source memory discriminations occur between available options of the two sources for participants to choose from, they have a 50% chance or being right/wrong (source correct or source incorrect responses), thus chance levels are at 50% in that analysis (as opposed to 33% if assessing it relative to the three response options of correct, incorrect and unknown; see discussion below for further considerations of the limitations of such alternative calculation methods^8^).

In excluding the ‘source unknown’ responses from the calculation of performance on source memory discrimination, it excludes the trials of when people had an absence of source memory ^193,194^. This approach was taken because we did not want to know how frequently people remembered correct source information at all (i.e.: out of all of the possible times that they could have), because this number would be artificially low, but rather how well they discriminated sources that they did remember. For example, consider a scenario in which people were tested on if episodic memory for where we learned various items of information in our lives. Then consider if the performance measures for memory measured the few times that we did remember correctly amid *<u>all</u>* the times when we couldn’t remember any source at all (likely most of the time, for naturalistic settings). Most people would [falsely] appear to be dense amnesiacs unable to retrieve source information above chance-despite likely having reasonable source discrimination capabilities for the times that they could retrieve source memory information. The same is true for clinical applications of the same principle: consider cases studying reality monitoring of auditory hallucinations in schizophrenia patients, whereby the pertinent question is about understanding how well the source determination is made when a source of the information is available to the patient-not the other instances in which it isn’t.

Our approach to measuring source memory performance is a common one that is sensibly used in other source memory studies ^58^. For example, our standard approach used to measure source memory performance is not just the same as we have published with previously ^1^ but is also the same approach used by additional other studies measuring source memory performance ^242^, where they note that: “*Conditional source measures control for overall differences in recognition by calculating the probability that the correct source is identified given that the item was correctly identified as old (p|source| = source hits/source hits + source misattributions)”*. For these reasons noted above, we utilized the present method of calculation for source memory discrimination that excluded the ‘source unknown’ responses, as it provides the most appropriate treatment of participant’s memory while minimizing the extent of possible confounds. This approach also avoided the potential confounds of not knowing what response criterion a subject is adopting when making the ‘unknown’ response for source memory, as no two participants could be presumed to use the same such criterion. Importantly, it also ensured that the pursuit of replication could be achieved by using the same measures as were used in the preceding work we sought to asses for reproducibility ^86,87^.

### Statistics

Our hypotheses focused upon identifying three different processes of episodic memory: recollection, item familiarity, and context familiarity. We hypothesized that recollection would exhibit both an FN400 for item-familiarity and an LPC for recollection but would not exhibit a late broad central negativity (BCN) effect, whereas context familiarity would exhibit only an BCN and neither an FN400 or LPC, and that item-familiarity would be associated with only an FN400 effect but not an LPC nor BCN effect. As the current investigation was based upon clear a priori-defined hypotheses derived from prior findings ^1^, analyses utilized planned paired t-tests to assess differences between the targeted conditions (two-tailed unless indicated otherwise) with an alpha level of .05. In cases where exploratory analyses were conducted to explore unplanned comparisons, ANOVA was used to qualify potential differences that may exist and were corrected with Geisser-Greenhouse corrections when necessary. Analyses conducted to assess the extent to which prior results could be found to replicate in independent data sets and/or were principally based upon a priori predictions that effects would differ in one way (i.e.: context familiarity exhibiting more negative-going ERP amplitudes than correct rejections, item-familiarity ^1^); such analyses thus applied targeted directional hypotheses utilizing one-tailed t-tests where noted in order to maintain a properly principled statistical approach, though most were also still significant with non-directional two-tailed tests as well (i.e.: multiply the p value by 2).

In reporting results of behavioral response times, we followed convention in the field to report the mean values of reaction times. When outlier data points were removed from the mean values (N = 2, using criteria of exceeding 3 standard deviations from the mean) the results from the analysis of mean response times remained the same. In this case, the two outlier subjects had longer mean RT values, merely indicating that some subjects sometimes responded a few seconds slower on uncertain memory conditions, which is common knowledge of reasonable variability in cognitive capabilities during conditions of uncertainty. Also, these inclusive data points are the aggregate of multiple responses for a person instead of a singular outlying value that might otherwise be considered as potential artifacts. Removing outliers based upon their post-hoc values would thus be inappropriately biasing the data based upon post hoc visual inspection of it. Therefore, we report mean reaction times with outlier data points included, since they represent the real average response times of subjects.

Because a non-significant p-value in conventional frequentist t-test methods is unable to inform whether there is actually evidence for the null hypothesis or if there is merely not sufficient evidence for any conclusion at all, analyses that revealed a null finding were subsequently quantified using Bayes factor analysis. Bayes factor analysis is a tool that allows researchers to quantify the relative strength of evidence for the null hypothesis (invariance) ^243–253^. A resulting Bayes factor value can generally be considered as the ratio of how likely a hypothesis is as compared to the likelihood to another hypothesis (for specific details see ^247,250^). For interpreting Bayes factor result values, it is conventionally viewed that a Bayes factor (BF_01_, evidence for the null hypothesis) of 0-1 typically represents there being no evidence available for the null hypothesis, 1-3 typically represents anecdotal evidence for the null hypothesis, a BF of 3-10 represents relatively moderate evidence for the null hypothesis, and a BF of 10-30 represents relatively strong evidence for the null hypothesis ^247,248,253^. Thus, Bayes Factor results are reported as such for the pertinent analyses as representing the relative strength of evidence for the null finding as noted above (BF_01_). The Bayes factor analyses were computed for paired or one-sample t-test designs. Results were calculated using the standard scale of r = 0.707 and the resulting outputs are provided as values in favor of the null hypothesis using the recommended Jeffrey-Zellner-Siow Prior (JZS, Cauchy distribution on effect size) ^250^.

### Reproducibility

The reproducibility of the effects of context familiarity was assessed in several ways through the present investigation. First, the present investigation (Experiment 1) was an assessment of the extent to which previously-reported effects ^1,39,67^ of context familiarity originally differentiated from the neural correlates of recollection in a sample of N = 25 could be found to be reproducible amid a larger sample (N = 54) that had been previously published in the domain of memory metacognition ^65,66^. Those original effects replicated in entirety in the present data. The sample of data of the present study thus demonstrated sensitivity of effect sizes and statistical power to detect the ERP effects of context familiarity originally discovered by the original study ^1^ by virtue of it having reproduced those findings of differences between context familiarity and recollection in a sample (N = 54) ^65^ that was more than double the original sample size (N = 25) ^1^.

Second, when the present findings (Experiment 1) then discovered that the neural correlates of context familiarity were dissociable from those for item familiarity, we sought to assess the extent to which these findings would have been observed in the original study ^1^ if those authors had only thought to look (they hadn’t). These effects of the present investigation were also found to be replicated in entirety (Experiment 2), both within-subjects and between-subjects, and across the same control analyses. Third, we also sought the extent of reproducibility of the findings across a range of several control analyses conducted in both Experiments 1 and 2, which systematically varied different conditions to assess potential confounds and possible alternative accounts. The control analyses found that the results remained reproducible even when reducing signal to noise ratio of trials and when reducing statistical power limited to only low levels of familiarity strength (excluding trials of strong familiarity responses).

Fourth, as the present study was a secondary analysis of existing independent datasets that had already been published with sufficient power and sample size to detect differences in previously published papers ^1,39,65,66^, it was not possible nor necessary ^254–258^ to perform an a priori power analyses for the present investigation since the sample size was already immutably established before the present work’s secondary investigation of analyses began. Similarly, widespread agreement in the literature that post-hoc power analyses would be both inappropriate and uninformative ^254,257^ ^256^ precluded their use here. The present results were fully replicated in independent data, and in twice the sample size of previously published results that the present investigation also replicated; thus, the sufficiency for statistical power in the study is self-evident.

## Key Resources Table

### Resource Availability

#### Lead Contact

Requests for further information and resources should be directed to and will be fulfilled by the lead contact, Richard J. Addante, PhD.

#### Materials Availability

- All materials and code used to generate and run the present experiment is available as a downloadable zip folder at https://github.com/IMAP-Lab/IMAP-Lab-IMAP-Implicit-Memory- Automated-Pipeline/blob/main/ItemSourceMemoryERP_Experiment_In_Presentation_AddanteLab.zip.
- Peer reviews and author responses from several rounds of reviews are available on Biorxiv at https://www.biorxiv.org/content/10.1101/2024.07.15.603640v5.supplementary-material.
- This paper analyzes existing, publicly available data, accessible via the previously published study using the same data^66^, freely available under the terms of the GNU General Public License at https://github.com/IMAP-Lab/IMAP-Lab-IMAP-Implicit-Memory-Automated- Pipeline.
- This paper does not report original code.

## Footnotes

1 Authors thank an Anonymous Reviewer for suggesting this valuable control analysis.

2 Authors thank an anonymous Reviewer for suggesting this valuable analysis

3 Authors thank an Anonymous Reviewer for suggesting this valuable control analysis.

4 Authors thank an Anonymous Reviewer for suggesting this valuable control analysis.

5 Similar results were also found when assessing the time windows used in the present study (Figure 3) of 400 to 600 ms, 600 to 900 ms, and 800 to 1200 ms: with main effects of memory F(1, 55) = 5.144, p = .027, *n*^2^ =.086, site (F(1.68, 55) = 7.70, p = .002, *n*^2^ = .123), time (F1.45, 55) = 44.797, p < .001, *n*^2^ = .449), and significant interactions of memory x time (F(1.41, 55) = 26.34, p <.001, *n*^2^ =.324), site x time (F1.56, 55) = 25.52, p <.001, *n*^2^ = .317), and a 3-way interaction of memory x site x time (F2.78, 55) = 4.66, p = .004, *n*^2^ = .078).

6 Authors thank a previous Reviewer for suggesting this possibility.

7 Authors thank two Reviewers for suggesting this possibility

8 Authors acknowledge a previous Reviewer for suggesting consideration of this alternative approach.

