## Supplementary material for "A third kind of episodic memory: Context familiarity is distinct from item familiarity and recollection": 4th Response to two (new) Reviewers

**Reviewer #2: In the manuscript, the authors observed distinct behavioral and ERP patterns across the three experimental conditions and claimed the presence of a third memory retrieval process. Upon evaluating the authors' revision in response to the reviewers' comments, I believe that the concerns raised in the previous rounds have been satisfactorily addressed. Therefore, I recommend accepting this manuscript for publication**.

> We thank the Reviewer for the time, effort, and consideration of the manuscript, in addition to acknowledging the satisfactory addressing of the concerns expressed by another pervious reviewer at *Nature Communications*.

**Reviewer #1: Review of iScience-D-24-11132**

**This article has been through several rounds of reviews at another journal. I was not a reviewer for the previous manuscript. I was asked to examine the reviews and the manuscript to help the *iScience* editors to reach a decision on the submitted manuscript.**

**Comments on the previous reviews: The previous reviews were very dense, but I read the concerns and responses carefully. The previous reviewers raised important issues. One key issue concerned the possibility that the central-negative ERP (called the BCN by the authors) reflected delayed recollection. The authors responded effectively to this concern, but I have to say, I don't understand the logic of this concern. Recollection produces a specific pattern of ERP activity (old ERPs are more positive than new ERPs), whereas the effect described in this paper is the opposite polarity (new more positive than old). The reviewers never suggest a reasonable explanation as to why the polarity of this effect would be reversed. It doesn't seem at all logical to me to look at that pattern and conclude that recollection was delayed.**

**A second reviewer issue concerned the possibility that the BCN is nothing more than another version of the Late Posterior Negativity (LPN). I, as one who published several LPN articles, agree with the authors that the BCN does NOT fit the LPN pattern. Even if the reviewers are correct, then the BCN is a unique LPN case. Either scenario begs for additional studies; therefore, this issue will need to be sorted out in the literature. In other words, It is the type of issue that should spark some debate and additional research; rather than being an issue that should prevent publication. Thus, I agree with the authors on this point as well.**

**The last major issue raised concerned the statistical analyses because some were based on a small number of observations. The previous editor sent the paper to a statistical expert and the expert only describes issues with the post-hoc power analysis and small sample sizes that plague all Psychological and ERP research. My determination is that the manuscript is reasonably free of statistical errors, which is about the best that any of us can do at any point in time with one research paper. In my opinion, the authors addressed all the reviewer concerns, and I don't understand why the article was not accepted for publication**.

> We thank the Reviewer for the fair assessment of the extensive prior sets of Reviews from *Nature Communications: Psychology,* in addition to acknowledging the publication merits in its previous editorial decision at *Nature Communications.*

**Evaluation of the manuscript:**
**As I said in the previous section, there has already been extensive feedback provided to the authors and the paper could be published in the present form. However, I provide some feedback in an attempt to help the authors improve the contribution of the work.**

> We thank the Reviewer for the thoughtful feedback and constructive comments, which response to have now bee incorporated into an updated, revised manuscript that has been improved thanks to the Reviewer's insights and suggestions.

**1. In general, I found the paper to be very dense. This is partly due to the fact that ERPs are the dependent measures and the novel contributions rely on MANY detailed analyses. However, some of the passages are dense because of the wording rather than the subject matter. I encourage the authors to identify and rewrite these sections. This paper has the potential to be very useful to memory researchers who are not ERP experts and students of all kinds. Reducing the cognitive load by using plain, simple text wherever possible will improve the accessibility of the work. Let me provide one example to illustrate my point**.

> The Reviewer makes a great point, and the manuscript has now been revised with an effort to simplify the text wherever possible, to improve the accessibility of the work.

**2. In section "5.4.3. The late parietal…", the authors write: "*However, 'similar' does not mean 'same', and upon closer inspection there are several critical distinctions between the present findings that differentiate them on orthogonal vectors of sensitivity to different memory conditions, reported scalp locations, and timing. Thus, there are a series of reasons why the LPN interpretation remains problematic and difficult to reconcile with the available data. Because the general similarities of the two patterns of effects can seem ambiguous at first, we provide a thorough address to disambiguate the two below*." More simple, concise text such as: "There are several reasons, described fully below, why the LPN interpretation does not fit the observed patterns." can get the point across without adding to the cognitive load**.

> We agree with the Reviewer's point and have simplified the text to now read as was suggested by the Reviewer. The revised text now reads:
>
> > "*However, upon closer inspection there are several reasons, described fully below, of why the LPN interpretation does not fit the observed patterns in the present data sets. Thus, the LPN interpretation remains problematic and difficult to reconcile.*"

**3. The text in section "2.6.1. Statistics" mentions RT analyses conducted on median RT values. Where are these analyses (in the supplementary info)? This could be helpful to support the authors' claim if it is consistent with the other RT analyses**.

> This reference in the Method section to an analysis of median response times was an inadvertent error of inclusion that had been a carryover from an earlier draft of the manuscript, but was found to be unnecessarily distracting from the primary findings being reported and should have been removed. The justification for using the industry standard analysis of average response times remains included in the text of the Method section.

We had determined that a median analysis of response times, while providing the same basic findings as the analysis of means, is not standard practice in the field for handling response time data, so had little to compare to for reference to existing literature findings. It thus was found to offer relatively little benefit while adding further to the unnecessary ‘denseness’ of the paper’s already extensive analyses. We meant to remove it from the manuscript long ago but was an oversight that has now been corrected in the updated revision.

That being said, if it may help the Reviewer to know, the results of the median RT analysis are included here for demonstration that the results remained the same as that of the primary mean RT results that had originally motivated them:

> *Response times for the three memory conditions were subjected to a one-way repeated measures ANOVA with three levels and Greenhouse-Geisser correction (Mauchly’s W = .79, p = .014), which revealed a significant main effect of condition (F(1.65,61.13) = 22.23, p < .001, n2p = .38). Post hoc pairwise comparisons using the LSD correction revealed that responses significantly differed for item familiarity and context familiarity, such that context familiarity was responded to reliably slower than item familiarity (MD = 326.08 msec, p = .033, 95% CI [27.69,624.48], as well as recollection (MD = 852.26 msec, p < .001, 95% CI [572.51,1132.02]. Item familiarity responses were also slower than recollection (MD = 526.18, p < .001, 95% CI [332.34,720.03]. Within source memory judgments, response times for the three memory conditions revealed a significant main effect of condition (F(2,74) = 4.17, p = .019, n2p = .10). Further investigation using post hoc tests with a LSD correction revealed that context familiarity responses were faster than responses for both recollection and item familiarity (MD = 406.24 msec, p = .002, 95% CI [158.75,653.73; MD = 349.93, p = .039, 95% CI [17.96, 681.89]. Response times during source memory for recollection and item familiarity conditions did not differ (MD = 56.32 msec, p = .738, 95% CI [-281.97,394.61].*

**4. In section "2.6.2 Reproducibility.", the last paragraph seems overly defensive and confusing because it describes a debate between reviewers and the authors. I recommend (1) removing the paragraph or (2) take the points in the paragraph and revise them into a footnote**.

The Reviewer makes an excellent point here, and in the revised manuscript we have removed that section of details, and replaced it with a simpler accounting excerpted in the text below:

> *“Fourth, as the present study was a secondary analysis of existing independent datasets that had already been published with sufficient power and sample size to detect differences in previously published papers (Addante et al., 2023; Addante, Ranganath, Olichney, et al., 2012; Addante, Ranganath, & Yonelinas, 2012; Muller et al., 2021), it was not possible nor necessary (Heinsberg & Weeks, 2022; Hoenig & Heisey, 2001; Lakens, 2022; Levine & Ensom, 2001; Zhang et al., 2019) to perform an a priori power analyses for the present investigation since the sample size was already immutably established before the present work’s secondary investigation of analyses began. Similarly, widespread agreement in the literature that post-hoc power analyses would be both inappropriate and uninformative (Heinsberg & Weeks, 2022; Hoenig & Heisey,*

*2001) (Zhang et al., 2019) precluded their use here. The present results were fully replicated in independent data, and in twice the sample size of previously published results that the present investigation also replicated; thus, the sufficiency for statistical power in the study is self-evident."*

**5. The descriptions of the samples could be cut (because they are described in other published papers) if space becomes a premium**.

This is a good suggestion that we would like to take, but we have kept this section as-is because the Method section does not contribute to the length parameters per this journal's policy, and because the journal's policies for STAR methods reporting require including the details of the descriptions of the samples of human participants.

**6. Section "3.2 Response times…." I suggest the authors clarify how they measured RTs. I assume that time 0 is the onset of the stimulus and the first measure is the time to make a recognition judgment. The source response time is less clear. Is that from time 0 or from the recognition response? That should be clarified. Also in this section, there is potential confusion between the type of response and type of item being contrasted. I am not sure what can be done except maybe use "source responses", "recollected items", "familiar items" or similar labels to draw a sharper distinction between response and item**.

Response times for item recognition and source memory responses were measured as follows: item recognition times were measured from the onset of the memory probe (i.e. the word) to the button press of the recognition decision; source memory response time was measured from the onset of the source memory prompt (which happened immediately upon the end of the item recognition judgment) to the button press of the source memory decision.  This information has now been added to the Method section: Conditions of Interest. Additionally, the section has also now been revised to draw a sharper distinction between information of response type and item, via adding clarifications of source memory "judgments" among descriptors for response times of the three different memory response conditions, and also adding clarification of the specific confidence numbers that were/weren't included in the various control analyses.

**7. Section "Experiment 1: Verifying if recollection…." the last line is missing a ")"**

This has been now been corrected.

**8. Section "Experiment 1: Verifying if recollection…." second paragraph - I suggest presenting the results in the same order as Figure 4 to make the results easier to follow.**

The reason why the results for this section are reported in the sequence that they are reported in (ERPs of context familiarity vs. ERPs of correct rejections, from 800-1200 ms for the BCN, then 600-900 ms for the LPC, then 400-600 ms of the FN400), is because that sequence is what follows from the direct hypotheses that guided our work's goal of assessing the primary findings of the 2012 paper (Neuroimage), which identified independent ERP effects later in the epoch for the BCN of context familiarity, and which were dissociated from recollection (600-900 ms LPC); the FN400 time window analysis was a confirmatory one that was ancillary to the primary question of replication, and thus it is reported thereafter the two main ones that begin the section.

**9. When reporting the Bayes Factors I suggest being clear that they are BF01 values rather than BF10 values, which would be interpreted differently**.

The Reviewer makes a good point here, and the manuscript has been updated accordingly in the text of the Results section. The Method section of the paper (in the subheading of "Statistics") also states the following:

> "*For interpreting Bayes factor result values, it is conventionally viewed that a Bayes factor ($BF_{01}$, evidence for the null hypothesis) of 0-1 typically represents there being no evidence available for the null hypothesis, 1-3 typically represents anecdotal evidence for the null hypothesis, a BF of 3-10 represents relatively moderate evidence for the null hypothesis, and a BF of 10-30 represents relatively strong evidence for the null hypothesis (Jarosz & Wiley, 2014; Jeffreys, 1998; Schönbrodt & Wagenmakers, 2018). Thus, Bayes Factor results are reported as such for the pertinent analyses as representing the relative strength of evidence for the null finding as noted above ($BF_{01}$).*"

**10. Section "4.0 Results" - it is somewhat circular argumentation to use data that had the effect to confirm that it exists. Can the authors be more specific about what data are used to establish the effect and what data are used to replicate the effect?**

The Reviewer raises an excellent point here, which we can certainly appreciate could be seen as being circular. However, we can assure that the analyses contained in Experiment 2 were definitely not circular.

The data from Experiment 2 came from previously published data sets in prior papers (Addante et al., 2011, PNAS; Addante et al., 2012a, Neuroimage; and Addante et al., 2012b, Neuropsychologia). Those prior papers published different findings using different comparisons of conditions, and including different total numbers of participants, than what is reported in the present study's utilization of those prior data sets in the current analyses, as detailed in the itemized points below:

1. The primary analysis of **the 2012a and 2011 data was comparing the conditions of 'context familiarity' vs. 'correct rejections'**.
    a. That analysis was not performed in the 2012b data, which instead identified a BCN effect via the contrast of source correct vs source incorrect responses.
    b. Neither of the three prior studies performed an analysis that directly contrasted the conditions of context familiarity vs. item familiarity, as we defined them here in the present study's Method section (and reported in the present study's Experiment 1).
    c. Of the analyses that were performed in these prior studies, they were only within-subject analyses, whereas we now also offered novel analysis of between-subjects comparisons.

2. The primary analysis of the present study focused, instead, upon **direct analyses between the conditions of 'context familiarity' vs. 'item familiarity'** (which had not been assessed in any earlier study).
    a. <u>Note A:</u> the present study's comparisons of context familiarity to correct rejections (Experiment 1) were performed merely to

assess the extent of replicability to those prior findings' that had focused upon recollection, e.g.: a proof of concept for reproducibility before engaging in fundamentally new analyses.

b. Note B: the present study's comparison of context familiarity to correct rejections in Experiment 2 aggregated across all three prior data sets (2011, 2012a, 2012b)- not the two data sets as were included in the 2012a paper, and thus constitute new data since 2012b data had not been included, nor had it even performed that same contrast in those participants, for instance.

c. Note C: The analysis of the prior data in the present study also included between-subject analyses that were not performed in the pervious publications.

**In summary, there are two different kinds of replications that occurred in the present work:**

> **The First Replication: Exp. 1 directly replicates a physiological discovery reported earlier** (of ERP differences between context familiarity vs. correct rejections; Addante, Ranganath, & Yonelinas, 2012, *Neuroimage*), using a sample more than double the original size (N = 54 vs. N = 25).
>
> **The present investigation then performed entirely novel analyses that made fundamentally new discoveries**: namely, that ERPs for context familiarity are reliably dissociable from ERPs for item familiarity, both within-subjects and between-subjects.
>
> **The Second Replication: these novel findings from the present investigation (Exp. 1) were then found to also be independently replicable among the other published public data sets** (N = 56; Addante et al., 2011, *PNAS*; Addante et al., 2012a, *Neuroimage*; Addante et al., 2012b, *Neuropsychologia*), referred to here as Exp. 2.

Thus, the analyses cannot be circular if they are performing different analyses on different conditions being compared, and among a non-identical group of participants included in the analyses. The same is stated as such in the manuscript, in the Method subsection on Reproducibility, where it still states:

> "…*when the present findings (Experiment 1) then discovered that the neural correlates of context familiarity were dissociable from those for item familiarity, we sought to assess the extent to which these findings would have been observed in the original study if those authors had only thought to look (they hadn't).*"
>
> and also in the Participant section of the Method, where it states: "*The participants included in Experiment 2 were aggregated across three previously published studies that used nearly identical experimental protocols of item recognition confidence followed by source memory judgments as were used in the present Experiment 1, for a total sample of N = 56. These studies had only*

*directly contrasted conditions of recollection and context familiarity, but had never directly assessed if the ERPs for item- and context-familiarity differed*.*"*

**11. In section "4.3.2.", the use of parentheses is confusing in this text: "(the UC Davis Dynamic Memory Lab (2008-2010, majority Caucasian) vs. California State University - San Bernardino (2019, majority Hispanic), using different EEG hardware**."

The updated manuscript has been revised accordingly and has removed the text in parentheses to make it read more straightforwardly.

**12. Section "4.3.3." typographical error (extra period at the end of the sentence) "...thus the main finding was replicated again and with a larger sample than original finding in Experiment 1**."

The updated manuscript has been revised accordingly and has removed the "."

**13. Section "5.1." This paragraph could be better organized so that it ends with the models that are being advanced. In other words, funnel the evidence to the conclusion. Also, articulate the "established norms" that are referenced here. Otherwise, the statement is vague and confusing**.

The Reviewer makes a great point, and the updated manuscript has now been revised to articulate what the established norms are for using ERP findings to deduce distinct cognitive operations, and re-organized this paragraph to funnel the findings towards an overall conclusion. The paragraph from section 5.1 now ends with the following text:

"*Item- and context-familiarity thus appear to be operating as fundamentally different kinds of memory processes, based upon established norms for identifying cognitive processes via neuroimaging and behavioral data (Duzel et al., 2001; Duzel et al., 1997; Henson, 2005; Leynes et al., 1998; Mecklinger & Jäger, 2009; Rugg, 2009; Rugg & Coles, 1995; Rugg, Mark, et al., 1998; Shallice, 1988; Simons, 2009; Yonelinas, 2009), such that dissociable physiological activity occurring at different times, at different locations, and for different conditions, cannot be taken as evidence that they are reflecting the same cognitive process and must therefore reflect at least partially non-overlapping cognitive operations. Together, this provides strong evidence that these three observations of ERP effects of memory are reflecting mutually exclusive, distinct episodic memory processes, and therefore advancing an updated model of the organization of episodic memory as discussed in further detail across each finding throughout the sections below."*

**14. Section "5.2" the sentence: "There are a few ways to view these patterns of results : (1) They are consistent with existing findings of the relative speeds of recollection and familiarity (recollection being slower and familiarity being faster, 32,53; (2) they are contrary to traditionally-held patterns of recollection being slower than familiarity." These statements are contradictory; therefore, this became difficult to follow. I suggest revising this segue to the discussions that follow**.

We agree that this section was worded poorly. The updated manuscript has now been revised to more clearly communicate the intended point, stating that:

"*There are two different ways that one can interpret these patterns of behavioral results*".

**15. Section "5.2" the authors write "Moreover, recent studies noted below have reported recollection to have faster responses than familiarity during recognition, suggesting that the traditionally assumed speeds of recollection and familiarity should not necessarily always be taken as simply 1-directional (i.e.: Familiarity > Recollection)..." Do the authors mean familiarity < [faster than] recollection here**?

Yes, we did mean that familiarity was faster than recollection, but we understand the reviewer's point that it was worded in a confusing way, and we have now updated that sentence of the Discussion to read more clearly as:

"*Moreover, recent studies noted below have reported recollection to have faster responses than familiarity during recognition, suggesting that the traditionally assumed speeds of recollection and familiarity should not necessarily always be taken as simply 1-directional (i.e.: that familiarity must always be faster than recollection) and that is likely more complex in directionality*."

**16. Section "5.4.0" the title "Alternatives, Limitations, and Future Directions" was a bit misleading because there is a separate future directions section. A better title would be helpful here**.

This section has now been re-labeled as "***Possible Alternative Interpretations and Limitations."***

**17. Section "5.4.1" did not appear to be very cohesive. I suggest separating some of the arguments. The authors could deal with the topography of the FN400 as a limitation and separate the unitization argument. That would improve the section**.

The Reviewer makes a good point again here. We have now revised the section to be simpler and read more cohesively, following the Reviewer's suggestion to separate the topics of the FN400 topography and the unitization argument, which now are presented in their own separate paragraphs.

**18. Section "5.4.2." I suggest emphasizing the polarity difference of the BCN because it is strong evidence against a delayed recollection argument**.

We thank the Reviewer for making a good suggestion here. We have now updated the Discussion section of the revised manuscript to emphasize the polarity difference of the BCN effect from the LPC effect. The updated version now reads as follows:

"*The term of 'delayed recollection' appears only sparsely in the memory literature. One of the few extant studies of "delayed recollection" using ERPs (Liesefeld et al., 2016) found that 'delayed recollection' occurred under conditions of "changed viewing conditions"- which were not a part of, nor directly applicable, to the present study. Notably, that study did not find any negative-going ERP effects to be associated with their condition of 'delayed recollection', which was instead associated with the traditional, positive LPC measures (the opposite of the present findings). Since recollection typically produces a specific hallmark pattern of positive ERP effects (Curran, 2000; Friedman, 2013; Kwon et al.,*

*2023; Rugg & Curran, 2007), and we instead observed a distinct opposite polarity of negative-going ERP effects, the logic to support a 'delayed recollection' account remains lacking. That is, there is not a reasonable explanation for why the polarity of the canonical recollection-related physiology would have been reversed among our other control analyses demonstrating the standard positive-polarity of recollection effects in all other standard conditions of memory. This provides an additional line of evidence against a 'delayed recollection' account and it is thus difficult to support as a viable account of our present findings."*

**19. Section "5.4.2" a typo in "Why then would it now be something different and special for this one condition of accurately retrieving the source- especially especially…" [redundant word**]

The revised manuscript has now been updated to correct this typo.

**20. Section "5.4.3" needs a better title to match the others. Maybe "Is the BCN a case of the LPN?"**

This section subheading has now been revised to read: "*Is the BCN a case of the Late Parietal Negativity (LPN)?"*

**21. Section "5.5" typo in "consistency to the literature of known EPR effects of episodic…" [should be ERP**]

The revised manuscript has now been updated to correct this typo.

**22. Section "5.6" this claim "(Figure 8), which is unique in being specifically physiologically-informed, as prior models of memory have instead been either informed by cognitive psychology or neuroanatomical/neuroimaging studies…" is a little overstated because Mecklinger and Bader (2020) used physiological data. I think there are others that incorporate it also. I recommend softening this claim**.

The Reviewer makes a good point, and we have now revised the manuscript to soften that claim, and to also acknowledge the model of Mecklinger & Bader (2020), though noting that while that model only focused upon specific episodic memory subprocesses of familiarity (absolute and relative familiarity), our model proposed in the present manuscript is broader than that and encompasses other episodic memory processes of recollection, as well as represents a fuller suite of long term memory processes. The updated text in our Discussion is as follows:

"*We propose an updated model to conceptualize the organization of specific episodic memory processes among a broader framework of long term memory (Figure 8), which is specifically physiologically-informed, whereas most prior models of long term memory have instead been either informed by cognitive psychology or neuroanatomical/neuroimaging studies (Cohen & Eichenbaum, 1993; Diana et al., 2007; Eichenbaum et al., 2007; Milner et al., 1998; Squire, 2009; Squire et al., 2007; Squire & Zola, 1997; Wixted & Squire, 2010, 2011b; Yonelinas et al., 2019), though see Mecklinger & Bader (2020) for an example of a physiologically informed model of familiarity-specific memory processing*."

**23. Section "5.6" I found a lot of this text to be highly speculative and prone to people misunderstanding the authors' points by thinking that recognition works this way in every case. I suggest revising this section to focus on the model (BIC) that is being tested. In other words, come back to the model that motivated the work**.

> The Reviewer makes a great observation and suggestion here, and we have incorporated it into a revised text of the manuscript’s Discussion for that section. In this effort, we made clear that the proposed model was based upon a motivation from the BIC model, and emphasized that we do not suggest that recognition works this way in all cases, with excerpted sections as noted below:
>
> > “*Together with the present studies, these findings converge to suggest an updated framework for the organization of long-term episodic memory from the canonical declarative memory model used ubiquitously throughout textbooks, following from the BIC model that had originally motivated the hypotheses tested in the present work (Diana et al., 2007). We propose an updated model to conceptualize the organization of specific episodic memory processes among a broader framework of long term memory (Figure 11)*...” And later follow with the statement that:
> >
> > “*How might the subjective experience of context familiarity occur during cognition? We draw upon our proposed additions to the BIC model/variants (Diana et al., 2007; Eichenbaum et al., 2007; Yonelinas et al., 2019) to suggest a cognitive framework that can be built to understand how the condition of context familiarity may arise or manifest (note: this is a proposal of how it might occur in some instances but should not be taken as suggesting that it is how recognition works in every case).”*

**24. In Figure 3, describe what the white boxes indicate**.

> The manuscript has now been updated to include a description of what the white boxes denote in Figure 3 (they denote the sites from the topographic maps that are then shown for the corresponding ERP figures beneath the maps, as well as statistical analyses).

**25. Figure 5 has "N = 11 with n > 11" This is very confusing. I think they are referring to the signal-to-noise ratio. Maybe add "trials" to have "n > 11 trials" in the figure**?

> This figure has now been updated accordingly, thank you for the good suggestion.

**26. Figure 11 - "Absolute/Relative" appear under fluency when they are forms of "Familiarity**".

> Thank you for pointing this out. This figure has now been updated accordingly, with those elements removed from the depiction.
