## Supplementary material for "A third kind of episodic memory: Context familiarity is distinct from item familiarity and recollection": Third Response to Reviewers at NatureCommsPsych

September 1, 2024

Dear Editor,

Hello, please consider our transferred manuscript, "*A third process of episodic memory: Context familiarity is distinct from item familiarity and recollection"* for publication in *Nature Communications*. In short, it reports the discovery of a new process of human episodic memory, which we call 'context familiarity'. Like the discovery of a new animal species, the discovery of a new process in human cognition is quite noteworthy; we report a new one. <u>This is an ideal study for the Replication Collection for three reasons</u>:

**First, it directly replicates a physiological discovery reported earlier** (Addante, Ranganath, & Yonelinas, 2012, *Neuroimage*), while using existing data from the published literature (Addante et al., 2023, *Psychophysiology;* Muller, et al., 2021, *European Journal of Neuroscience*) to perform a secondary analysis aimed to directly test the extent of replication. That finding focused upon ERPs for memory of context being dissociable from that of recollection, and here we found those results to be fully reproducible using a sample more than double the original size (N = 54 vs. N = 25).

**Second, the present investigation then performed entirely novel analyses that makes fundamentally new discoveries**: namely, that familiarity of context is reliably dissociable from familiarity of items in episodic memory- and furthermore, that these two kinds of memory were evident as dissociable among multiple different physiological measures as well as multiple different behavioral measures, both within-subjects and between-subjects. These findings persisted even when subjected to the most stringent control analyses designed to directly test and falsify the findings using rigorous statistic approaches including Bayes Factor analyses to quantify meaningful null findings, too.

**Third, these novel findings from the present investigation were then found to also be independently replicable among four other published public data sets** (N = 56; Addante et al., 2011, *PNAS*; Addante et al., 2012a, *Neuroimage*; Addante et al., 2012b, *Neuropsychologia*; Addante, 2015, *Neuroimage*). These findings wholly reproduced results from the primary investigation (Experiment 1): for both within-subjects and between-subject analyses and including every rigorous control analysis that were specifically designed to falsify the hypotheses. These findings cannot be accounted for by existing theoretical models of memory, and thus constitute the discovery of a new model of memory processes which we propose in the manuscript, in which there are three (instead of two) processes of human episodic memory: recollection, item familiarity, and context familiarity. This constitutes a major discovery.

**As such, be believe it is an ideal fit for the Collection on Replication,** and serve as an exemplar of rigorous efforts in science to ensure validity and reproducibility of findings that yields important new insights into the understanding of the human mind and brain. These conclusions are critically based upon core efforts of reproducibility in science, derived from the results of rigorous replications among several independent data sets from different labs (UC Davis, CSU – San Bernardino) and diverse cohorts (Caucasian/Asian, Hispanic/Latino) collected across different decades (2009, 2018).

**The relevant history & context of our transfer within the Nature Springer system:**
This manuscript has already been subjected to extensive peer reviews in *Nature Communications: Psychology* that were mostly all positive. **Two of the three original Reviewers reported having all of their comments satisfied with our comprehensive revision, and they commended us on the excellent work**. The third reviewer first reported having "half" or her original comments satisfied, (e.g. providing 2.5 of 3 Reviewers being satisfied), following which we engaged in a documented series of extensive clarifications and cogent rebuttals to the lingering items she commented upon. We firmly believe that our prior response to reviewer comments in the process at NCP comprehensively demonstrated that her lingering comments were vanquished and unable to provide a viable account for our findings. That is, we showed convincingly that she misunderstood and was incorrect in the reviewer comments that were made, and that they should not preclude publication; we now stand by that position even more.

**A 4th Reviewer was solicited to assess the statistical approaches used.** That Reviewer reported back that our calculations were indeed correct. We thoroughly agreed with the Reviewer 4, though, that certain analyses were problematic (namely, post-hoc power analyses and post-hoc sensitivity analyses) and that they "represented misconceptions and a lacking understanding about statistics". ***The reason we agree 100% is because we did not want to do those incorrect analyses at all, but had them forced upon us in a coercive review process by both Reviewer 3 and then insisted upon by the Editor in Chief's several email letters to us.***

That is, when we did not want to do those incorrectly-suggested post-hoc analyses by Rev. 3, the Editor fully rejected the revised manuscript for specifically-stated reasoning that it did not perform the ill-conceived post-hoc statistical tests, and then she later reverted to accept the revision when we did reluctantly add those misconceived statistics to the revised manuscript. That, in turn, led the Editor to reject the paper (for doing the wrong statistics that she forced us to do). We believe that series of events is incorrect, inappropriate, and unethical in the spirit of open science integrity. Thus, for a manuscript in which all other Reviewer comments have been commended as satisfied and/or reasonably vanquished, and with no critical flaw having yet been found, we therefore believe that it presents a fair candidate for assessment at the related journal of *Nature Communications*- particularly due to its shared Collection on Replication.

**The manuscript has already been extensively Reviewed and Re-Reviewed by at least 4 Reviewers, and with positive recommendations to publish from the original 2.5 of 3 Reviewers. We have now revised the manuscript in alignment with Reviewer 4's comments to simply remove the problematic part of post-hoc power analyses** that Reviewer 3 had required, and removed the problematic part of post-hoc sensitivity analyses that the Editor had required, too. These are easy fixes and have greatly improved the manuscript. They are also the proper fixes and revert the results to the way that we had originally correctly-reported the findings. Within the newly revised manuscript, we note the following in regard to the issue raised by both Reviewer 3 and the Editor, and which was corrected by Reviewer 4:

> "*as the present study was a secondary analysis of existing independent datasets that had already been published with sufficient power and sample size to detect differences in previously published papers*[1-4]*, it was not possible nor necessary*[5-8] *to perform an a priori power analyses for the present investigation since the sample size was already immutably established before the secondary investigation of analyses began*. … There is widespread statistical agreement in the literature that such "*post-hoc analyses are not informative*"[8], are "*fundamentally flawed*"[5], are "*conceptually flawed and analytically misleading…and do not provide sensible results*"[7], and "*should not be used*" [8] since they are "*simply not informative for data interpretation*" [8]. Prior authors have clearly noted that: "*we encourage authors and peer-reviewers to avoid using or requesting post hoc power calculations*" [8].

Despite the core points from the literature cited above, we still had peer-reviewers and Editors at *NCP* request the same post-hoc power calculations that the experts state not to do, and it negatively affected the Editorial decision on our otherwise-positive manuscript. We believe that a better-informed Editorial discretion may be available in the present transfer of our manuscript. To aid in that discretion, we reiterate below the clear and consistent facts from the literature about the statistical measures that we had previously been rejected for having included on our earlier manuscript under Editorial demands:

RESEARCH ARTICLE

**Post hoc power is not informative**

Lacey W. Heinsberg, Daniel E. Weeks

First published: 01 June 2022 | https://doi.org/10.1002/gepi.22464 | Citations: 6

Read the full text › PDF TOOLS SHARE

#### Abstract

Post hoc power estimates are often requested by reviewers and/or performed by researchers after a study has been conducted. The purpose of this commentary is to provide a heuristic explanation of why post hoc power should not be used. To illustrate our point, we provide a detailed simulation study of two essentially identical research experiments hypothetically conducted in parallel at two separate universities. The simulation demonstrates that post hoc power calculations are misleading and simply not informative for data interpretation. As such, we encourage authors and peer-reviewers to avoid using or requesting post hoc power calculations.

**The abuse of power: The pervasive fallacy of power calculations for data ana...**
John M Hoenig; Dennis M Heisey
*The American Statistician;* Feb 2001; 55, 1; ABI/INFORM Global
pg. 19

### Statistical Practice

#### The Abuse of Power: The Pervasive Fallacy of Power Calculations for Data Analysis

John M. HOENIG and Dennis M. HEISEY

It is well known that statistical power calculations can be valuable in planning an experiment. There is also a large literature advocating that power calculations be made whenever one performs a statistical test of a hypothesis and one obtains a statistically nonsignificant result. Advocates of such post-experiment power calculations claim the calculations should be used to aid in the interpretation of the experimental results. This approach, which appears in various forms, is fundamentally flawed. We document that the problem is extensive and present arguments to demonstrate the flaw in the logic.

KEY WORDS: Bioequivalence testing; Burden of proof; Observed power; Retrospective power analysis; Statistical power; Type II error.

culations as a matter of policy (Anon. 1995; Anon. 1998). We emphasize that these calculations are sought primarily with the thought that they are useful for explaining the observed data, rather than for the purpose of planning some future experiment. We even found statistical textbooks that illustrate the flawed approach (e.g., Rosner 1990; Winer, Brown, and Michels 1991; Zar 1996). Researchers need to be made aware of the shortcomings of power calculations as data analytic tools and taught more appropriate methodology.

It is important to understand the motivation of applied scientists for using power analysis to interpret hypothesis tests with nonsignificant results. The traditional, widely accepted standard has been to protect the investigator from falsely concluding that some treatment has an effect when indeed it has none. However, there is increasing recognition that a "reversal of the usual scientific burden of proof" (e.g.,

#### INTRODUCTION

Power analysis plays a key role in designing and planning prospective studies. For clinical trials in biomedical and psychosocial research, power analysis provides critical information about sample sizes needed to detect statistically significant and clinically meaningful differences between different treatment groups. Power analysis also provides critical information for evaluating cost–benefit ratios so that studies can be conducted with minimal resources without compromising on scientific integrity and rigour.

What is interesting is that some journals also ask for power analysis for the study data that were already analysed and reported in a manuscript before considering its publication. Although the exact purposes of such requests are not clearly stated, it seems that this often happens when manuscripts include some non-significant results. As such post hoc power analysis is conceptually flawed, concerns have been raised over the years.[1–4] Despite these warnings, some journals continue to ask for such information and use it as part of their decision process for manuscript publications.

As most research studies are conducted based on a random sample from a study population of interest, results from power analysis become meaningless, as the random component in the study disappears once data are collected. Power analysis shows the

Check for updates

© Author(s) (or their employer(s)) 2019. Re-use permitted under CC BY-NC. No commercial re-use. See rights and permissions. Published by BMJ.

[1]Department of Family Medicine and Public Health, University of California System, Oakland, California, USA
[2]Naval Health Research Center, San Diego, California, USA
[3]Leidos, San Diego, California, USA

**Correspondence to**
Dr Xin M Tu, Family Medicine and Public Health, University of California San Diego, La Jolla, CA 92093, USA;



#### DISCUSSION

Power analysis is an indispensable component of planning clinical research studies. However, when used to indicate power for outcomes already observed, it is not only conceptually flawed but also analytically misleading. Our simulation results show that such power analyses do not indicate true power for detecting statistical significance, since post hoc power estimates are generally variable in the range of practical interest and can be very different from the true power.

In this report, we focus on the relatively simple statistical model for comparing two population means of continuous outcomes. The same considerations and conclusions also apply to non-continuous outcomes and more complex models such as regression. In general, post hoc power analyses do not provide sensible results.

**Contributors** All authors participated in the discussion of the statistical issues and worked together to develop this report. AR, RR, SR and XT discussed the problems with the justification of post hoc power analysis and interpretation of such power analysis results within the contexts of their studies and how to approach and clarify the issues in clinical research. YZ, RH and XT worked together to develop

The findings reported in our manuscript have important implications for understanding the core of human memory, of resolving differential results among other laboratories, and providing clinicians much more precision in measuring memory process impairments in neuropsychological patients. We believe it will be of broad interest to the public, too, as the breadth of its message resonates with people when a new kind of memory is discovered.

We are happy to suggest several well-qualified expert reviewers, including the following:

1. Dr. Lynn Nadel, PhD, University of Arizona (Expert on memory and context)
2. Dr. Steven J. Luck, PhD, University of California, Davis (Expert on ERP statistics)
3. Dr. Andrew Leynes, PhD, College of New Jersey (Expert on ERPs of memory)
4. Dr. Axel Mecklinger, PhD, University of Saarland (Expert on ERPs of memory)
5. Dr. Ali Mazaheri, PhD, University of Birmingham (UK) (Expert on ERP statistics)
6. Dr. Andrew Yonelinas, PhD, University of California, Davis (Expert on memory)
7. Dr. Ron Mangun, PhD, University of California, Davis (Expert on cognition ERPs)
8. Dr. Tim Curran, PhD, University of Colorado, Boulder (Expert on memory ERPs)
9. Dr. Arne Ekstrom, PhD, University of Arizona (Expert on memory, context, ERPs)

I sincerely appreciate any consideration that is available, and thank you for the time, effort and consideration.

Respectfully,

Richard J. Addante, Ph.D.
Associate Professor of Psychology, Biomedical Engineering
Director, EEG Research Laboratory
National Institute of Health, LRP Fellow

#### <u>Response to Reviewers, Round 3: *Nature Communications/ Psychology*</u>

**Reviewer #4:**

**I was specifically asked to comment on the power and sensitivity analyses, but will state that all in all the authors do make bold claims based on limited data**.

We thank the Reviewer for the time and expertise devoted to providing the astute, and fair, and correct Review. We completely agree that the manuscript does make bold claims, but those claims are only made considering rigorous methodology, comprehensive control analyses, and independent replications among our fairly large datasets relative to standard ERP studies of memory (detailed further below).

We disagree only with the subjective claim that there is 'limited data', and instead we are proud to have provided a relatively large set of data relative to the history of other published standard event-related potential (ERP) studies of memory, and particularly the historical ones of foundational findings detailed below [1-13]. In fact, we provide a substantially larger amount of ERP data (N = 54, N = 56) that is more than triple the size of the foundational ERP studies of memory, which have historically (and successfully) provided reliable findings that have stood the test of time despite samples ranging N = 12 to N = 16. Below, we enumerate the comparative size of major ERP studies that weren't deemed to be 'limited' data:

- Paller, & Kutas. (1992). *Journal of Cognitive Neuroscience*, **N = 12**.
- Wilding & Rugg. (1996). *Brain*. **Exp. 1: N = 18, Exp. 2: N = 19**.
- Duzel, et al.(1997). *Proceedings of National Academy of Sciences,* **N = 16**
- Rugg, M., et al. (1998). *Nature.* **Exp. 1: N = 16, Exp. 2: N = 18**.
- Rugg, M. et al. (1998). *Experimental Brain Research*. **N = 16.**
- Duarte, A., et al. (2004). *Cognitive Brain Research*, **N = 13**.
- Leynes, et al. (2005). *Consciousness & Cognition.* **Exp1 N = 24, N = 30**
- Woodruff, Hayama, & Rugg, M. (2006). *Brain Research*. **N = 16.**
- Curran & Hancock. (2007). *Neuroimage*. **N = 33**.[6,14,15]
- Gruber & Otten (2010), *Journal of Neuroscience*. **N = 24**
- Addante, et al. (2011). *PNAS*. **N = 17.**
- Addante, et al. (2012). *Neuroimage*. **Exp. 1 = N = 25, Exp. 2 = N = 22**.
- Addante, et al. (2012). *Neuropsychologia.* ***Exp. 1:* N = 6, Exp. 2: N = 3.**
- Leynes & Addante (2016). *Cog., Affect. & Behav. Neuro.* **N = 36; N = 33.**
- Ozubko, et al., (2021). *Cognitive, Affective & Behavioral Neuro.* **N = 40.**
- *Addante, et al.* ***(<u>Present Manuscript): Exp. 1: N = 54, Exp. 2: N = 56.</u>***

Furthermore, if one still considered our primary findings from Exp. 1 (N = 54) as 'limited', then it presupposes that the findings are in error (i.e. a false positive, or false negative, etc.) since the allegedly 'limited' data would be inherently insufficient for detecting reliable results (either null, or spurious findings). That concern is, and ought to be, ameliorated then by the full and complete replication of Exp. 1 results (N = 54) in an independent dataset (Exp. 2, N = 56).

Even further, the ERP data that we provided was accompanied by separate findings from behavioral data that was evident in response time differences in two different measures (item recognition, and source memory). Finally, the data itself is taken directly as secondary re-analyses from several different studies that are already published in the peer-reviewed literature, and which have been subjected to rigorous peer-review from those respectable publications, all of which determined that the data was more than merely 'limited' and was in fact quite sufficient for supporting the claims made by each prior publications- and, we contend here, the present as well.

**Regarding the technical aspects of the analyses, as far as I can tell, the described procedures were performed correctly. I am a bit surprised that the authors differentiate Cohen's d and dz in the text. As far as I can tell, both values they use are actually dz and thus they are comparing effect sizes that have been normalized using the same procedure. This is relevant since dz takes into account the correlation between two measures and thus using this estimate of the effect size will usually lead to higher estimates of power in an experiment without increasing the number of participants. However, since the comparison is consistent, I see no problem with this approach. My recalculation of a random example indicated that this procedure was applied correctly and I have understood exactly what the authors did.**

We thank the Reviewer for kindly acknowledging that our calculations were correct, and independently verified via the Reviewer's efforts to re-calculate.

**However, to answer the question whether the two procedures allow the authors to conclude that their study was sensitive to the effects they were trying to find, I would like to briefly explain, what, to my understanding, these procedures are doing.**

**1. The authors are computing an effect size point estimate for all significant tests. They then use this estimate to perform a post hoc power analysis that gives them a power value (1 – βerror, confusingly called β in the text). This power value is essentially meaningless, as the point estimate has been conditioned on there being a significant effect and thus the calculation will always indicate enough power (this is because t-values can be directly converted to dz). This becomes immediately clear, when considering the confidence interval around the point estimate, which contains values close to zero, indicating that the true effect size might be much smaller than the estimate. The power of a study is related to the true effect size and not the effects size estimate. The effect size estimate can, within limits, inform future studies, but cannot be used in this way within a study.**

**2. The authors compute the effects size that the study had 80% power to detect (the only thing changing the values here is the sample size of the subsamples used for each analysis). This is in itself not problematic and allows the reader to**

**estimate whether they think effect sizes in ERP research should be this large and thus whether to trust the present data. One caveat is that we are using dz here, which most researchers have little intuition for, since it takes into account the correlation of the two measures. In any case, comparing this effect size from the sensitivity analysis to the point estimate is meaningless for the same reason, as given in 1., since the point estimate is still conditioned on there being a significant effect**.

**As can be seen from this description, it is in my opinion not possible to conclude that the study in question was sensitive to the effects that the authors wanted to study. That is not to say that is was not, but merely that these procedures do not give any evidence for such a conclusion. Also, these analyses or any of the others do not exclude type 1 or 2 errors, which the authors seem to be constantly claiming, this is quite frankly not something that can be done using statistics and indicates to me some lack of understanding, how statistics work. As mentioned in the beginning from my reading of the entire manuscript, I do think that the authors are making bold claims based on relatively limited data. It seems that this strategy is extended to the power and sensitivity analyses, where they seemingly provide evidence for a robust finding, but it turns out that these claims are in fact based on a misconception of how effect size estimates work**.

> We very much appreciate the Reviewer's kind and detailed explanation of the flaws in the post-hoc analyses of power and sensitivity that was performed in our prior manuscript. Importantly*: we fully agree with the Reviewer on those points- especially the part about it "indicating a lack of understanding of how statistics work", and "based upon a misconception of how effect size estimates work"*. The revised manuscript has now been updated to have those 'misconceived' statistics removed, which did not affect any of the other data, results, or conclusions.
>
> *More importantly, though, we very much want to let the Reviewer know that the "misconception" and "lack of understanding was not ours": we initially refused to do those analyses but had them forced upon us the Editor in Chief of the journal* (Nature Communications: Psychology) as a coerced pre-requisite for our laboriously-revised manuscript being sent back out for a 3rd and final review.
>
> That is, when we submitted our invited re-submission for a 'revise & resubmit' following two of three Reviewers reporting being satisfied with no further comments, the Editor rejected it in-hand and them documented a long email letter in which she clearly elucidated the requirements to do the analyses that this Reviewer 4 now confirms "*indicate a lack of understanding how statistics work*" and are "*based upon a misconception of how effect sizes work*". When we reluctantly added those ill-conceived analyses back into the paper for an Appeal, the Editor in Chief accepted the appeal, approved the manuscript's statistics, and send it back out to review- where it was rejected for having done their forced test.

For full transparency, as evidence of this, below is a copy of that Editorial demand, effectively holding the paper ‘hostage’ to ill-conceived statistics:

Gmail

**Rick Addante <>**

**Decision on Communications Psychology manuscript COMMSPSYCHOL-23-0270C**

**** <> Thu, Apr 18, 2024 at 11:23 AM
Reply-To:
To:
Cc:

Dear Rick

Thank you for submitting your revised manuscript titled "A third process of episodic memory: Context familiarity is distinct from item familiarity and recollection". I have now had the opportunity to read your work and discuss it with the other editors in my team. I consulted in particular with Editorial Board Member Prof Anna-Lena Schubert, who serves on the board as an Editor for cognitive psychology and as a statistics expert.

When we request to strengthen the conclusions of a manuscript, the editors carefully assess whether appropriate revisions have indeed been incorporated into the revised manuscript before deciding whether to send it back to the referees.

After careful consideration and discussion with my colleague, I am sorry that we do not feel that the editorial concerns expressed in my previous emails are sufficiently addressed to justify sending this revision back to the reviewers.

In detail, we specified concerns about the strength of the statistics, which were also mentioned by previous referees. In the last decision letter, I requested a sensitivity analysis that would offer insight into how well-powered the study was to detect the smallest effect size of interest. I also specified that to achieve sufficient power, you would in all likelihood need to add additional data.

"In sum, we would welcome a revised manuscript that addresses all of Reviewer #3's concerns and especially their point about how process pure the conditions are. However, in keeping with the journal's focus on reproducibility and strength of evidence, the revision would need to present analyses that are sufficiently sensitive (>=80% power for the smallest effect size of interest) to address all existing concerns. This will likely require additional data. We would expect that the sensitivity analysis enclosed in the revision is generally reproducible, and that an expert in the field, i.e. a researcher familiar with EEG, would be able to immediately judge its soundness from the presentation in the paper."

The current version includes a power analysis but lacks a sensitivity analysis. Furthermore, upon reviewing all the statistics presented in the paper, we find insufficiently compelling evidence to support some of its main conclusions. While we noted that you helpfully provide Bayes Factors, these do not alleviate concerns about strength of evidence where they do not exceed a BF>3 (in favour of the H1) or BF<0.3 (where the interpretation is that an effect is absent).

For Communications Psychology, the strength of evidence and robustness of the data are key criteria. The concerns about the statistical strength of your work therefore preclude further consideration for publication in Communications Psychology and we decided against sending the work back to the referee.

This was Editorial demand excerpted above also was preceded by earlier Editorial demands to do the same, which we initially resisted (and for which we were rejected for having properly resisted), as documented for transparency in the excerpts below:

****
to me, Marike.schiffer, raddante

Thu, Mar 14, 8:15 AM

** Please ensure you delete the link to your author homepage in this e-mail if you wish to forward it to your coauthors **

Dear Rick,

Thank you for your patience during the peer-review process. Your manuscript titled "A third process of episodic memory: Context familiarity is distinct from item familiarity and recollection" has now been seen by 3 reviewers, whose comments are appended below.

Please let me apologize again for the unconscionable delay in returning to you with a decision. We invited Reviewers #1,#2,#3 from the first round to re-review your work and they all kindly agreed to do so. Unfortunately, not all of them were able to meet the usual turnaround times, which caused an initial delay.

As you will see below, Reviewer #3 voiced not only methodological and conceptual concerns that remain unresolved, but also noted the small sample size. You will also see that we obtained additional comments from Reviewer #1 on the most critical issues raised by Reviewer #3, a process we occasionally elicit when the reviewer reports are in conflict and we believe strongly that additional input will benefit the decision.

Editorially, we have significant concerns about the inclusion of analyses at less than 80% power for the target effect size. Some of your control analyses, including those that speak to some of the referees' concerns fall within that range. We invited a statistics expert to comment on the sensitivity analysis and power issue; as I mentioned earlier, because the first statistician who agreed to vet this aspect was ultimately unable to return a report, this added to the initial delay in our decision. The most recent delay is a result of another (second) statistician agreeing to vet the manuscript and subsequently withdrawing from the process; although they did not submit a report, they highlighted that they were unable to reproduce the sensitivity analysis and therefore unable to comment.

In sum, we would welcome a revised manuscript that addresses all of Reviewer #3's concerns and especially their point about how process pure the conditions are. However, in keeping with the journal's focus on reproducibility and strength of evidence, the revision would need to present analyses that are sufficiently sensitive (>=80% power for the smallest effect size of interest) to address all existing concerns. This will likely require additional data. We would expect that the sensitivity analysis enclosed in the revision is generally reproducible, and that an expert in the field, i.e. a researcher familiar with EEG, would be able to immediately judge its soundness from the presentation in the paper.

I appreciate that the extensive delay and now repeated requests for extended revisions with an uncertain outcome, which we generally avoid, are frustrating. If you decide against submitting a revision given the history and future requirements, please let us know in due course.

If you do opt for a revision, we would appreciate it if you could keep us informed about an estimated timescale for resubmission, to facilitate our planning.

#### Appeal request for COMMSPSYCHOL-23-0270A-Z

**** <> Tue, Nov 7, 2023 at 10:12 AM
Reply-To:
To:
Cc:

Dear Rick,

Thank you for your correspondence asking us to reconsider our decision on your Article, "A novel third process of episodic memory: Context familiarity is distinct from item familiarity and recollection". Having read the revised manuscript and point-by-point response to the reviewers which you submitted in appeal, I decided that we would be willing to send a suitably revised version of your manuscript back to the reviewers.

In detail, we require you to further revise the manuscript - and encourage you to subsequently revise the point-by-point reply - to fully align with the journal standards. I will highlight a few issues that are a precondition for re-review in the following, but we ask you to make sure that your revision is fully compliant with all of our format requirements, which are summarized on the following checklist: Communications Psychology formatting checklist and also in our style and formatting guide Communications Psychology formatting guide.

First, the journal places no strict upper limit on the length of an Article; the guiding principle is that reporting should be transparent and comprehensive and that key results, i.e. results that pertain to the core research question and are mentioned in the Discussion, should be presented in the main manuscript, not the supplement. There is no word limit for the Methods, and while we recommend the remainder of the Article (Introduction + Results + Discussion) does not exceed 5500 words, we will allow significantly longer manuscripts where this is necessary to report the results in great clarity.

We ask that the Methods section in the main manuscript reports all key analyses, those that were included in the previous version as well as those that were undertaken in revision (including those on additional datasets). Only ancillary tests and "sanity checks" should be relegated to the Supplementary Information. The results of the additional analyses you undertook should also be incorporated into the main manuscript. The same applies to the sensitivity analyses, which should be conducted for the key analyses you present.

Thus, as documented above, we only performed the ill-fated and ill-conceived statistical procedures of post-hoc power calculations because it was demanded of us by the Editor, against our will. We acknowledge here that the Reviewer 4 is entirely correct, and that the same core, foundational points against such post-hoc power and sensitivity analyses also prevail throughout the literature, which we excerpt as converging evidence below:

RESEARCH ARTICLE

### Post hoc power is not informative

Lacey W. Heinsberg, Daniel E. Weeks

First published: 01 June 2022 | https://doi.org/10.1002/gepi.22464 | Citations: 6

Read the full text >

PDF TOOLS SHARE

#### Abstract

Post hoc power estimates are often requested by reviewers and/or performed by researchers after a study has been conducted. The purpose of this commentary is to provide a heuristic explanation of why post hoc power should not be used. To illustrate our point, we provide a detailed simulation study of two essentially identical research experiments hypothetically conducted in parallel at two separate universities. The simulation demonstrates that post hoc power calculations are misleading and simply not informative for data interpretation. As such, we encourage authors and peer-reviewers to avoid using or requesting post hoc power calculations.

**The abuse of power: The pervasive fallacy of power calculations for data ana...**
John M Hoenig; Dennis M Heisey
*The American Statistician;* Feb 2001; 55, 1; ABI/INFORM Global
pg. 19

### Statistical Practice

#### The Abuse of Power: The Pervasive Fallacy of Power Calculations for Data Analysis

John M. HOENIG and Dennis M. HEISEY

It is well known that statistical power calculations can be valuable in planning an experiment. There is also a large literature advocating that power calculations be made whenever one performs a statistical test of a hypothesis and one obtains a statistically nonsignificant result. Advocates of such post-experiment power calculations claim the calculations should be used to aid in the interpretation of the experimental results. This approach, which appears in various forms, is fundamentally flawed. We document that the problem is extensive and present arguments to demonstrate the flaw in the logic.

KEY WORDS: Bioequivalence testing; Burden of proof; Observed power; Retrospective power analysis; Statistical power; Type II error.

culations as a matter of policy (Anon. 1995; Anon. 1998). We emphasize that these calculations are sought primarily with the thought that they are useful for explaining the observed data, rather than for the purpose of planning some future experiment. We even found statistical textbooks that illustrate the flawed approach (e.g., Rosner 1990; Winer, Brown, and Michels 1991; Zar 1996). Researchers need to be made aware of the shortcomings of power calculations as data analytic tools and taught more appropriate methodology.

It is important to understand the motivation of applied scientists for using power analysis to interpret hypothesis tests with nonsignificant results. The traditional, widely accepted standard has been to protect the investigator from falsely concluding that some treatment has an effect when indeed it has none. However, there is increasing recognition that a "reversal of the usual scientific burden of proof" (e.g.,

**INTRODUCTION**

Power analysis plays a key role in designing and planning prospective studies. For clinical trials in biomedical and psychosocial research, power analysis provides critical information about sample sizes needed to detect statistically significant and clinically meaningful differences between different treatment groups. Power analysis also provides critical information for evaluating cost–benefit ratios so that studies can be conducted with minimal resources without compromising on scientific integrity and rigour.

What is interesting is that some journals also ask for power analysis for the study data that were already analysed and reported in a manuscript before considering its publication. Although the exact purposes of such requests are not clearly stated, it seems that this often happens when manuscripts include some non-significant results. As such post hoc power analysis is conceptually flawed, concerns have been raised over the years.[1–4] Despite these warnings, some journals continue to ask for such information and use it as part of their decision process for manuscript publications.

As most research studies are conducted based on a random sample from a study population of interest, results from power analysis become meaningless, as the random component in the study disappears once data are collected. Power analysis shows the

Check for updates

© Author(s) (or their employer(s)) 2019. Re-use permitted under CC BY-NC. No commercial re-use. See rights and permissions. Published by BMJ.

[1]Department of Family Medicine and Public Health, University of California System, Oakland, California, USA
[2]Naval Health Research Center, San Diego, California, USA
[3]Leidos, San Diego, California, USA

**Correspondence to**
Dr Xin M Tu, Family Medicine and Public Health, University of California San Diego, La Jolla, CA 92093, USA;



**DISCUSSION**

Power analysis is an indispensable component of planning clinical research studies. However, when used to indicate power for outcomes already observed, it is not only conceptually flawed but also analytically misleading. Our simulation results show that such power analyses do not indicate true power for detecting statistical significance, since post hoc power estimates are generally variable in the range of practical interest and can be very different from the true power.

In this report, we focus on the relatively simple statistical model for comparing two population means of continuous outcomes. The same considerations and conclusions also apply to non-continuous outcomes and more complex models such as regression. In general, post hoc power analyses do not provide sensible results.

**Contributors** All authors participated in the discussion of the statistical issues and worked together to develop this report. AR, RR, SR and XT discussed the problems with the justification of post hoc power analysis and interpretation of such power analysis results within the contexts of their studies and how to approach and clarify the issues in clinical research. YZ, RH and XT worked together to develop

The manuscript has now been revised to have those post-hoc analyses removed, which did not affect any other results or conclusions. As such, it greatly strengthened the manuscript, and we very much appreciate the Reviewer's correct detailing of the issue.

Finally, in regard to those "misconceived" post-hoc analyses of power and sensitivity, extensively detailed above, it is important for us to also note that the notion for them actually originally stemmed from a "minor point" that was first requested by Reviewer 3 in the first round of Reviews in ~Sept. 2023, as copied below for full transparency:

> Minor points:
>
> Where participants compensated for their participation?
>
> What was the range of the years participants had been speaking English?
>
> With respect to power of the study: I understand that there was no a priori power analysis as the study was designed for another purpose. However, it would be nice to read some words about the sensitivity of the sample to detect the effect sizes relevant in the study.

This invalidation of Reviewer 3's conceptions, as provided by Reviewer 4's statistical expertise identifying the "lack of understanding" in the recommendation made by Reviewer 3, adds substantively to the array of demonstrations we have already provided thus far through three rounds of extensive responses to Reviewer 3's flawed reviews. This continues a common occurrence, whereby we have consistently shown that Reviewer 3's comments were based upon misconceptions and poor understandings- now independently confirmed by Reviewer 4. It should weigh against whatever residual misunderstandings Reviewer 3 may still have, as noted below.

**Reviewer #3 (Remarks to the Author):**

**I recognize the Editor's decision to give me the opportunity once again to evaluate whether my concerns were dispelled. However, given that the response letter is extremely long, a detailed evaluation is beyond what I can offer. I will only point out the major remaining issues.**

> First, we thank the Reviewer for the efforts of service to the industry provided by the effort of peer review. Second, we note for posterity here, that the Reviewer notes that the Author's prior Response letter is "very long"- but then neglects to acknowledge that it is only so long because it was responding to the extensively long (and wrong) Review that we were provided by this Reviewer, and to which we needed to provide clear refutation of in a comprehensive and point-by-point.

**In their response, the authors keep reiterating that my considerations are speculative. It is true there is no external validation of a delayed recollection / search process account. However, in my view, the same holds true for the context familiarity account. So, this summarizes very well what I think is the problem of the paradigm: it leaves too much space for speculations. To be clear, I don't think that the recollection or search process accounts are more likely, I just think they are as plausible as the context familiarity account**.

> *First, the Reviewer directly admits that her alternative account of the findings is entirely speculative- that should be an indictment of her own review, scientifically speaking, and effectively negate it*. Whenever possible, we should not deal with speculations, but with data. We noted in prior rounds of review that it is we who have provided clear, replicable, independent data to support our conclusions- which the other two primary reviewers found reasonable and persuasive. Yet, incredibly, the Reviewer 3 here states that "the same holds true" for our conclusions as being based upon speculation! They are not based upon any speculation but upon empirical observations of experimental data that persist after being subjected to rigorous controls and replication. Her comment here thus should hold no weight: it is just admitted speculations at this point, versus our actual data that we provide open access to publicly. She is free to disagree, and write competing commentaries, etc., but she should run her own study to refute it with her own data if wants to critique data with more than mere speculation.

**I'm still not convinced that there is any evidence for context familiarity in the reaction times. The authors did not reject the explanation that RTs for immediate recollection are faster than those for delayed recollection or a search process (context familiarity condition). The authors instead raised the point that the context familiarity source judgments were faster than those for recollection. In my eyes, it's possible that recollection was delayed during the item judgment (where RTs in the context familiarity condition were longer) and at the same time**

**for the subsequent source judgment, this information was readily available and led to fast responses**.

First, with all due respect, yes, we actually did providing compelling reasons to reject that alternative hypothesis about the response time data, and it was extensive discussed in several Discussion sections, including: *Section 5.3 "Behavioral Findings*", *Section 5.3.1 "Why might context familiarity effects occur later than item familiarity*?", *Section 5.4.2 "A delayed recollection process occurring after the ERP measures*?".

Second, the Reviewer's comment also presupposes that 'delayed recollection' even existed in this study, which the Discussion sections noted above thoroughly debunked as highly improbable, implausible, and lacking evidence.

Third, the Reviewer's proposal at the end of this comment about her account of the different speeds of recollection and context familiarity responses appears to (again) misunderstand and conflate the two different responses across the two different judgments of item memory and source memory. That is, they are two different response conditions (defined in Method section as comprising fundamentally different and mutually exclusive trials) that are measured in two different tasks (item recognition memory, and source memory judgments). From that, there were significant differences found to be statistically reliable in those contrasts and the Reviewer's account incorrectly presupposes that they are not significantly different while also not providing any data-based reasons for why they do significantly differ. Our account does just that and is thus a scientifically stronger account that remains unimpeached by the Reviewer's comment/misunderstanding above.

**With respect to the delayed recollection account, I think there is a misunderstanding. I did not mean that recollection would be delayed to several seconds or minutes later as in the Rivas-Fernández et al. (2020) study cited by the authors but just delayed within the item judgment display (from the 600 to 900 time window to the 800-1200 time window). In this perspective, I also have to reject the authors' argument that the delayed recollection account "would predict no differences in ERPs between the condition we call context familiarity and that of correct rejections because if there wasn't memory/processing yet (i.e. it occurred via a 'delayed recollection') then there would be no reason for the condition to differ from a non-memory condition baseline of correct rejections." In contrast, if recollection is delayed just by 200 ms, we would indeed predict differences between CRs and low confidence item memory + correct source memory**.

First, the reviewer begins this comment by admittedly back peddling from what she originally had proposed/said in the earlier round of reviews, because our subsequent response to it had clearly vanquished that misunderstanding as it was levied then. Now, the Reviewer is moving the goal posts further away, be re-defining the standards of what she was proved wrong about earlier. That is not a part of a good scientific process.

Second, the Reviewer's comment claims to reject the author's argument, but she fails to do so with any evidence or data- only via a merely speculative opinion which we can also then demonstrate is incorrect (see below). That is, she goes on to presuppose that, for the contrast of context-familiarity vs correct rejections, that she would predict to see significant ERP differences "if recollection is delayed by just 200 ms". <u>Let's unpack that just a bit here:</u>

1. The putative ERP correlate of recollection occurs from approximately 600 to 900 ms as a large positive-going effect, as widely found and agreed upon in the field.

2. We verify and validate that finding in the present experiment, where we establish statistically significant results of the so-called late/left parietal component "LPC" or parietal old-new effect, occurring precisely during that time of 600 to 900 ms, and at left parietal regions, and for the condition most agree with is indisputably 'recollection' (high confidence item recognition accompanied by correct source memory).

3. The Reviewer proposes, or hypothesizes, that the effect we attribute as "context familiarity" might actually be "recollection", if the recollection is "delayed by just 200 ms"

    a. That hypothesizes that the established time of recollection effects in the present study should then be shifted 200 ms forward from the 600 to 900 ms time window, and instead be observable from 800 to 1100 ms time window.

    b. This is precisely, and nearly identically, the same time window that we looked for exactly those effects in the present study (800 to 1200 ms), and the results directly and fully rejected the Reviewer's hypothesis here, because it did not find any ERP effects even closely reminiscent of recollection (positive-going effect at left parietal sites) but instead found consistent *negative-going* effects that were maximal at mid-central sites instead.

        i. It was literally a giant negative-going effect when she predicted a giant positive-going effect, and it occurred at the place were we hypothesized it to be- not the place that she hypothesized it to be.

c. We are all wrong at times, but it is important to admit it when it happens. This time it is the Reviewer who is wrong (again) but will not admit it. We can, and did, prove it.

d. All of this is already self-evident in the manuscript as previously written, and so required no further modifications in the manuscript to draw this conclusion.

**The authors argue that participants in the item familiarity condition would necessarily engage in a search process (and therefore we would see the LPN also in this condition) because they are asked about source information. However, it is also possible that they start this search process only after they have made their item judgment which they could easily make based on their available item familiarity signal**.

First, it is again a problematic comment that is weakened by merely being based upon “however its possible…”, which is speculation instead of evidence, whereas the conclusions drawn in the manuscript are based upon evidence and thus should bear more weight than the speculative possibility offered here by the Reviewer.

Second, the Reviewer ends the comment by stating that an item judgment could “easily be made based upon their available item familiarity signal”- but this is entirely unsupported (and refuted) by the data we provided in the manuscript. That is, if this hypothesis of the Review was true, then it would predict that there ought to be neural correlates of item familiarity evident while making the item memory judgment (if it was “easy” for the participant “based upon their available item familiarity signal”), but the data simply do not show that for the condition of context familiarity: it instead shows that there was reliable evidence of absence for the putative neural correlates of item familiarity (the FN400, 400 to 600 ms), and this was fully replicated in a large independent data set of other ERPs.

Furthermore, that hypothesis from the Reviewer fails a second time, because it predicts that the condition of context familiarity would actually be the same as item familiarity during item judgments, yet the opposite is found when we look for that in the data: the two conditions are found to reliably differ precisely during the time of the putative neural correlates of item familiarity (the FN400, 400-600 ms). Those reliable physiological differences between the conditions of context familiarity and item familiarity were found to survive rigorous control analyses designed to test & falsify them and were also found to be independently replicable. In contrast, the Reviewer’s hypothesis as stated in the comment above offers to answer nor account for these findings.

We thus prove here that the Reviewer’s comment is wrong- not in opinion, but in fact, and based upon multiple lines of converging data. All of this data was in the prior manuscript’s results and figures, and so remain unchanged in the present version.

**The authors argue that they do not want to change their way of measuring source memory because they want to do it in the same way as they did before. I appreciate this approach. However, I do not think this should prevent the authors from additionally reporting source memory in the way I suggested, especially as they do not argue that this approach is invalid. (By the way, I meant that %source correct/incorrect should be calculated based on all trials). Moreover, I checked two of the references for the calculation of the source memory measure and they do not exclude "unknown" source judgments. Foley et al. explicitly say that on, "the total number of words attributed to the correct source ... was divided by the total number of items correctly identified as old". Finally, I’d like to note that although this experiment was not about individual differences. I always find it helpful to consider between subject comparisons when thinking about the meaningfulness of a specific measure**.

In prior rounds of Reviews, the same Reviewer literally acknowledged that the new way of calculating source memory that she invented for reviewing the present manuscript was in fact flawed, and would be improper to do, after we pointed out the fallacies of it in the Response to Reviewers. Now the same Reviewer may have forgotten that and wants it added back in. We will not do that, for the reasons stated in the Method section detailing that (2.3.2 Measuring source memory performance.). As a short primer, what the review is suggesting to have done here is a flawed approach and invents a new calculation method for measuring source memory, which we could not find cited anywhere in the field, and which through three rounds of reviews she has not been able to cite support for in the literature, either. We dispense with it here and point to it as another reason accumulating to exercise appropriate Editorial discretion in evaluating the decision to publish the manuscript.

Regarding the minor point about references that the Reviewer cited: this was another error on the part of the Reviewer, because those references were included as part of quotes from statements made by other authors that we had quoted & cited in that particular section. So, they were not our references, but theirs. But to ease any concern, we easily removed those two references from the updated manuscript’s section, and the premises, and support for them, remain unaffected, because the other references provided continue to fully substantiate our approach.

**Previously, I argued that relative timing together with morphology is an important characteristic when identifying ERP components. I highlighted that the LPN usually occurs directly after the positive component associated with recollection and that this also applies to the BCN. The authors responded that this "is precisely what was not found in the present studies. Instead, those studies never found the negative ERP effects after a positive component associated with recollection- and these null findings were quantified with Bayes Factor analyses, demonstrating meaningful evidence of absence". I do not agree. It is true that the authors did not find a difference between their context familiarity condition and correct rejections for the LPC. However, the component itself was present**.

First, the Reviewer then acknowledges that "it is true" that we did not find a significant difference between ERPs for context familiarity and correct rejections (even though she somehow preceded that by stating that she "did not agree"). But then the Reviewer goes on to state that "the component itself was present":

1. It is not clear what that even means when the Reviewer states that "the component itself was present". That facet, and 'component' is not relevant to the present studies of ERP effects, because ERP effects are defined by subtractions of two different conditions, not waveforms for single components.

2. The Reviewer is apparently misunderstanding a few things. Namely, that the Reviewer appears to be thinking that there was a BCN effect that followed an LPC effect in timing: that did not happen in the present experiments for context familiarity.

3. The reviewer seems to be conflating that with studies in the literature of a different ERP effect, the LPN (which review extensively in Discussion), in which the LPN effect followed an LPC effect- but for the same condition/contrast. That is quite different from our findings, which we found both an LPC effect and a BCN effect- but for fundamentally different conditions (recollection, and context familiarity, respectively), and at fundamentally different times (600-900 ms, 800 to 1200 ms, respectively), but they never co-existed together in sequence for the same singular condition when the conditions were directly specified.

4. This seems to be another instance yet again of a core misunderstanding by this Reviewer, which we are glad to have an opportunity to respond to and refute for clarification purposes, much as Reviewer 4 did about this Reviewer's original misconceived suggestion to perform the post-hoc power analyses already addressed in the sections above. This places additional context within which to weigh the validity of the Reviewer's review amid otherwise positive reviews.
