## Supplementary material for "A third kind of episodic memory: Context familiarity is distinct from item familiarity and recollection": Second Response to Reviewers at NatureCommsPsych

FLORIDA TECH
College of Psychology and Liberal Arts

March 23, 2023

RE: Re-Consideration of Manuscript (COMMSPSYCHOL-23-0270B).

Dear Dr. Schiffer,

Hello. Thank you for the kind re-consideration of our manuscript, *A third process of episodic memory: Context familiarity is distinct from item familiarity and recollection* (ID# COMMSPSYCHOL-23-0270B). Enclosed please find a copy of our revised manuscript which has been updated in accordance with incorporating the quality feedback provided by the Reviewers, as well as our response to the Editor below.

**Editor Comments: As you will see below, Reviewer #3 voiced not only methodological and conceptual concerns that remain unresolved, but also noted the small sample size. You will also see that we obtained additional comments from Reviewer #1 on the most critical issues raised by Reviewer #3, a process we occasionally elicit when the reports are in conflict and we believe strongly that additional input will benefit the decision.**

We are grateful that both of the two Reviewers found the revised manuscript to have satisfactorily addressed their previous comments and to have substantially strengthened the manuscript to be suitable for publication. Reviewer 3 also commended good progress and responded that she had satisfied at least half of her prior concerns. While it is rarely if not impossible to satisfy all questions of all scientists, in our present re-submission we have aimed to provide thorough response details to Reviewer 3's remaining comments, in hopes that further information helps clarify existing understandings.

Reviewer 3 makes several good points but also several that we believe may be better suited towards separate commentary papers about opinions, or future empirical studies that we hope will stem from reporting the present findings, and we have added thorough discussion of several of these issues in the updated manuscript's Discussion section, including a dedicated section fruitful steps for future research (Section 5.5.4) . Some of the Reviewer comments are, in effect, also creating new standards in the field where they have not existed before (i.e., the criteria for identifying cognitive processes using ERP correlates). Similarly, some comments appear to be 'moving the goalposts' of a 'moving target' after we had already responded to those points in prior rounds in ways that the Reviewer said that she agreed with our reasoning (i.e.: novel ways to measure source memory that were first suggested in the prior round of reviews) yet now changes to request other things measures instead. We have included these agreed-upon points in the

Florida Institute of Technology
150 W. University Blvd.
Melbourne, FL 32901-6975

321-674-8104

floridatech.edu/copla

FLORIDA TECH
College of Psychology and Liberal Arts

Discussion as directions for valuable future work (Section 5.5.4). Other points raised are very good ones that we took careful time to thoroughly respond to, and we are grateful for the Reviewer's kind attention to detail. We believe that her dedication to quality has uplifted our manuscript, and we are grateful for her collegial challenges that make us all better.

**Editorially, we have significant concerns about the inclusion of analyses at less than 80% power for the target effect size. Some of your control analyses, including those that speak to some of the referees' concerns fall within that range. The most recent delay is a result of another (second) statistician agreeing to vet the manuscript and subsequently withdrawing from the process; although they did not submit a report, they highlighted that they were unable to reproduce the sensitivity analysis and therefore unable to comment. In sum, we would welcome a revised manuscript that addresses all of Reviewer #3's concerns and especially their point about how process pure the conditions are. However, in keeping with the journal's focus on reproducibility and strength of evidence, the revision would need to present analyses that are sufficiently sensitive (>=80% power for the smallest effect size of interest) to address all existing concerns. This will likely require additional data. We would expect that the sensitivity analysis enclosed in the revision is generally reproducible, and that an expert in the field, i.e. familiar with EEG, would be able to immediately judge its soundness from presentation in the paper.**

We very much appreciate the Editor's recommendations on the power analysis and have now updated the revised manuscript to report these results with far better clarity Section 2.6.3) . We noticed that in the previous version of the manuscript we had omitted a few of the input data (e.g. effect size, sample size) as typographic oversights, but this has now been remedied. Of note, each result of the power analysis has been updated to have the data needed to be reproduced, and we also included a copy of the raw outputs from G-Power programmatic analyses as an addendum to this letter for full transparency and reproducibility by anyone interested. We also re-organized the Method section describing the power analyses (Section 2.6.3), focusing first on the novel behavioral findings which were not included in the original 2012 paper, and then moving to the ERP analyses which replicated and expanded upon what was previously reported in 2012. Of note here is that the main behavioral findings had power values above .80 (.823 and .999) as was requested by the Editor. Furthermore, one of the ERP analyses had a power value of .952, and another one was also nearly at .80 (.76). We hope that the Editor finds this to be both helpful and compelling, as well as clearing any remaining points of confusion that might have existed (though see further

Florida Institute of Technology
150 W. University Blvd.
Melbourne, FL 32901-6975
321-674-8104
floridatech.edu/copla

College of Psychology
and Liberal Arts

elaboration in the sections below). An excerpt of the revised power analysis in the updated manuscript is provided below:

> *First, the main behavioral effect of response time (RT) differences for item recognition judgments among context-familiarity and item-familiarity conditions (N = 38, Cohen's d = .481) had a reasonably high power of .823, and the effect of RT differences observed between context-familiarity and recollection response had a power of .999 (N = 38, Cohen's d = 1.21). That sensitivity to detecting differences between RT's for item- and context-familiarity judgements was naturally reduced in the ensuing control analysis requested by a Reviewer that excluded the high confidence recognition responses from the item familiarity condition, which then provided a power of .530 with the inherently lower signal to noise ratio (SNR) (N = 37, Cohen's d = .344). The ERP correlate of the same control analysis (removing the high-confidence item familiarity responses to match memory strength, see Results) had a power of .952 (N = 3, Cohen's d = 3.21, one-tailed), and the larger sample size available of the same analysis in Experiment 2 had a resulting statistical power of .580 (N = 12, Cohen's d = .569, one-tailed). Moving to the main ERP analyses, for the investigation's finding of ERPs for context familiarity vs correct rejections at Cz from 800-1200 ms, the resulting power was .681 (N = 27, Cohen's d = .486). Next, for the main finding dissociating item- and context-familiarity ERPs within-subjects there was a power value of .626 (N = 11, Cohen's d = .762). Finally, the between-subject effect for context- vs. item-familiarity (N = 9, N = 17, effect size of 1.14) had a power of .760*.

Conceptually, it is important to note that the power analysis we performed was a post-hoc analysis as opposed to the typical a priori one commonly seen in the literature, specifically because that was what was requested by a Reviewer in the first round of Reviews. That Reviewer specifically noted that it was understood that since this was a secondary analysis of an existing dataset that had already been collected (and published in 2023), it thus could not have an a priori power analysis performed to determine sample size and therefore requested a post-hoc one to better inform the relative amount of power that existed to detect differences in the present findings. It was an ancillary analysis to help better inform the significant results we observed. As noted above, in several of the primary novel analyses, our power exceeded levels of .80; however, in some of the other ERP analyses the power ranged from .62 to .68.

As we note in the manuscript, these slightly lower values simply meant that the study was slightly under-powered to detect those findings. However, the significant findings were detected even at these lower power values. Furthermore, these findings were then independently replicated. This likely means that the effects

Florida Institute of Technology
150 W. University Blvd.
Melbourne, FL 32901-6975
321-674-8104
floridatech.edu/copla

FLORIDA TECH
College of Psychology
and Liberal Arts

seen in our study are real and meaningful effects that are even more likely to be observed in future studies using larger sample sizes. The below-threshold (i.e. .80) power observed in some results in our study would have been more problematic if it was for a null finding, not the rejection of the null that we observed. That is a strength of our study, not a weakness. The only way that such kinds of reduced power could be seen as a weakness would be if it was a non-significant effect (though we instead found them to be a significant effect), or, if it was for a finding that may have in fact been a type II error (a false positive). The fact that we were able to successfully reproduce each of those findings with replications in other independent data sets greatly reduces the likelihood of a type II error (i.e., false positive), as we have now also noted in the updated Section 2.6.3 of the manuscript. Therefore, we hope that our initial misunderstanding about the power can be successfully resolved moving forward.

Overall, our approach and interpretations are grounded in the core premises of the field of cognitive neuroscience for the identification of cognitive processes. It's an example of rigor in replication science, whereas other studies typically fall short of the range of rigorous testing and protocols that have plagued psychological science in the past. We believe our findings are valid, and are demonstrably reproducible; furthermore, we believe that no viable alternative account has been provided to explain the range of the findings converging across behavior, physiology, and multiple different laboratories. Where dissent exists, we believe that it is healthy, and that the best route is for the findings to inspire follow-up studies capable of refuting our interpretations based upon providing multiple experiments of new data designed to test these findings while demonstrating how they vary under conditions alternatively proposed to differentially account for them.

We would be glad to make any other changes that may be requested. Accordingly, we would also be more than happy to continue any points of discussion via direct call or video conference if it were helpful. Moreover, we have included source data for each figure, and the Data Availability statement links to full EEG datasets from the primary data set published previously in *Psychophysiology* (2023). Thank you for the excellent editorial handling of the manuscript, effort, & consideration.

Very Sincerely,

Richard J. Addante

Richard J. Addante, Ph.D.
Associate Professor of Psychology
NIH LRP Fellow, National Institute of Neurological Disorders & Stroke

Florida Institute of Technology
150 W. University Blvd.
Melbourne, FL 32901-6975

321-674-8104

floridatech.edu/copla

College of Psychology
and Liberal Arts

### Addendum: Power Analysis producibility data

***Note: all effects assessed were statistically significant (see manuscript)***

[1] -- Monday, March 18, 2024 -- 14:46:41
Response Time for item recognition judgments: context- vs. item-familiarity (Fig.2)
t tests - Means: Difference between two dependent means (matched pairs)
Analysis: Post hoc: Compute achieved power
Input: Tail(s) = Two
Effect size dz = .481
α err prob = 0.05
Total sample size = 38
Output: Noncentrality parameter δ = 2.9650831
Critical t = 2.0261925, Df = 37
Power (1-β err prob) = 0.8231311

[2] -- Monday, March 18, 2024 -- 14:46:56
RT for item recognition judgments: context-familiarity vs. recollection (Fig. 2)
t tests - Means: Difference between two dependent means (matched pairs)
Analysis: Post hoc: Compute achieved power
Input: Tail(s) = Two
Effect size dz = 1.21
α err prob = 0.05
Total sample size = 38
Output: Noncentrality parameter δ = 7.4589409
Critical t = 2.0261925, Df = 37
Power (1-β err prob) = 0.9999999

[3] -- Monday, March 18, 2024 -- 14:50:32
Response Time for item recognition judgments: context- vs. item-familiarity$_{low}$
t tests - Means: Difference between two dependent means (matched pairs)
Analysis: Post hoc: Compute achieved power
Input: Tail(s) = Two
Effect size dz = .344
α err prob = 0.05
Total sample size = 37
Output: Noncentrality parameter δ = 2.0924703
Critical t = 2.0280940
Df = 36, Power (1-β err prob) = 0.5305305

[4] -- Monday, March 18, 2024 -- 14:56:28
ERP effect of context-familiarity vs. item-familiarity$_{low}$, Exp. 1 (Fig 5C)
t tests - Means: Difference between two dependent means (matched pairs)
Analysis: Post hoc: Compute achieved power
Input: Tail(s) = One
Effect size dz = 3.21
α err prob = 0.05
Total sample size = 3
Output: Noncentrality parameter δ = 5.5598831
Critical t = 2.9199856
Df = 2
Power (1-β err prob) = 0.9522621

Florida Institute of Technology
150 W. University Blvd.
Melbourne, FL 32901-6975
321-674-8104
floridatech.edu/copla

FLORIDA TECH
College of Psychology and Liberal Arts

[5] -- Monday, March 18, 2024 -- 14:54:55
ERP effect of context-familiarity vs. item-familiarity$_{low}$, Exp. 2 (Fig. 7B)
t tests - Means: Difference between two dependent means (matched pairs)
Analysis: Post hoc: Compute achieved power
Input: Tail(s) = One
Effect size dz = .569
α err prob = 0.05
Total sample size = 12
Output: Noncentrality parameter δ = 1.9710738
Critical t = 1.7958848
Df = 11
Power (1-β err prob) = 0.5804899

[6] -- Monday, March 18, 2024 -- 15:02:51
ERP effect of context-familiarity vs correct rejections Cz, 800-1200 ms (Fig. 4)
t tests - Means: Difference between two dependent means (matched pairs)
Analysis: Post hoc: Compute achieved power
Input: Tail(s) = Two
Effect size dz = .486
α err prob = 0.05
Total sample size = 27
Output: Noncentrality parameter δ = 2.5253301
Critical t = 2.0555294
Df = 26
Power (1-β err prob) = 0.6812790

[7] -- Monday, March 18, 2024 -- 15:03:15
ERPs: context familiarity vs item-familiarity, Cz,300-500 ms, within-subjects (Fig. 5)
t tests - Means: Difference between two dependent means (matched pairs)
Analysis: Post hoc: Compute achieved power
Input: Tail(s) = Two
Effect size dz = .762
α err prob = 0.05
Total sample size = 11
Output: Noncentrality parameter δ = 2.5272681
Critical t = 2.2281389, Df = 10
Power (1-β err prob) = 0.6258220

[8] -- Monday, March 18, 2024 -- 15:04:05
ERPs: context familiarity vs correct rejections, Cz,8-1200, between-subjects (Fig. 6A)
t tests - Means: Difference between two independent means (two groups)
Analysis: Post hoc: Compute achieved power
Input: Tail(s) = Two
Effect size d = 1.14
α err prob = 0.05
Sample size group 1 = 9, Sample size group 2 = 17
Output: Noncentrality parameter δ = 2.7654378
Critical t = 2.0638986, Df = 24
Power (1-β err prob) = 0.7559807

Florida Institute of Technology
150 W. University Blvd.
Melbourne, FL 32901-6975
321-674-8104
floridatech.edu/copla

Dear Dr. Schiffer,

Hello. We sincerely apologize for a simple and earnest misunderstanding on our part. Specifically, we initially reported statistical power as sensitive to detecting differences, rather than reporting sensitivity to detect minimum effect sizes at .80 power. Following this, we made the minor adjustments needed for our calculations for the respective analyses, which have now been included in an updated revision of the manuscript. These adjustments further support our findings. While we regret the initial misunderstanding, we appreciate the Editorial recommendation, as it prompted us to enhance the rigor of our statistical approaches, based on clearer a priori hypotheses. This ensured that the data were sufficiently sensitive. Therefore, we respectfully request an Appeal.

The description of the sensitivity analyses is now included in Method section 2.6.3 and 2.6.2, as well as reported in-text for each direct comparison, and the manuscript is also now supplemented with a full Summary Table that we have added to the Supplementary Materials (and included here below). Specifically, and importantly, as is demonstrated in Supplemental Table 1, **every significant principle analysis performed across Experiments 1 and 2 were identified as being either sufficiently sensitive to detecting minimum necessary effect sizes at .80 power and/or independently replicated in mutually exclusive datasets collected nearly ten years apart in different laboratories (i.e. not a Type I error)**.

**Similarly, any relevant null findings were associated with a minimum Bayes Factor of at least 3.0 conventionally taken to represent a meaningful evidence of absence and/or replicable in the independent dataset sets with sufficiently sensitive Bays Factor evidence of at least 3.0 (and thus not due to being under-powered for a Type II error**). *That is, the results indicate that the study was both sufficiently powered and sensitive to detecting the observed results (*it cannot be attributed as Type I or II errors, but instead are demonstrably valid). Collectively, this effort provided convergent, consistent confidence in the validity of conclusions as reliable results. In short, it really strengthened the paper.

The previous Editorial letter raised two critical questions: **whether the findings were sensitive enough to detect reliable effect sizes at .80 power and if they were reproducible. Both questions can now be unequivocally answered 'yes'. We have demonstrated that the findings are indeed sufficiently powered, sensitive, valid, and replicable.** The manuscript reports replications of the main findings across four previously published datasets and also confirms old findings in new data. Utilizing archival datasets already vetted through publication, we can be confident that the data is not a spurious finding but has been rigorously validated by editors and reviewers from reputable journals (*PNAS*, *Neuropsychologia*, *Neuroimage*, *Psychophysiology*, and the *European Journal of Neuroscience)*. The new findings presented here are significant and valid, and we agree with Reviewers that it can make important new contribution to the field for both specialists and generalists alike.

As such, we humbly Appeal to respectfully request re-consideration of the prior decision. We believe that the paper is now stronger than ever, and its previous version had already been endorsed by 2.5 out of the 3 Reviewers- for which we now also provide substantive replies to the lingering questions from Reviewer 3 that we hope will be deemed satisfactory. We sincerely believe that the paper has been handled excellently here at NCP and with exceptional Editorial discretion, and we hope that our minor error in understanding the calculation being requested does not stand in the way of what we and the Reviewers have both found to be exciting potential for publication.

### Supplemental Table 1.0

| Contrast | Figure | Exp. 1: If significant: effect size (d), minimum effect size with .80 power (dz), power (B) | Exp. 2: If significant: effect size (d), minimum d with .80 power (dz), power (B) | Exp. 1: Bayes Factor (if null) | Exp. 2: Bayes Factor (if null) | Sufficiently sensitive, powered, or replicated? |
|---|---|---|---|---|---|---|
| Behavioral RT: cFam v. iFam | 2 | d= .48, dz = .47, B = .82 | N/A | - | - | Yes |
| Behavioral RT: cFam v. Rec. | 2 | d= .76, dz = .47, B = 99 | N/A | - | - | Yes |
| Behavioral RT: Rec. v. iFam | 2 | d= 1.24, dz = .47, B = .99 | N/A | - | - | Yes |
| ERPs: old-new, FN400 | 3, S2 | d= .53, dz = .34, B = .99 | d= .82, dz = .34, B= .99 | - | - | Yes |
| ERPs: old-new, LPC | 3, S2 | d= .42, dz = .34, B = .91 | d= .41, dz = .34, B= .91 | - | - | Yes |
| ERPs: old-new, BCN | 3, S2 | d= .39, dz = .34, B = .88 | d= .58, dz = .34, B= .99 | - | - | Yes |
| ERPs: cFam v. CR, FN400 | 4, S1 | ns | ns | 5.12 | 1.25 | Yes |
| ERPs: cFam v. CR, LPC | 4, S1 | ns | ns | 4.32 | 3.17 | Yes |
| ERPs: cFam v. CR, BCN | 4, S1 | d= .49, dz = .49, B = .80 | d= .56, dz = .50, B= .96 | - | - | Yes |
| ERPs: iFam v. CR, FN400 | 4, S1 | d= .46, dz= .58, B= .63 | d= .61, dz = .47, B= .98 | - | - | Yes |
| ERPs: iFam vs CR, LPC | 4, S1 | ns | ns | 1.27 | 3.76 | Yes |
| ERPs: iFam v. CR, BCN | 4, S1 | ns | ns | 1.83 | 5.66 | Yes |
| ERPs: Rec v. CR, FN400 | 4, S1 | d= .54, dz = .49, B = .86 | d= .84, dz = .38, B= .99 | - | - | Yes |
| ERPs: Rec v. CR, LPC | 4, S1 | d= .57, dz = .49, B = .89 | d= .64, dz = .38, B= .99 | - | - | Yes |
| ERPs: Rec v. CR, BCN | 4, S1 | ns | ns | 4.91 | 5.72 | Yes |
| ERPs: cFam v. iFam | 5, 7 | d= .76, dz = .81, B = .76 | d= .48, dz = .58, B= .66 | - | | Yes |
| ERPs: cFamLow v. iFamLow | 5, 7 | d= 3.21, dz = 2.3, B = .95 | d= .60, dz = .77, B= .62 | - | - | Yes |
| ERP: cFam v. iFam, B/w subj | 6 | d= 1.2, dz = 1.1, B = .86 | d= .79, dz = 1.1, B= .59 | - | - | Yes |
| ERP: cFam v Miss, 800-1200 | S3 | d= .44, dz= .56, B= .60 | d= .45, dz = .49, B= .74 | - | - | Yes |
| ERP: cFam v Miss, 200-400 | S3 | d= .49, dz = .56, B = .69 | d= .50, dz = .49, B= .82 | - | - | Yes |

**Table S1. Summary of effect size, power, and Bayes Factor for principle findings.** Table provides a comparison of analyses in Experiment 1 and Experiment 2. If the effect was statistically significant beyond $p < .05$, then the table provides the effect size (Cohen's d), the minimum effect size (dz) necessary for a power level of .80 as determined by a sensitivity analysis, and the observed power (B). If the result was instead found to be non-significant at $p < .05$ (a relevant null finding), the Bayes Factor (BF) is provided to assess the evidence for absence of the effect. Convention follows BF of 3 or greater to represent a meaningful evidence of absence (see Method section 2.6.1). In order to inform whether significant results represent a Type I error (a false positive), or if null findings represent a Type II error (a false negative), or if the results were valid representations of true effects, the right column provides a summary indication of if each particular effect was found to be replicated among the other experiment and/or if it met the standard threshold for being sufficiently sensitive to detecting a minimum effect size for its sample size. Every contrast was sufficiently sensitive, powered, and/or replicated, and thus not attributable to Type I/II errors. N/A indicates not available, since the archival data provided from Exp. 2 (2012) did not collect response times; though those effects in Exp. 1 were identified as being sufficiently sensitive and adequately powered to detect reliable effects.

A note of reminder for this particular manuscript, amid the shuffle of the many other manuscripts being evaluated, is that in its prior status it had received the Editor's positive invitation to "revise & re-submit" form the Editor, after 2 of the 3 Reviewers had reported being entirely satisfied with our earlier revisions, and the 3rd remaining reviewer had noted having "half" of her prior comments fully satisfied, with only a few additional comments remaining (which our present responses had been submitted in reply to, hopefully further addressing towards complete amelioration).

The hold up from there had simply been an Editorial decision about a sensitivity analysis that had requested (and which we misunderstood to be asking for a calculation of statistical power, as opposed to the computationally related statistical sensitivity of effect sizes): that was the premise for the rejection that we are now appealing. Our appeal now provides the full characterization and report of the sensitivity analysis, in hopes that this minor change in calculation (which made the manuscript stronger). We hope that this can facilitate the continuation of the pre-existing process of revise & resubmit via sending the updated Revision back to the remaining Reviewer/Editor for the last stages of evaluation. While it is rare to find complete consensus among any three Reviewers, we believe the present 2.5 of 3 Reviewers is already a very strong position, and that this may even increase by further persuading the additional Reviewer with the present revisions, and ultimately permit the Editorial discretion needed in decision-making about the merits of publication.

Thank you for your time, effort, and continued consideration.

Sincerely,

Richard J. Addante

Richard J. Addante, PhD
Associate Professor of Psychology & Biomedical Engineering
Director, Neuroscience Research Laboratory
Florida Institute of Technology

FLORIDA TECH
FLORIDA'S STEM UNIVERSITY™

National Institutes of Health
LRP
AMBASSADOR
LEARN ABOUT THE AMBASSADOR NETWORK
www.lrp.nih.gov

REVIEWER REPORTS:

**Reviewer #1 (Remarks to the Author):** ***The revised manuscript has been substantially strengthened;  authors have addressed all of my concerns***.

**Reviewer #2 (Remarks to the Author):** ***The authors have addressed my comments and I commend them on a thorough revision***.

We sincerely thank both Reviewer 1 & 2 for their dedication and generous contributions to the review of the manuscript. Their insightful suggestions were indispensably helpful.

**Reviewer #3 (Remarks to the Author):**

**The authors put great effort in their revision of the manuscript and I am impressed by the additional analyses as well as by the passionate responses to all the points previously raised by the reviewers. Some of my comments have been answered to my satisfaction. However, I'm afraid but my main concerns remained and I still think the data provided does not justify the big conclusions made by the authors. I still think, the endeavor of finding an ERP correlate for context familiarity is worthwhile but that for such strong claims to make the authors would need to find the BCN in another paradigm**.

We thank the Reviewer for her time and expertise in providing a thorough and thoughtful review and are glad that many of the original comments have been answered to her satisfaction. Many points raised are very good ones that we took careful time to thoroughly respond to, and we are truly quite grateful for the Reviewer's kind attention to detail. We believe that the Reviewer's dedication to quality has certainly uplifted our manuscript, and we are exceptionally grateful for the collegial challenges that make us all better.

In regard to the comment here about thinking that the conclusions are not justified by the data, we respectfully, and alternatively, do not see how the conclusions are not justified by the data, and concur with the other two reviewers that it has been substantially strengthened in response to the thoughtful and helpful input from all three Reviewers. Our procedures provided an extensive rigorous process to systematically test the conclusions being drawn, trying repeatedly to disprove them, but the various control analysis continued to consistently support them instead, thus the submission of the claims for consideration of publication in the literature. Furthermore, when Reviewers previously suggested analyses that could disprove the conclusions, those too ended up supporting the conclusions instead. *We have, additionally, updated the manuscript with sensitivity analyses that emphasize that every principle contrast performed to analyze the data is sufficiently sensitive to detect the minimum effect size using a power of .80 and/or is independently replicated in Experiment 2's data that was identified as being sufficiently sensitive/powered. That is, we now show that the results are not due to Type I nor Type II errors (see Method sections 2.6.3 and 2.6.2, and Supplemental Table 1*).

We also agree with the Reviewer that finding the BCN in another paradigm will be a useful endeavor inspired from the present work, but we believe that it is one that is best suited for future studies to do, which we have now Discussed in a new section of the manuscript we have dedicated to Future Directions (Section 5.4.4). The present work includes five different data sets, all previously vetted by peer-review and published in highly-ranked journals  (i.e., *PNAS*, *Neuroimage*, *Psychophysiology* [1-5]); and towards the goal of reproducibility, is importantly strengthened by the fact of utilizing similar paradigms- not different ones.

Within these similar paradigms useful for assessing exact reproducibility, we underscore two points: first, that they are highly sophisticated paradigms that include combinations of both item recognition confidence and source memory confidence judgments- which otherwise would each be considered separate paradigms of their own. Second, we emphasize that the findings we report and ascribe to the mnemonic process of context familiarity were actually found convergent across several different paradigmatic measures (i.e. simple old-new recognition [Fig. 3], context familiarity as compared to correct rejections [Fig. 4], context familiarity vs item familiarity [Fig.5], separately identified between subjects [Fig. 6], and then independently replicated all of those separate measures in other data sets from a different laboratory [Figs. 7, S1, S2], and vs recognition misses as well [Fig. S3]; and also previously identified via source correct vs. incorrect trials in the original report in 2012). That amounts to the effects being identified in at least six different paradigmatic measures, and independently replicated, too.- while being grounded in an established theoretical framework (the BIC model) This is also a much, much higher bar and standard that has been applied to other pioneering ERP studies of episodic memory in the past, as is further detailed in sections below in response to R3.1.

As is normal in the scientific process, different paradigms can and should be utilized in follow up work inspired by these original findings. Additionally, while observations of convergent findings across different paradigms can be quite useful, they are not necessary: consider a plethora of situations across scientific disciplines whereby very sensitive effects are obscured, conflated, and difficult to disambiguate without having the precise scientific conditions at hand (e.g. particle physics, biochemistry, etc.), whereby a very delicate finding only emerges to become clear under precise experimental conditions. For instance, as was noted by Yonelinas (2009)[6] about earlier findings of familiarity contributing to source memory[7]: "*under just the right conditions, familiarity can support accurate source judgments*".  Observing a delicate, sensitive, and elusive component of human cognition under precise, rigorously controlled experimental conditions here makes the finding no less true and likely makes it even that much more valuable to communicate (i.e. a Black Swan effect remains a true effect, and an impactful one).

We wholeheartedly encourage exploring other paradigms in the future to test and potentially disprove the conclusions that are drawn from the present evidence, but do not think it adds more to what is already a full and substantiated report here. Skeptics are free (and encouraged) to disagree with our interpretations, but then it requires more than just speculation about what it ‘might’ alternatively be and instead requires engagement in testing it with several novel studies of their own to provide explanatory evidence for how our consistent reproducible findings were observed, and what factors future studies manipulated that revealed support for an alternative account. Indeed, we encourage and applaud such efforts. The motivation of future studies is another good reason why the present work should warrant publication in the literature.

In sum, and in due respect for Reviewer 3, we *do* think the extensive data reported in our study justifies the (big) conclusions we have made. If there are no objective points in the methods used or the data reported to justify withholding the publication of our results, nor obvious confounds available, then they should be made available to the field for assessment and placed under scrutiny by other scientists for the shared advancement of science.

Overall, our approach and interpretations are grounded in the core premises of the field of cognitive neuroscience for the identification of cognitive processes. It’s an example of rigor in replication science, whereas other studies typically fall short of the range of rigorous testing and protocols that have plagued psychological science in the past.  We believe our findings are valid, and are demonstrably reproducible; furthermore, we believe that no viable alternative account has been provided to explain the range of the findings converging across behavior, physiology, and multiple different laboratories. Where dissent exists, we believe that it is healthy, and that the best route is for the findings to inspire follow-up studies capable of refuting our interpretations based upon providing multiple experiments of new data designed to test these findings while demonstrating how they vary under conditions alternatively proposed to differentially account for them

**Response to R3.1:**

**It is clear that in many studies the conditions in which FN400 and LPC effects are elicited are not process pure. The original work, however, that identified the components took very much care that the two components were associated with properties that were expected only for one of them but not the other. For example, the FN400 was shown not to differentiate between old items and similar lures, to vary with levels of confidence and to be present although a response deadline is applied, all of which is not assumed for recollection but for familiarity. In contrast, the LPC is larger for remember trials than high confidence trials or trials with correct compared to incorrect source judgments, all of which is not assumed for familiarity but recollection. However, here, the operational definition of context familiarity is low confidence item memory with correct source memory which reads just as a mixture of item familiarity and recollection. To be clear, I like the idea of a context familiarity signal and find it a great post-hoc explanation for the 2012 data. However, to strengthen the link between the observed ERP effect and the construct of context familiarity, I find the operational definition too weak (see below for further elaboration of this point). I think that this has become evident across all three**

**reviews which all have raised the alternative interpretation of a controlled memory search process**.

First, a point of clarification and mutual agreement: we are not claiming process-pure ERP measurements of memory, but instead we consistently make note of what Rugg & Curran's Review paper on the ERP correlates of recognition[8] (as well as our own previously published papers described as the 'putative neural correlates' [2-5,9-13]) as that the ERP effects do not hold any particular 1:1 mapping to cognitive processes, per se, but instead serve as the putative neural correlates. We maintain that position here in the present findings (see Introduction, Section 5.3, Section 5.3.1, see also Section 4.2.6, Response to prior Reviews R3.1). Of note to the point made by the reviewer here and below about requiring even more findings from other paradigms: the original studies of the FN400 and LPC didn't require cross-paradigm validations in order to publish their findings to communicate to the broader scientific community- in fact to the contrary: once published as single studies or single paradigms, the scientific community responded by systematically assessing the findings across various additional paradigms as the progress of science continued. As such we believe it makes more of a case for publication of our findings, not less.

Second, we believe that the Reviewer's next part of the comment here fundamentally mischaracterizes the history of literature on the ERP correlates of episodic memory processes- especially and particularly when stating that "*The original work that identified the components…[showed that] the FN400… varied with levels of confidence and despite application of response deadlines*" and that "*The LPC is higher for remember trials than high confidence trials or in with correct vs. incorrect source judgments*". We do not find that those are accurate representations of the historical studies, since the "original" studies reporting these effects did not perform those comparisons or paradigms, but rather, the Reviewer is making a good summary of the broader literature that occurred afterward, over the scope of a few decades worth of different studies, as detailed below. We believe that our present findings are in alignment with this pattern, and we hope that they can similarly inspire a range of follow up studies exploring these effects throughout the field.

Historically, the putative ERP correlates of familiarity and recollection as separable cognitive processes were originally dissociated as cognitive processes of awareness for memory within a single experiment and a single paradigm: an original study in *PNAS* recorded ERPs while participants made Remember/Know/New judgments and concluded that "*These results provide physiological evidence for two types of conscious awareness in episodic memory retrieval*"[14]. This was followed closely thereafter by a separate finding in *Nature*[15] that manipulated shallow and deep encoding of verbal words to reveal dissociable ERP effects we now regard as the FN400 and LPC for familiarity- and recollection-based cognitive processing, respectively. This discovery was then followed up with years' worth of convergent findings from various different paradigms. To further elaborate, a recent meta-analysis review[16] stated about the different ERP correlates of familiarity and recollection (FN400, LPC, respectively) that:

"*These ERP effects have been demonstrated within experiments to be temporally and topographically distinct, providing evidence that they reflect different mnemonic processes* [15,17-20]." That is, individual findings were published within a single experiment/paradigm, which led to drawing conclusions that different effects related to carefully controlled experimental conditions of interest reflected different cognitive mnemonic processes (see also [14,15,21,22]). That standard is effectively criticized here by Reviewer 3, because it is also what we do here in our manuscript (and more rigorously, too, we contend). Our approach is valid, appropriate, and very conservatively couched within the confines of the rigorous standards of the field that have preceded it for using ERPs to characterize differential cognitive processes; furthermore it accomplishes this also within the modern high standards of reproducible science.

The Reviewer also stated that: "*the operational definition of context familiarity is low confidence item memory with correct source memory which reads just as a mixture of item familiarity and recollection*"**.** We appreciate the opportunity to clarify this point: we used that condition to measure context familiarity, but that does not necessarily mean that it is the only definition of context familiarity, just one of many possible constructs for measuring it. That is, in the present study, we used the memory condition of response combinations of low confident recognition and correct source memory as the measurement of instances in which we hypothesized that the otherwise-elusive process of context familiarity could be found to operate. However, we would consider that the process of context familiarity may simply be operationalized as the familiarity of context or contextual information from prior episodic events. We would surmise that, like both item familiarity and recollection-based memory processes, it could be found in future studies using differently defined conditions amid very different paradigms that systematically vary the context while holding items constant (and vice versa; indeed, we cited several fMRI studies that we believe reported similar lines of supporting evidence in our Discussion sections 5.1 and 5.2, supporting our conclusions. We have also added these points for future researchers in a new Discussion section 5.4.4 Future Directions.

But the point the reviewer states about the condition being simply "*a mixture of item familiarity and recollection*" is actually precisely the hypothesis that the original 2012 experiment[3] directly tested, and entirely refuted; and furthermore is also what the present series of experiments also directly tested as a hypothesis and refuted yet again. There is simply no evidence for that position (e.g. of context familiarity condition here being a mixture of recollection and item-familiarity) which nearly all memory models already assume is true for standard recollection responses anyways (i.e. it is widely acknowledged that non-clinical recollection responses are a mix of recollection and familiarity), and from which the present findings were reliably dissociated from. The Reviewer's hypothesis is contrasted to the voluminous extent of evidence we have now provided to the contrary of it. That is, we showed consistently that there was no evidence that the condition was associated with the otherwise-evident neural correlates of either item familiarity nor recollection (and certainly not a mixture of the two), while providing a wealth of evidence to the contrary that the condition of context familiarity instead reflected its own, separate, significant effects (both in behavior, and physiology) representing a distinct third process of episodic memory that cannot be attributed as either item familiarity or recollection- nor a mixture of the two. We hope that may help to clarify, and we have now updated the revised manuscript to make this point more clear, reiterating in Section 5.4.4. that we would expect context familiarity to be measurable in other ways pending the properly sophisticated experimental designs that do not conflate it with recollection or item-familiarity.

Since the Reviewer's next/final point here then mentioned that all three Reviewers raised concerns about the construct of context familiarity vs. the alternative of "a controlled search process" in their initial first round of Reviews, then it is quite important for us to also note here in response that all of those same Reviewers have now, subsequently, agreed that our rebuttal of those original claims of a controlled search process was apparently convincing and compelling, as have now reported having all of their initial alternative interpretations satisfied with our updated, strengthened manuscript. So, we don't find that comment's appeal to their original reviews to be compelling in light of their more updated Reviews that now provided strong support.

**Response to R3.2:**

**If I understand the authors correctly, the argument that there has been absolutely no ERP evidence for recollection for the context familiarity condition means that the ERPs did not resemble a delayed version of the LPC. Actually, I do not think that the ERP correlate of delayed recollection (or a search process) would necessarily look like the LPC as also other processes presumably take place during this later time window and component overlap could alter morphology and topography.**

Yes, we have consistently found meaningful evidence for an absence of recollection-related ERPs contributing to the condition of context familiarity- but more so, too: we also did not see any resemblance to recollection in the patterns of behavioral responding, either. We think that the Reviewer's new account here of '*presumed-recollection not looking like standard-recollection*' is not a viable one to account for the propensity of data: it is effectively resting upon saying that recollection is there but that it is undetectable, even while we are reporting that something else (cFam) is very much significantly detectable (and reproducible). That kind of account is not falsifiable, and it defies data, logic, and parsimony.

The Reviewer goes on to state that "*other processes presumably take place during the later time window…*" but does not say what those "processes" could be. Our account does: we believe it is the evidently independent mnemonic process of context familiarity. The Reviewers theory of 'delayed recollection' is further complicated by problematic aspects: it would predict no differences in ERPs between the condition we call context familiarity and that of correct rejections because if there wasn't memory/processing yet (i.e. it occurred via a 'delayed recollection') then there would be no reason for the condition to differ from a non-memory condition baseline of correct rejections. But of course, that is not what we find in the data several different times in rejecting that null hypothesis: there is in fact a significant difference between the condition of context familiarity and correct rejections, and we need to then know why- or, at least, be able to offer a plausible account to explain why that difference exists. Our account provides at least one viable explanation (it is the dissociable memory process of context familiarity), whereas the delayed recollection account does not provide an explanation (the 'controlled search process' was found not viable in the satisfactory prior round of Reviews.

Additionally, the term ‘delayed recollection’ is a relatively new account for the field of memory research, and appears only sparsely available in the literature, remaining as an otherwise vague construct lacking empirical support. The only report we found on a Pubmed.gov search for “delayed recollection” also linked it with ERPs[44] and found that ‘delayed recollection’ occurred under conditions of changed viewing conditions- which were not a part of the present study. Notably, that study did not find any negative-going ERP effects to be associated with their condition of ‘delayed recollection’, but instead found an opposite pattern of it being associated with positive-going LPC measures. This study hence provides evidence against a ‘delayed recollection’ account, and is thus difficult to support as a viable account of our present findings.  A related ERP study[45] reported a later negative-going difference for hits vs correct rejections, but in their paradigm their presumed LPN effect actually became more diminished in their ‘delayed memory condition’, and was instead larger in their condition of immediate memory retrieval. This is the opposite of what would be predicted from a ‘delayed recollection’ account of the present findings, and thus it, too, ends up providing evidence against the ‘delayed recollection’ account of the present data, and therefore further provides support to the conclusions we draw from the array of present data. We have now updated the revised manuscript to include these and other additional lines of evidence against the delayed recollection hypothesis, in Discussion Section 5.4.2.

**With respect to the behavioral evidence, in my eyes, we would need behavioral responses indicating recollection or (context) familiarity to take them as unambiguous evidence of a differentiation between the two processes. Reaction times can rarely be interpreted unambiguously (see also below). Moreover, the finding that (immediate) recollection responses were faster than the responses in the context familiarity condition would fit quite well with an interpretation as any kind of delayed process**.

We provide clear evidence that behavioral responses for the conditions operationalized in the existing literature as recollection and context familiarity do significantly differ and do so separately for measures of item recognition and source memory.  Notably, context familiarity source memory judgments were reliably faster than recollection, which directly argues against the theory of it being “delayed recollection.” How could it be delayed if the memory response is happening faster?

**I disagree with that delayed recollection necessarily implicates more high confidence source judgments. I think it is possible that a situation where a participant was unsure about their item memory, they would maybe refrain from giving a high confidence source**

**response just because they wanted to be consistent. In other words, uncertainty about whether the item was old or new in the first place, might induce uncertainty about whether the recollected source is correct**.

The reviewer raises an interesting point here, but we are limited in the ability to respond to what may or may not be, due to the speculation of possibilities and "maybes" in this comment since it is guessing about the reasons why certain (or all) subjects make choices about memory judgments (i.e.: they want to be consistent). Participant's various motivations (their surmised desires for response consistency) are beyond the purview of the data when we have no reason nor way to know about participant's internal views and motivations (they were not instructed about that factor) and they likely are quite variable across both different participants and different trials. The theory proposed in the comment would hold that a delayed recollection arrives later in time, even though the source judgments for this particular condition are provided faster than the source judgments in the actual recollection condition. As such, it is difficult for that to be reconciled in light of the pattern of behavioral responses.

It is also possible that uncertainty about item judgments need not be coupled to uncertainty about source in ways like the Reviewer suggested, because people simply could become certain about the source for an item that they first endorsed with low confidence, if that additional source context information comes to them following the first item judgment. Additionally, there is no need for a 1:1 mapping of certainty across responses, since we also see the opposite pattern of many high confidence recognition judgments having low confidence source judgments in the present data.

If recollection is "delayed" to account for the pattern of data we observed, then the common features of recollection should also be evident in at least a reasonable proportion of those response conditions, and those features of recollection very commonly include high confidence responses for source. Indeed, most leading models consider recollection as usually having high confidence responses [23-37], and so this purported theory of 'delayed recollection' cuts across that dominant view. We believe that these possibilities do not hold up well to scrutiny with the available data and should not preclude publication. Rather, they are good ideas best suited for motivating competing studies designed to similarly test competing hypotheses motivated by the present work- which moves the field forward in a positive way. We have now updated the revised manuscript to include these and other additional lines of evidence against the delayed recollection hypothesis, in Discussion Section 5.4.2 and 5.4.4 for Future Directions.

**I also disagree with that participants in the item context condition would necessarily engage in a search process. Why should they if they can give a response based on item familiarity?**

Because they are being asked about their source memory: thus they would engage in a search process for that source memory information if at first it is not available. The evidence for concluding that it is not available comes from the participants, who directly self-report that they do not know the source of the recognized items.

**Response to R3.3:**

**Actually, I think we are all very bad in remembering source information. Turning the above example the other way round: If we would calculate source memory performance the way the authors suggested, we might conclude that we are as good in remembering source information as we are in remembering item information. This would be – as the authors themselves described in the above response – would be an erroneous conclusion**.

We thank the Reviewer for her kind acknowledgement of agreement with our points in the earlier Response to Reviewers about ways for calculating source memory performance. Regarding the statement noted above about being as good at source memory as in recognition memory, there is actually some evidence to support the validity of what the Reviewer is instead claiming here to be an erroneous conclusion (i.e.: that people can be as good at remembering source information as they are at remembering item recognition): for instance, ERP studies of source and item recognition have reported better performance at source memory than for item recognition[38]. Nevertheless, it is somewhat besides the main point here, which is later agreed to by the Reviewer below: that the Reviewer's original suggestion in the first round of Reviews would have been inappropriate for us to use as an alternative way to calculate source memory instead of. the standardized method that we use in the present work to replicate the prior work. We appreciate that kind acknowledgement below.

**More specifically, I am not convinced that the chosen source memory measure is akin to the standard recognition measure of memory performance (%hits - %FA). This measure is used when subjects have to make yes/no discrimination judgments to correct for a response bias. However, it is not calculated for 2AFC displays. As the source memory task is also a 2AFC task, there is no need to correct for false alarms. Usually, percent correct (relative to all trials) is used as the dependent variable. The same logic should apply for a 2AFC source judgment. If source memory is calculated this way, person A with (60% correct, 30% incorrect,10% unknown) is considered better as person B with (10% correct, 0 % incorrect, 90% unknown). If source memory is calculated as suggested by the authors, person B would be considered better than person A only because they were more conservatively responding. In my view, the latter way does not reflect source memory performance for those two participants in a meaningful way. To reconcile the approaches taken by the authors and my concerns, the authors could consider a source memory measure of '%source correct minus %source incorrect' to correct for source misses. In this case, the above example would result in measures of 30% for person A and 10% for person B, which reflects that person A remembered more sources than**

**person B, but also corrected for the fact that they incorrectly guessed in 30% of the cases**.

First, “akin” does not mean “the same”, but means “similar”; so, it should not be confused or mistaken as that we were saying that our method of source memory was “the same” as recognition performance measures but instead that it was (and is) indeed merely “similar”.

Second, in response to the Reviewer’s theoretical vignette of comparing people’s source memory performance against each other: we do not do that kind of comparison of ranking individual participants in this study, but instead we follow the standard and well-accepted statistical procedure of assessing measures of central tendency (average) of the group and do not rank nor compare the scores of different participants. So, the comparative issue among different subjects’ performance that was raised by the Reviewer’s example is not relevant to the present study. Also, our task gives people at least 3 options for source memory (correct, incorrect, and unknown) along a 5-point scale, and thus does not easily fit the Reviewer’s characterization of a 2AFC (two alternative forced choice) response because there are more than just two alternative choices to choose from. Additionally, the Reviewer’s suggested equation to ‘reconcile’ the approaches of measurement does not specify which denominator to use for calculating percentages (i.e.: all total trials, or, only those receiving source judgments), and thus is merely suggesting to do it in her new way and does little to reconcile anything, thereby returning to the same premise that was already acknowledged by the same Reviewer to be in agreement with the Authors about. So, we can’t adopt that novel measure- but, in the updated manuscript we now suggest it as a fruitful path for future research in Section 5.4.4, and give further discussion in Method section 2.3.2, which we believe strengthens the manuscript from the Reviewer’s helpful comment.

Third, and importantly, since this particular project was specifically conducted and submitted as a replication study, it is incumbent upon us to ensure that our measures are replicated precisely to those used in the original findings [3] we sought to assess the reproducibility of: that study used the exact same measure of source memory performance as we do in the current study, has had no problems, issues, complaints, nor dissent about it in the literature in the 12 years since then and amid its reasonably well-cited breadth of impact in the field (173 citations). Therefore, we believe it would be inappropriate for us to change calculation methods at this stage, and then seek to make claims about reproducibility while fundamentally changing from the approach used in the original findings[3]. In fact, doing what the Reviewer suggests (changing calculation methods post-hoc) would (or at least, could) be problematic for publication due to the post-hoc changing of the apriori-defined measures for reproducibility. It would also reproduce nothing, since it would be a new thing instead of testing an old thing.

Furthermore, our approach is a common one that is sensibly used in source memory studies for the reasons we elucidated in the prior response to reviewers and which was subsequently incorporated into the Method section of our manuscript. For example, our standard approach used to measure source memory performance is not just the same as we have published with before and which has remain uncontested in the literature for 12 years [3] but is also the same approach used by others for measuring source memory performance[38], where they note that: "*Conditional source measures control for overall differences in recognition by calculating the probability that the correct source is identified given that the item was correctly identified as old (p|source| = source hits/source hits + source misattributions)[39,40]*". This has been added to the revised manuscript Method section 2.3.3 about source memory measurement.

We have now updated the manuscript to incorporate these additional lines of support and justification for the procedures used in the current studies. The broader field of memory science is free to quibble with different ways of measuring source memory, the relative advantages and disadvantages, etc. and determine publicly as a field what may or may not be the best approaches and how that may or may not differ among different experimental situations. But we believe that process should be best accomplished within and across the literature of many scientists in public discourse, rather than in the current setting pre-publication, since such nuanced opinions are sure to vary across both scientists and experimental conditions. Our approach in the present work remains as a well-established standard one (used by many other established studies) that is well-suited for comparisons to results from other studies, is a principled one (well-justified in logical rationale) that is well-suited for our particular experimental paradigm, and is also a well-premised one (required to be the same as the study being tested for the high standards of rigor in reproducibility).

**I agree that setting chance level to 50% and not excluding the "don't know" responses might not reflect what we really want to know, i.e. whether the subject guessed in case they made a response. To be clear, I agree here with the authors because I agree with their argument concerning variations in the bias to use the "unknown" response. This is, however, the same reason why I think it is not appropriate to calculate source memory (only) in the way suggested**.

We again thank the Reviewer for her kind acknowledgement of agreement with our earlier points.

**Response to R3.4:**

**Although close replications are always commendable, I think that the alignment between different measures within the current study is more important. If I understood correctly, ERPs indicating context familiarity and recollection were both calculated across source memory confidence levels (see Figure 1). Therefore, I would suggest to reanalyze the RT data correspondingly. As both levels of source confidence will be used for context familiarity and recollection, there should be no confound**.

The Reviewer raises a very good point, and we very much appreciate the opportunity to clarify the reasoning and justification for the measures used in the present study. The reason why the behavioral measures of response times only included the low confidence source memory judgments was two-fold: first, because it ensured that they were not confounded by source memory confidence, and only varied on the factor of recognition confidence and thus were effectively matched in source memory strength; and second, because it was the first time this analysis has ever been performed, so it was critical to ensure the strictest of rigor in holding variables constant. Indeed, this is the identical approach as was taken in the original 2012 paper's first characterization of the ERP effects for the same conditions [3] (Experiment 1, 2012). Subsequent follow-up ERP investigations (Experiment 2, 2012) then broadened the inclusion criteria for the ERP analysis to also include the high confidence source memory judgments; that is why we felt it reasonable to replicate that here for the ERPs, not RTs, since that approach also permitted inclusion of more trials and hence more subjects for a larger sample size, SNR, and statistical power. But in both of the analyses performing 'first-time' assessments of conditions (2012 ERPs, present study RTs), we believe it strongly merits ensuring that the prime findings cannot be attributable to any other possible confounds or conditions; thus requiring the approach we have adopted here, which we believe is both rigorous and reasonable.

Regarding the stated belief that there would not be a confound in including high and low memory confidence in the source memory RT responses, we note that the high confidence recognition response (5, recollection) also had much more high confidence source memory responses, whereas the low confidence recognition responses (4, context familiarity) had far less (nearly none) high confidence source memory responses, and so there very much remained the potential for confounding of the conditions since they would vary on two dimensions of both item recognition confidence and source memory confidence. As described above, we sought to minimize that potential confound given that it was the first behavioral exploration into response times of these sensitive and important conditions.

Nevertheless, if it may ease the Reviewer's concerns, when both high- and low-confidence source memory responses are included together in the conditions we used for context familiarity and recollection, the results for response times remained the same: context familiarity was significantly slower than item familiarity ($t(40) = 2.38$, $p = .022$, Cohen's $d = .372$, 95% CI [42.70 524.6]; cFam: $M = 2512$, $SD = 778$, $SE = 121$, iFam: $M = 2229$, $SD = 677$, $SE = 106$), and also significantly slower than recollection ($t(40) = 7.66$, $p < .001$, Cohen's $d = 1.20$, 95% CI [-1034 -602]; Rec.: $M = 1695$, $SD = 301$, $SE = 47$)- but again, the findings of these particular conditions also include differential levels of memory confidence, and are not matched for being either the same item or source memory judgments; therefore we feel it is most appropriate to constrain the behavioral response time data to the matched levels of source memory confidence since confidence would be more likely to affect response time than covert measures of memory (ERPs).

As a final note, even if the inclusion of higher source confidence responses were to affect ERP effects, then it would do so in the direction of a stronger test of our hypothesis, such that the inclusion of higher confidence trials would make the context familiarity condition more likely to exhibit positive ERP effects associated with recollection [or, ‘delayed recollection], since confidence is known to exhibit positive posterior scalp activity [11,20,41]- which of course none of our extensive results detected any evidence for. Thus, we feel the data strongly supports our conclusions.

**Response to R3.5**

**I don’t find the argument that the context familiarity condition differs in RT and ERP measures from recollection very convincing as it could do so because (as the authors claim) context familiarity and recollection differ in their temporal and ERP characteristics, but it could also do so because late recollection and immediate recollection differ in their temporal and ERP characteristics. Moreover, I am not sure whether this late recollection or memory search process should necessarily result in high confidence source judgments. As the successful memory search was preceded by low confidence item memory, participants might be unsure whether they can trust their search results (see also my comment to the response to point 3.2)**.

We respect the perspective of the Reviewer about interpretation of differences in RTs and ERP effects. However, the fact remains that the two conditions (recollection, context familiarity) do indisputably differ in measures of both behavioral RTs and physiological ERPs. Both of those differences are meaningful, and warrant communicating to our scientific community. The alternative possible interpretation provided by the Reviewer (i.e.: that each measure of conditions differ because there might be early and late recollection characteristics) remains without evidence across our five different data sets, and so is by comparison a weaker argument.

The Reviewer is also implicitly implying or directly suggesting that Recollection should or could be associated with low confidence source memory judgments, which is generally inconsistent with how the process of recollection is typically modeled to operate (i.e. which is as a threshold process usually constrained to high confidence responses [25,30,31,36]). If, as the reviewer now claims here, that a participant deduces that they are unsure if they can trust their unconfident-but-accurate source memory response, then we would contend that this is effectively describing what we also describe as familiarity of the context. In that way, the Reviewer is, effectively, making our same point.

As was noted above in response to R3.2: The Reviewers theory of ‘delayed recollection’ is further complicated by problematic aspects: it would predict no differences in ERPs between the condition we call context familiarity and that of correct rejections because if there wasn't memory/processing yet (i.e. it occurred via a 'delayed recollection') then there would be no reason for the condition to differ from a non-memory condition baseline of correct rejections. But of course, that is not what we find in the data several different times in rejecting that null hypothesis: there is in fact a significant difference between the condition of context familiarity and correct rejections, and we need to then know why- or, at least, be able to offer a plausible account to explain why that difference exists. Our account provides at least one viable explanation (it is the dissociable memory process of context familiarity), whereas the delayed recollection account does not provide an explanation (the 'controlled search process' was found not viable in the satisfactory prior round of Reviews). We have updated the Discussion section about Delayed Recollection accordingly (Section 5.4.2)

Furthermore, to the Reviewer's point of the context familiarity condition being low-confident correct source memory judgments, we also point out here that our prior work in 2012 verified that the same results are evident when including the high-confident source correct responses (2012, Experiment 2)[3]. This weighs against the Reviewer's claim of being unsure that 'late recollection' should result in high confidence responses, and that people might be unsure about if they can trust their memory search results; as such it weighs in favor of the account we continue to hold (see also the details in our final response above to R3.4). We hold this account of the findings because they are premised in consistent convergent findings across several different studies and methodologies, while the alternative account proposed here by the Reviewer appears to instead rest upon being "unsure" about what only theoretically "could" and "might" be happening in the absence of any supporting evidence. We hope that the present findings will motivate additional studies that can test these conclusions against new evidence that might emerge in the scientific process.

**Response to R3.6:**

**I still find the explanation given in the manuscript a bit short. Especially, as during the course of the report, the time window changes back and forth and it is hard to follow why which time window was choses for a specific analysis. Specifically, it is not clear to me why in section 3.5 for the control analysis with the lower trial number criterion the 400-600 time window was used while in the original analysis in the same section the earlier time window was used. And why did the time windows between the between-subjects analyses for Experiment 1 and 2 differ? For reasons of clarity and readability, I would suggest to choose only one of the time windows (and electrode sites) for the main manuscript, and whenever the authors want to demonstrate that the effect can also be found in the other time window, they add this information in the supplemental materials**.

The Reviewer makes a good point, and the reasons and rationale for the time windows used are provided in the updated manuscript. In short, In Experiment 1, the within-subjects direct assessment of familiarity conditions focused on the time window of 300 to 500 ms (Section 3.5) because "we investigated if the two conditions varied during the conventional time window traditionally reported for familiarity-based processing of the FN400", and thus the between-subjects analysis of that data set also used the same time interval of 300-500 msec. Later, in Experiment 2 (Section 4.3.4), it drew upon data from existing data sets in which the FN400 effect had been previously well-characterized, thus "as familiarity effects for these prior studies were reported for 400 to 600 ms, a targeted analysis was conducted during this latency". Given how commonplace these two time intervals are in the literature, and the very near overlap of them, we considered it a negligible difference and principally based upon the known priors of the preceding work.

**Response to R3.8:**

**The reason why I related the ERP findings to the response times was the following: Generally, we might assume that the search process reflected in the LPN/BCN is initiated whenever other information was not informative for a successful memory judgment. Further, if participants respond after the information retrieved by the search process putatively reflected in the LPN is available, the process presumably contributes to successful memory judgments. In contrast, when participants – for any reasons – respond before the information of this search process is available, the process cannot contribute to a successful judgment. In addition, I don't think that just saying that others took the same view before is not a very good argument in itself. Therefore, I would remove this from the Discussion again**.

No, we generally do not assume that the effects reflect a controlled search process nor the LPN. This was otherwise successfully resolved in the prior round of reviews with everyone else, as attested to by the other two Reviewers reporting that their concerns have now been well-satisfied by our exhaustive response to them in that updated manuscript- which we still believe holds up well. Beyond that, it is a little difficult to decipher the proposed account described in the Reviewer's comment here about what might (or might not) happen. But in general, it seems to be saying that the LPN happens when a memory search happens and people respond after the success of the memory search (i.e. they remember); whereas it then posits that people respond to a memory test's question "before" the memory search information is available, and thus is not a successful response. The relation of that vague account to the present findings is presently unclear to us, though we would be glad to respond if more clarity of details could be provided to help us better understand.

The reason that the updated manuscript had included citations and quotations of what other leading scientists have attested to (i.e. the authoritative Review paper on the LPN stating that the LPN was not the same as our present findings we here refer to as the BCN), was entirely because it was directly responsive to what all 3 Reviewers had raised as an initial alternative interpretation of our findings. That approach of citing what other authorities on the topic have taken as the same view as ours was evidently effective and compelling at convincing the other two Reviewers who reported now being satisfied with a much strengthened paper. So, we opt to retain that section in the Discussion in assuming that it may similarly help other future readers to better understand the findings, unless the Editor's discretion indicates otherwise (in which case we are glad to adjust).

**Response to R3.9:**

**The LPN is usually evoked when there is a high need to search for source information (see citation above). However, during the item memory judgment there was no need to search for source information when a response could be given based on item familiarity and recollection. Faster response times for these two conditions are in line with the interpretation that participants had all of the information to make a response more readily available than in the context familiarity condition. Possibly, when prompted for source memory in the next display, they resumed a search process resulting in longer response times. However, this cannot be visible in the ERPs time-locked to the item recognition prompt**.

If the Reviewer is suggesting that what we identify as context familiarity responses could have been made based upon recollection, then this does not make sense to us since we instead provide evidence of absence (Bayes Factor analyses of null findings) for the well-known physiological putative correlates of recollection during those memory judgments being speculated upon here, and that finding is replicated again and again across our data sets from various different studies. The suggestion lacks evidence, whereas our account is based upon the evidence provided.

Regarding the comment about time-locked ERPs to account for longer RTs in certain conditions, one strength of the present findings is that they rely upon the standardized putative neural correlates of memory processes well-established in the field over several decades; these standard correlates in the field have been found as time-locked measures to the stimulus onset (and not response locked), and so the present study preserved that critical foundation from which to compare and interpret the present findings. We agree that response locked analyses could be very interesting for future studies to look at, but also note that they risk being confounded or at least conflated in their baseline period due to the preceding activity of the item recognition prompt/decision in the present paradigm- so future studies would need to be carefully designed to ensure those factors are disambiguated. We have included this good suggested by the Reviewerer into an updated section of the manuscript's Discussion Section 5.4.4: Future Directions.

Overall, the Reviewer's account continues to hinge upon speculation of what might or might not have happened for hypothetical participant's mental decision processing, and lacks evidence that can confront the array of evidence that we alternatively continue to provide across the studies and measures. Furthermore, the Reviewer's account of RTs still ignores the fact that the RTs for context familiarity were dissociable from the conditions of item familiarity and recollection across both item and source memory judgments, and does not offer a reason why that difference exists, nor how that pattern of differences an cohesively be accounted for the parallel observations of dissociations in ERP measures of those same conditions. Our interpretation and conclusions continue to put forth an account of the findings, and a formalized model, that can and does account for these findings, and thus we believe that it weighs in more favor than the suggested accounts that rely upon speculation of “possibilities” in the absence of supporting and integrative evidence.

**Response to R3.10:**

**I did not mean that we could expect the LPN any time or anywhere. With respect to timing: The authors themselves are reviewing in the Discussion (4.2.5.1) that the early variant of the LPN occurs approximately 600 to 1200 ms post-stimulus which exactly covers the time window of the BCN in the current study. That leaves only the topography as diverging from standard findings which might fall into the natural variability the authors are willing to accept, especially as they accepted this natural variability also for the FN400 which was found more centrally than in the classic FN400 studies. Besides the absolute timing and topography, I consider it as important to also take into account the morphology and the relative timing. So, as I see it, the LPN usually occurs directly after the positive component associated with recollection. Whether the LPN is immediately differentiating between conditions, takes more time to unfold or is extended over a longer time period is presumably a matter of exact task conditions. With respect to the relative timing, the BCN exhibits the same characteristics as the LPN. As a final note regarding the LPN interpretation, I don’t think it is so important whether the effect reported here resembles the LPN in previous studies or not, and whether all predictions for the LPN are fulfilled. It is possible that the BCN effect is very specific to the current task demands and therefore does not resemble the LPN. However, the eliciting condition leaves open the interpretation that the effect reflects a successful search process for context information that results in uncertain responses**.

We respectfully disagree, for the reasons already noted in the first Response to Reviewers and which was integrated into the updated manuscript that both of the other two Reviewers also found to be compelling and satisfactory to their noted similar observations.

The Reviewer stated: “***as I see it, the LPN usually occurs directly after the positive component associated with recollection*”,** but that is precisely what was *not* found in the present studies. Instead, those studies never found the negative ERP effects after a positive component associated with recollection- and these null findings were quantified with Bayes Factor analyses, demonstrating meaningful evidence of absence.

The Reviewer then goes on to state that “***I don’t think it is so important whether the effect reported here resembles the LPN in previous studies or not, and whether all predictions for the LPN are fulfilled*”** and that “***It is possible that the BCN effect is very specific to the current task demands and therefore does not resemble the LPN*”.** We do not find these to be compelling arguments in support of a framework that can attribute the present findings. They are not compelling because their account is, in effect, not falsifiable, and thus not a very good scientific model to rely upon if other options of potential interpretations exist which account for the pattern of data more parsimoniously and without contingencies of what “might” be “possible”. The adaptations we provide to the BIC model (Fig. 8) can directly account for the findings more parsimoniously [33,42], and which is couched in a much broader (and consistent) literature which we copiously review in our Discussion thereof.

The continued discussion here is a valuable one, but at this point it continues to represent an untested theoretical position on the one hand (the Reviewer) vs. a position based upon the present data on the other hand (the authors). We believe this is a worthwhile discussion to be had in the public literature where healthy scientific dissent flourishes around findings published for all to evaluate, and where it can be evaluated against comparative studies that can be conducted to provide data to inform the theoretical positions purported here. That is, the Reviewer's point here makes a better case for publication because of how it inspires future work that can resolve ideas that can be tested by dissenting views.

**Response to R3.j:**

**I'm not convinced that in this specific case, pictures and faces could be seen as contexts, as they were presented alone. Isn't a more parsimonious explanation for the patient NB data that the sparing of the right perirhinal cortex (and maybe also face processing related areas) left item familiarity for pictures and faces intact?**

We appreciate the Reviewer's insight about this topic and have updated this section again to better incorporate this good suggestion. We think that, ultimately, both views on this topic of interpretation are inconclusive due to lack of data on this rare perirhinal cortex lesion patient (NB), and thus equally meritorious possibilities. In respect to parsimony, we are reluctant to assume laterality equivalents across brain hemisphere regions due to oversimplification that it suggests, and that it is well known that there can be unilateral functions that do not always clearly delineate. Nevertheless, we have updated the revised manuscript to reflect this good suggestion from the Reviewer as stated below and Section 5.5: Implications for models of memory:

> "*This patient was deficient in familiarity for items (words) but showed intact abilities of familiarity for pictures and faces, which we consider here as broadly representing kinds of context in episodic events, and this could be seen as performance consistent with the patient's intact parahippocampus service of context familiarity (though it is possible that faces and/or picture stimuli might have potentially instead be treated by that patient as an item in certain studies with other contexts, or alternatively the face/picture memory performance might have potentially been supported by the other hemisphere's intact perirhinal cortex).*"

**Additional points, which came up during reviewing the revised manuscript:**

**Although control analyses are of importance, I don't think they add much value if they are terribly underpowered, such as N=3. Moreover, using parametric test statistics seems inappropriate to me in those cases**.

The Reviewer raises a good point. The analysis noted here (Section 3.5, Experiment 1) was included at the advice and request of Reviewer 1 (R1.4 in the previous Response to Reviewers) and the post-hoc power analyses of these findings indicated a quite-high power level of .952, which led that Reviewer to report that "the revised manuscript has been substantially strengthened and authors have addressed all of my concerns". That is one reason why we are inclined to keep that section included, in case it may similarly help any other readers address potential questions. Furthermore, Section 4.3.3 for Experiment 2 also then followed upon that initial small sample size from Experiment 1 and independently reproduced it in a separate data set that was four times as large (N = 12), which we hope may assuage any lingering concerns.

**The authors added a very detailed discussion on why recollection responses were faster than the other conditions. The simplest explanation, in my eyes, however, is lacking. Isn't it very plausible that responses with high confidence are made faster than those with lower confidence? Moreover, although familiarity is seen as a faster process than recollection, it just means that the information retrieved by these two processes is available earlier or later, respectively. Whether responses are made faster or slower for one or the process demands very much on task instructions**.

The Reviewer makes a good point. However, that account does not explain why conditions of item- and context-familiarity were found to differ in RTs, since even when the higher confidence responses were removed from the item-familiarity condition, in order to ensure that both conditions were matched in low-confidence recognition levels, the results remained the same (i.e.: they reliably differed in RTs). This point was raised in point R1.4 of the previous round of Reviews and was incorporated into the text of that updated manuscript, where it remains in section 3.5, and then replicated again in section 4.3.3 of Experiment 2. Another complication to the account proposed by the Reviewer is that the response conditions are not just single conditions of confidence but are combined conditions of both recognition confidence and source memory retrieval. Finally, the task instructions were the same for all conditions in the present studies.

**I was not sure whether in the discussion on absolute versus relative familiarity, these processes were described as being qualitatively or quantitatively different. While they are both operating on item information, I think they are still qualitatively different. Moreover, it is not clear to me why absolute familiarity should play a role in the experiment reported here as pre-experimentally known stimuli were used for both old and new items, so only relative familiarity would be diagnostic**.

The reason why this point was suggested in the Discussion as a possible account of the spatial distribution of the FN400 effect observed here was because there could still have been differences in some stimuli for their absolute familiarity from pre-experimental exposure of some words relative to others used in the stimuli list. Admittedly, the study was not designed to assess nor compare factors of pre-experimental familiarity for absolute and relative familiarity, and so we are not able to gauge that factor but merely mention it as a possible and plausible account of the observed findings for the spatial distribution of FN400 effects, which are comparatively the weakest and most variable part of the ERP method[43].

**In the discussion on the LPN, the authors implicate that if the BCN would be an LPN, there should also be a late right frontal old/new effect. It would be easier to follow the discussion, it the authors would explain why these two components are necessarily linked together?**

The Reviewer raises a good point here. The reason that we mentioned the possibility of there also being a late right frontal old/new effect to be predicted in accord with the LPN, is because the late right effect is reported to be linked with post-retrieval monitoring that one might think could be associated with memory search processes (assuming an LPN-linked memory search process was occurring). Upon reflection from the Reviewer's comment here, we agree that this was not necessary, and this sentence of the Discussion has now been removed in the revised manuscript. We thank the Reviewer for the thoughtful point here, which improves the paper.

**Minor:**

**p.6. 'Image-ability'** - updated
**p. 7.' what the response criterion'** - updated
**p. 8. How many items per condition entered the ERP analysis?** - updated, Section 2.3.1
**p. 8 section heading 2.6.1: delete 'Reproducibility'** - updated
**p.10 'an significance criterion of a = .05'; change 'an' to 'a' and 'a' to 'alpha'** - updated
**p.10 '(.486) in the in the'** - updated
**p. 10 'the between-subject effects' delete s** - updated
**p.11 reference to 'Supplementary Table 2': the table is now in the main text** - updated
**p.11 Which response types were included for which condition in RT analysis is now described a bit less precise as in previous version. I suggest changing again.** -updated
**p. 16 'Experient'** - updated
**p.19 'is generally a slower process than.'** - updated
**p.20 'to better powered'** - updated
**p. 20 'which might seem odds'** - updated
**p. 21 'suggest the important'** - updated

We thank the Reviewer for these very kind suggestions of earlier typos & errors which have now been revised in the updated manuscript.

**Regine Bader (signed)**

**Reviewer #1 comment on Reviewer #3:**

**I agree with R3.1 that the conditions are not process pure, such that the observed effects may be a combination of item familiarity and recollection. In theory, if there is a way in which context familiarity could be measured independently from both item familiarity and recollection, such an experiment would be highly informative. However, said design may not be feasible.**

We agree with the Reviewers here that future experiments would be highly informative, and we hope that the present work serves to motivate conducting those studies. That being said, we believe that our ERPs and RT measures have indeed succeeded in providing at least one way in which “context familiarity could be measured independently from both item familiarity and recollection”. Each condition demonstrates mutually exclusive effects: they are dissociable temporally and occurring in differential locations on the scalp, and demonstrated as such in various different measures (hits v correct rejections, item familiarity vs context familiarity, etc.). Please also see components of our earlier responses to R3.0, and R3.1 for further detail and discussion.

**I agree with R3.2/R3.5 that the observed findings could be described as "delayed recollection," rather than context familiarity and that interpretations based on RTs are limited.**

Please see response above to R3.2 and R3.5 for extensive discussion.

This theory of delayed recollection is difficult to square when one thinks about it more deeply to its natural consequent end. That is, first, it is saying (without evidence) that their account of present evidence is that something else is happening later (i.e.: after 1200 msec) which is not measurable during the time that significant effects are nevertheless found (i.e. for context familiarity, 800-1200 msec), and using the non-measurable effect to explain the measured effect. It is, in effect, saying that recollection might be present later in time if only we were to look.

But the problem with that is that it is improperly assuming a direct 1:1 mapping of cognitive processes to ERP effects, de facto assuming that the timing of one effect (ERPs) must coincide with the timing of another effect (the cognitive process). This of course was discussed and rebutted at length in the prior round of Reviews that the Reviewers reported finding satisfactory and strengthening to the paper, and which is also discussed at length in the manuscript’s Discussion itself. Said again here succinctly: ERP effects are covert measures widely acknowledged to occur separate to the overt subjective experience of conscious memory reported by the subject[43]. Thus, it requires acknowledging that both item-familiarity and recollection are both very likely also (and normatively) occurring in a delayed manner evident in their reported response times (2.4 seconds, 1.8 seconds, respectively)  that is certainly preceded in time by the temporal precision identified for their respective ERP correlates (e.g. ~400 msec, ~800 msec, respectively).

Therefore, the Reviewer's theory of "delayed recollection" then rests tenuously upon the foundationally-incorrect proposal that only processes measured in our condition of context familiarity is delayed (in theory, 'delayed recollection'), ignoring the widely agreed-upon fact in ERP research that all of the cognitive processes of memory are likely occurring in a delayed fashion from their initial physiological 'pre'-processing via their putative ERP correlates that we generally and simply take as granted in the field.

From this, there is still no evidence-based explanation of what the significant effect of the BCN is in the present series of studies (we have extensively ruled out the 'controlled search' hypothesis as found satisfactory by both other Reviewers, nor why it would not also show the preliminary electrophysiological correlates of recollection that the other, specified condition of recollection did show evidence of  (that also occurred later in time, mean RT of 1.8 seconds). That is, all recollection is delayed, at least in the sense that the traditional recollection condition's responses are clearly and normatively delayed from the earlier ERPs that are well-established as their putative neural correlates. Why would it then, now be something different and special for this condition of accurately retrieving the source- especially when considering that the context-familiarity source memory responses are notably *faster* (e.g. happening less delayed) than both recollection and item-familiarity? We find this theory very difficult to maintain as a viable account in the face of this logic, the available data, and the absence of any countervailing data. It lacks parsimony and logic and becomes its own "magic bullet theory" that, if taken to its natural conclusions, could account for anything post-hoc in the absence of evidence.

If we are wrong (and we don't think we are, since we have tried extensive efforts to rigorously disprove our position), it is only determinable by independent data from other experiments to test & show that we are wrong. Those studies would only be motivated and relevant if refuting a published series of evidence, not a hypothetical finding from an unpublished manuscript). We would applaud and support that heartily. That is, it is a fantastic opportunity for the scientific process to work.

**The authors should calculate source memory in the manner suggested in R3.3**.

Please see response above to R3.3
