## Supplementary material for "A third kind of episodic memory: Context familiarity is distinct from item familiarity and recollection": First Response to Reviewers at NatureCommsPsych

FLORIDA TECH
College of Psychology
and Liberal Arts

November 28, 2023

RE: Re-Consideration of Manuscript ID# COMMSPSYCHOL-23-0270A-Z

Dear Dr. Schiffer,

Hello. Thank you for the kind re-consideration of our manuscript, *A third process of episodic memory: Context familiarity is distinct from item familiarity and recollection* (ID# COMMSPSYCHOL-23-0270A-Z). We also thank the Reviewers for being generally positive, with constructive comments and suggested analyses. They raised several good points, which has substantively improved the manuscript's findings.

We have now included a revised manuscript, comprehensively updated to be responsive to each of the points noted by the Editor and by those thoughtfully raised by the Reviewers. The revision now includes additional analyses that provided a full replication among three independent datasets (previously published in *PNAS, Neuroimage, Neuropsychologia*), as well as the sensitivity analysis responsive to a Reviewer's good suggestion. We hope they will be pleased to see that our findings are preserved when conducting their suggested analyses.

We have done our best to ensure compliance with all of the journal format requirements, as well as statistical reporting guidelines that were requested, and we would be glad to make any other changes that may be requested.

Very Sincerely,

Richard J. Addante, Ph.D.
Associate Professor of Psychology; NIH LRP Fellow

Florida Institute of Technology
150 W. University Blvd.
Melbourne, FL 32901-6975
321-674-8104
floridatech.edu/copla

REVIEWER COMMENTS:

**Reviewer #1 (Remarks to the Author)**:
**In this study, Addante and colleagues tested the hypothesis that there is a third memory process, context familiarity, that is distinct from both item familiarity and recollection. Participants studied words and performed a living/man-made orienting task on each word. Participants' item and source memory was tested for each word. Scalp EEG was recorded throughout. Test phase responses were coded as: recollection (high confidence hit + accurate source), context familiarity (low confidence hit + accurate source), and item familiarity (high and low confidence hits + source unknown). Both reaction times and electrophysiological responses dissociated the three response conditions. Responses during item judgments are slowest and during source judgments are fastest for the context familiarity condition compared to recollected and item familiarity conditions. Electrophysiologically, context familiarity is characterized by a more negative going FN400 at central sites relative to item familiarity. The authors conclude that their results suggest the presence of a third memory retrieval process, context familiarity, that is distinct from both recollection and item familiarity. The results reported would be of interest to those in the field and the direct replication of prior work is a major strength. A number of control analyses rule out alternative accounts of the findings. Some direct tests are needed to support claims made and some interpretations seem to go beyond the data. Some information is missing or lacks clarity. I outline these concerns below.**

> We thank the Reviewer for taking the time to provide a thoughtful, insightful, and helpful review, as well as for both supportive and constructive comments that we believe have led to valuable improvements to the manuscript. We have revised the manuscript in consideration of the constructive comments, adding several direct tests as requested to support the claims, and constraining the interpretations to the capabilities of the data, while improving clarity as well. We hope that the Reviewer may find it satisfactory, and again, we are grateful for the time, effort, and insightful suggestions.

**R1.1 The authors make some statements that require direct tests, as a significant effect in one time interval or over one location does not mean that the effect is significantly larger/more robust than another time interval/location in which an effect is not significant e.g.: "*indicating a temporal specificity to the FN400 latency*" - requires a direct test of the time interval of interest vs. other time intervals tested. E.g.: "*A third point of differentiation between the LCN and LPN is topographical: the LPN is generally found in posterior parietal areas, whereas the current effect was observed in superior central regions of the scalp and distributed to both frontal and central-parietal areas. A fourth distinction is the timing: the LPN tends to begin during periods after the epoch in the present study ended (1200 msec)*." - requires direct test of voltage dissociations either across areas/electrodes or time intervals.**

Regarding the first note from the reviewer citing the need for a direct test of time intervals: this is a great point, and we agree that we had originally written this section to be rather clunky. To simplify the manuscript, the updated revision removes the statement of temporal specificity, and has been improved to be more computationally efficient in reporting the core findings.

Regarding the comment that LCN and LPN topographic differences "*requires direct test of voltage dissociations either across areas/electrodes or time intervals*": We agree that the statement from the original manuscript was a bit unclear (though now has been revised for clarity), but it was merely referring to effects of the LPN reported by other studies in the literature (not a dissociation found in the current study). For instance, the current effects replicated the prior LCN findings of 2012, which had been reported then for topographically identifying effects to have a broad distribution across the scalp with frontal and central maxima, as opposed to the specific parietal regions usually reported for the LPN in other studies. Indeed, the authors of the leading LPN review (Mecklinger et al., 2016) state that the two effects (LPN, LCN) should not be confused for each other, as we also have agreed in our own previously published remarks (2012) (*for direct quotations please see response to Reviewer Comment R3.10 below*). The manuscript's point here was simply to note, and contrast, that the different ERP effects have been previously reported in the literature to be occurring in different time windows and scalp sites, which has weight for making reasonable interpretations about the conclusions from the data. The updated version of the manuscript has re-written these sections to make that aspect more clear.

Since (for reasons we elaborated upon in Supplemental Discussion and in response to Reviewers below) we do not believe the current effects (800 to1200 msec at central site Cz) are those of the later onsetting LPN (e.g., 1200 to 1800 msec at posterior parietal sites; Mecklinger, 2007), it is not possible to create a direct comparison of those effects. For instance, our recording epoch for ERPs was limited to stopping at approximately 1200 msec post-stimulus onset - which is the time the LPN is often found to begin. We cannot extend our analytic time interval further into the later epochs used in other studies because those times are when different the memory motor (and cognitive) responses are occurring for item and source memory judgments, which introduces other potential artifacts, and also raises potential problems of varying baseline period activities across both trials and subjects (see also the responses to Reviewer 2.4 below).

**R1.2. The authors make multiple claims in the discussion that their results support the interpretation that they are observing processing differences rather than representation differences (e.g. "not simply separate representations of memory features or information types"). It is unclear how the present results can adjudicate these two accounts or why this distinction is important to the central conclusion**.

The Reviewer makes a good point here. We agree with the Reviewer that such statements may not be necessary for the scope of the current article, and have removed it from the updated manuscript.

**R1.3. It is not clear which onset is used for the ERP analyses, the onset of the item judgment or the onset of the source judgment or whether both onsets are used. If one onset is used for all analyses, the motivation for this is unclear**.

We apologize for being unclear on this point in the previous manuscript, and the revised manuscript has now been revised to include these valuable details of the study. The ERP epoch was measured from the onset of the memory probe item, not the source judgment. This was the same as in the original 2012 paper that reported the same results we sought to replicate. The recording epoch for ERPs was limited to approximately 1200 msec post-stimulus onset, as after that, the motor responses for item and source judgments are occurring, which could introduce potential artifacts of motor and cognitive differences beyond the initial memory processing of the item that the present investigation was premised upon. The updated manuscript also makes these details more clear in the supplemental methods section on electrophysiological analyses.

**R1.4. Item familiarity is defined as high and low confidence hits (5,4 item judgment responses) and source unknown (3 source judgment responses). When compared to the context familiarity condition (low confidence hits 4 item judgment responses and source correct, 4 and 5 source judgment responses), this presents two rather than one clear difference between the conditions. I am concerned that the inclusion of the 5 item judgment responses/high confidence hits for the item familiarity condition is contaminating the results. For example, the inclusion of 5 responses could be dragging down the "iFam" reaction times in Figure 2 Item Memory. If the item familiarity condition is restricted to low confidence hits (4 item judgment response) are the behavioral and electrophysiological findings comparable?**

The Reviewer makes a fantastic point here, and we are grateful for it. ***Succinctly: Yes, the results remained the same, and no, the inclusion of the item5 responses are not contaminating the results with any confounds - other than simply reducing the power to detect differences in the item familiarity measure***. The revised manuscript has now been updated to include this valuable addition as detailed below, and we thank the Reviewer very much for the helpful contribution.

We pursued the analysis suggested by the reviewer and it revealed that there was still a significant difference in response times for item familiarity [‘item 4 judgments without source memory’] and context familiarity [‘item 4 judgments with correct source memory’] ($t(36) = 2.09$, $p = .044$, Cohen’s $d = .344$) during item recognition, whereby item familiarity became slower than context familiarity. A marginal effect was maintained for the source memory judgment, such that item familiarity remained slower than context familiarity ($t(36) = 1.94$, $p = .060$, Cohen's $d = .155$). This finding has now been added to the revised manuscript in the main Results reporting behavioral findings.

ERPs for this contrast were available from a limited sub-sample of the primary dataset due to there only being N =3 participants available meeting our strict, predetermined inclusion criteria for analysis of the 'item4+source unknown' item familiarity condition vs context familiarity (see Methods; n = 12 artifact free ERP trials in each condition compared pairwise). We pursued the Reviewer's suggestion to assess ERPs for item4+source unknown (item familiarity.) vs. item4+source correct (context familiarity), using a directional t-test based upon the hypothesis that context familiarity exhibits more negative-going ERPs than item familiarity at the same mid-central size (Cz) and time (300 to 500 msec) that was found in the primary analysis of the manuscript ('item 4&5 + source unknown' vs. 'item4+source correct').

Despite the limited size of the available for this analysis, results remained the same, in that ERPs for context familiarity were more negative than those for item familiarity, though this difference was not significant from 300-500 msec at Cz with this limited sample size ($t(2) = 1.67$, $p = .119$, one-tailed, Bayes Factor = 1.64; cFam: $M = -3.59$, $SD = 2.19$, $SE = 1.27$; $iFam_{Low}$: -1.678, $SD = 3.99$, $SE = 2.30$). However, there was a significant effect maintained in the adjacent time interval of 400-600 msec that has been used for measuring the FN400 in prior studies using this dataset (Miuller, Sirianna, & Addante, 2021): ($t(2) = 5.57$, $p_{bonf} = .015$, one-tailed, Cohen's $d = 3.21$; cFam: $M = -4.51$, $SD = 1.54$, $SE = .89$; $iFam_{Low}$: -1.36, $SD = 2.39$, $SE = 1.38$).

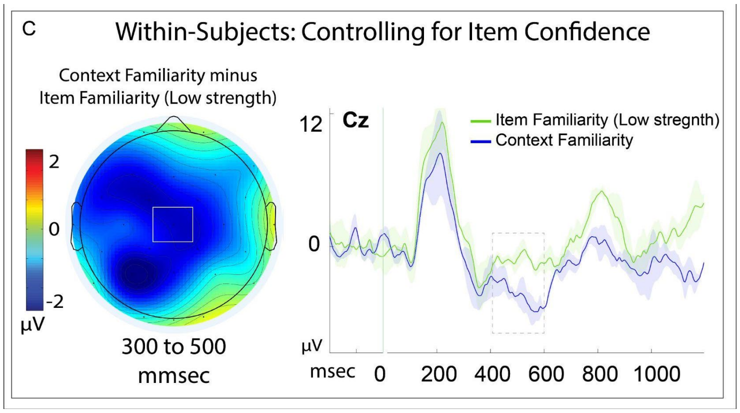


This finding was also replicated among a larger sample, too, derived from previously published data of participants performing the identical task (Addante et al., 2012a, 2012b) (N = 12): the same results were reproduced and found to be significant during the traditional FN400 time window of 300-500 ms). This revealed that there were indeed still comparable effects ($t(11) = 1.97$, $p = .037$, Cohen's $d = .569$), and thus the main finding from the primary study was replicated again.

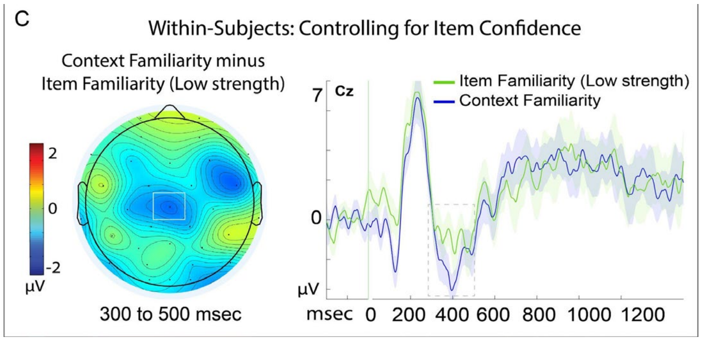


These findings have now been added to the revised manuscript in the Supplemental Materials Results section that describe the Experiment 2 replication of the primary ERP findings, which we believe substantively adds to the weight of the findings. Importantly, this revealed that neither the behavioral nor the physiological differences we discovered between conditions of item familiarity and context familiarity were confounded by inclusion of high confidence recognition trials in the item familiarity condition. We hope that the combined results here alleviate the concern about the item familiarity condition being potentially confounded by inclusion of high confidence recognition responses, and bolsters confidence in the robustness of the findings.  We are very grateful for the suggestion.

**R1.5. I understand the challenges with obtaining sufficient responses per condition in memory studies and appreciate the within and between subjects' analysis to address the limitation of there being only nine participants with sufficient trials for the within subject analysis. If the authors lower the threshold of minimum events (e.g. five per condition), how many more participants would be included in the item vs. context familiarity ERP analysis and are the results comparable to the currently reported findings?**

The Reviewer raises a very interesting point. However, it would be inappropriate of us to re-analyze the data based upon the suggestion to arbitrarily lower the inclusion criteria of trials per condition to a random number such as n = 5 in order to achieve statistical significance of any particular effect, as it would constitute the kind of p-hacking and harking that the journal's principles specifically guide against doing, and it also raises the possibility of including more noise instead of signal. The criteria we used for inclusion was provided in the manuscript (Methods) and was determined a priori based upon convention in the literature and for consistency with the same criteria used in the original study that the present investigation sought to replicate (and did). For that reason, unfortunately, we can not lower the threshold of minimum events needed for inclusion into the analyses.

However, we hope the Reviewer will be pleased to know that we have now updated a revised manuscript to include an analysis of an entirely independent data set of N = 56 subjects from previously published findings using the same paradigm in other laboratories, which replicated all of the ERP results reported in the present investigation and is now reported in our Supplemental Materials. The particular contrast mentioned by the Reviewer gained a sample of N = 20 meeting our a priori inclusion criteria for analysis (increased from the N = 11). We also performed a post-hoc power analysis that supported the findings as being detectable despite being fairly under-powered for detecting differences likely to be further detected in future studies having greater statistical power (*see details noted below in Response to Reviewer R3.c*). This addition of replication data provides a total sample of N = 110 people included in the manuscript's effort to gain insight into the possible differences between item and context familiarity, and we hope the Reviewer will find the convergence of evidence to be as compelling as we find it to be, and provide enhanced confidence in the findings.

**R1.6. The number of targets and lures are both inconsistently and I believe incorrectly reported. If there are 54 items per 4 study runs, that yields 216 targets which matches the methods (though not the caption of Table 1 which reports both 216 and 214 targets). However, if there are 54x6=324 test probes, there cannot be 116 lures. If in fact there were 108 lures, why were there half as many lures as targets rather than an equal number of targets and lures and how might this change participants' item and source judgments? To what extent are participants biased to respond "old" given the imbalance of targets and lures?**

We thank the Reviewer for pointing out the typos for numbers of trials that we had listed in the original manuscript. The numbers for trials have been updated in the revised manuscript (216 targets, 108 lures; 108 targets for each of the two source conditions of encoding task) and we apologize for that typo. In regard to the proportion of targets and lures, the proportion of targets to lures in the present study was designed to match the same proportion used in the preceding study that motivated the current work (Addante, Ranganath, & Yonelinas, 2012). The reason that the study used different numbers of trial types for old and new items in this experimental design is because this approach is nearly always used by studies to equate the number of each class of items (e.g., ~100 trials of old items from source1, ~100 trials of old items from source2 and ~100 trials of new items). This is standard for the field, and we do not suspect that it would account for the pattern of findings we observed.

We are not aware of any studies that would indicate that this type of proportion of targets and lures would alter accuracy. Rather, it has been found to increase a bias to say old to both old and new items (e.g., Rattcliff et al., 1992). Ian Dobbins and Mike Miller conducted many prior studies varying the proportion of lures to extreme extents, showing that it is extremely difficult to get subjects to change their criterion. The worst case scenario is that subjects may have a slightly higher response bias to say old, but that would likely only water down the observed effects, in that it would be watering down the pool of "old" trials. The present findings had no trouble detecting differences between old and new items, and the results for old and new items are very consistent with the other findings in the literature. Overall, we do not see a reason to assume any substantive impact of target/lure proportions on the pattern of results the present study focuses upon.

**R1.7.a: The authors report faster recollection compared to item familiarity responses during item judgments; however, this is counter to (some) of the findings in the literature (e.g. Rugg & Yonelinas, 2003; Boldini et al., 2004). Further discussion of this discrepancy is warranted.**

The Reviewer raises an excellent point about the patterns of response times observed in the present findings being somewhat counter to the traditional view of the respective timing of recollection and familiarity. However, a review of the literature also identifies well-known exceptions to these traditional patterns, which the present findings are well in alignment with. We have added a substantial discussion of these response time patterns to the revised manuscript, available in the Supplemental Materials: Discussion, as excerpted below:

> *"Traditionally, recollection has been conceptualized as a slower, deliberative process, whereas familiarity was modeled as a faster, automatic process (Yonelinas, 2002). Accordingly, item recognition tasks are usually considered to be a relatively better measure of familiarity-based responding, and faster response times observed here for item-familiarity than for context-familiarity in the present findings can thus be seen as it being the item (not the context) that is familiar during this early stage of the memory task.  In this view, that recollection responses were also faster in item recognition would be seen merely that their inclusion of the highest confidence responses was driving a concurrent higher amount of strong item familiarity and thus faster recognition time, consistent with traditional patterns. As the context familiarity responses then go on to become faster than item familiarity responses during the source judgments, it preserves the traditional notion that familiarity is also a faster automatic process- this time it is familiarity for context that is differentially faster (not items). That is, in a traditional view of RTs, our results are seen as showing that the two different types of familiarity (item, context) are faster than each other in each of the respective domains in which they are tested for representing differentially (item recognition, source memory), and ultimately consistent with the common findings that familiarity operates faster, while recollection is a slower process.*
>
> *Alternatively, the pattern of faster recollection during item recognition responses (and slower than context familiarity in the source judgments) could also be seen as departing from common findings that recollection is generally a slower process than. While some studies have found that recollection can sometimes be faster than familiarity in some instances (Gillund & Shiffrin, 1984; Trott et al., 1999), that discrepancy was suggested to likely be due to instructions to participants that they respond after both memory processes are complete- thereby delaying the responses times (Yonelinas, 2002).  The current paradigm also requires subjects to respond to a combination of both item and source*

*memory confidence ratings on each trial, which could similarly impose complex demands on their memory judgements. Moreover, recent studies noted below have reported recollection to have faster responses than familiarity during recognition, suggesting that the traditionally-assumed speeds of recollection and familiarity should not necessarily always be taken as simply one-directional (i.e.: Familiarity > Recollection) and that is likely more complex in directionality. For instance, a review of pupillometry studies of recognition memory concluded that, contrary to common observation of behavioral studies, recollection is not always the characteristic 'slow and effortful' process typically assumed, but can also be quite 'fast and easy' in its manifestation as a memory process (Goldinger & Papesh, 2013). When more advanced modeling methods were used for measuring the speeds of memory processes across six different experiments, recollection was again found to operate as both fast and slow (Brainerd et al., 2019), similar to findings in the present data. Such patterns are also consistent with proposals that recollection is a two-stage process that includes both an initial rapid assessment of memory, and a later second stage that is slower and more deliberative (Moscovitch, 2008). Together, the response time profiles observed in the present work are consistent with other studies, while also adding insight from specific memory response conditions that have not previously measured response times, to reveal that regardless of chosen interpretation views, the conditions remain nevertheless as neurocognitively distinct."*

**R1.7.b: More importantly, the authors state, "if our LCN was actually an LPN representing a controlled search process, etc., then it would have also presumably been seen in the recollection condition." However, given the fast reaction times for the recollection condition during item judgments, it is unclear whether a "controlled search process" should be observed for this condition. Do ERPs dissociate across slow vs. fast recollection responses**?

Regarding the 'controlled search process' ascribed to the LPN, it was only mentioned in regards to recollection because the LPN has been found in other studies to be associated with correct source memory retrieval (e.g. recollection). Regarding the Reviewer's question about whether ERPs dissociate across 'slow' and 'fast' recollection: we are not aware of any findings that do so. It may be important to note that ERP effects serving as the putative neural correlates of memory processes such as recollection (Rugg & Curran, 2008) are merely correlates of memory conditions, and represent covert neural activity upon initial processing a memory probe from stimulus onset (usually occurring prior to 1 second) (Luck, 2014), whereas memory judgments tend to occur slightly (if not much) after that in time, strongly suggesting that there need not be a direct 1:1 mapping of timing across behavioral and electrophysiological measures of initial, internal memory processing, and its later manifestation of outward memory judgments, as was noted in the TICS Review by Rugg & Curran (2008); *see also the responses to Reviewer 2.4 below*.

**R1.8. Some of the reported ANOVAs are a bit confusing and the interpretations are unclear. e.g. "A 3x2 two-factor ANOVA using factors of condition (i5s4, i4s4, and i45su) and time for response type (item RT, source RT) with repeated measures" and "a 2x2 two-factor ANOVA of response speeds among item and source judgments for recollection and context familiarity." Reaction time seems to be both a dependent and independent variable in these ANOVAs (e.g. "a condition by response-time interaction") which is puzzling since it seems that reaction time should only ever be a dependent variable in these analyses. The authors state "These findings indicated that, overall, the RTs for item and source judgments were the same, yet still differed across the memory conditions in different ways." This sentence is very vague and would benefit from explicit statement of the direction of the effects. The motivation for conducting these analyses could be made more explicit, as could how the results should be interpreted**.

The Reviewer makes an excellent point here, and we are grateful for pointing it out. The prior version of the manuscript was insufficiently clear about the elements of the behavioral analyses, and the revised manuscript has now been updated to be more clear and concise on this front, in addition to being more computationally efficient. For instance, the reviewer is correct that response time is a dependent measure, which we now note in the manuscript. We have also removed the poorly-worded parts noted by the Reviewer, because upon reflection we realized they were unnecessary analyses, and we added a better discussion of interpretation of the behavioral findings via an extensive review of the literature in the Supplemental Materials Discussion and *excerpted above in our Response to 1.7a*.

**R1.9. I realize that in prior work the authors found FN400 effects around Cz (though it does not appear that this selection was supported by appropriate direct statistical tests); however, convention holds that the FN400 is (by definition) frontally located. Are the results the same if a different electrode (Fz) or group of electrodes (e.g. F3/F4) are used instead of Cz**?

We thank the Reviewer for raising these important points. As correctly noted by the Reviewer, the present study's purpose was to assess replication of a prior finding (2012), and so we performed targeted analyses on the representative site (Cz) where those findings were reported, which we believe is a reasonable and sound approach to seeking replication. For whatever it may be worth since it was also raised by the Reviewer, the 2012 paper clearly described the process of electrode site selection (Methods: EEG acquisition and analysis, paragraph 4), where it notes that the FN400 was first identified at frontal site Fz via a priori hypotheses, but also observed to be maximal at its adjacent site of Cz, where appropriate statistical tests were employed to further characterize it. One benefit of the present study's aim to replicate the prior study's findings is that if any statistical tests in the first study may have been inappropriate (we do not think they were), then it presents the hypothesis that the results would be unlikely to replicate if the study was performed again (but they were replicated, as evidenced in our present findings).

Regarding the statement that "*convention holds that the FN400 is frontally located*", a growing body of work has more recently identified the topographic variability of the FN400 to also include posterior sites, beyond its earlier convention of having been assumed to be exclusively frontally-based. For instance, there are fantastic reviews of the literature done by Mecklinger & Bader (2020) identifying how the FN400 can be influenced by forms of either relative or absolute familiarity to manifest in either a frontal or posterior scalp distribution (see also Wolk et al., 2004; Bridger et al., 2012; 2014; Leynes et al., 2017).

Regarding the Reviewer's question about the extent to which effects may hold at other frontal sites: the topographic maps provided for each figure provide a visual depiction of the extent to which each of the respective effects hold for more frontal regions. To preserve the spirit of replication, we didn't expand the analyses of the FN400 to the other frontal regions, as even if they were significant, the data is clearly mid-central focused for this particular effect of item familiarity. We have made concerted efforts to update the revised manuscript to included discussion of the topography of this effect, as noted below excerpted from our Supplemental Discussion:

> *"The posterior distribution of the FN400 effect for the item familiarity condition is consistent with other findings ascribing it to a sub-type of absolute familiarity (as opposed to relative familiarity, which is reported to have a more anterior scalp distribution; (Bader & Mecklinger, 2017; Bridger et al., 2012; Mecklinger & Bader, 2020; Stróżak et al., 2020). The FN400 effect observed in the present study had the same mid-central topography as was reported originally (Addante et al., 2012), which is between the frontal and posterior distributions observed in other studies that manipulated factors of relative and absolute familiarity (Andrew Leynes & Upadhyay, 2022; Bader & Mecklinger, 2017; Bridger et al., 2014; Leynes & Mok, 2020; Leynes et al., 2017; Mecklinger & Bader, 2020; Stróżak et al., 2020), respectively, which likely reflects a combination of both relative and absolute familiarity with the stimuli in this particular paradigm since it was not designed to assess those factors. That the ERPs for conditions of item and context familiarity were found to differ in the N400 potentials from 300-500 msec, and the N400 component has also been known to be influenced by factors such as context, is consistent with the present interpretations of the psychological constructs represented by the condition we describe as 'context familiarity*'. "

**R1.10. It is unclear why a failure of context BINDING is relevant for a participants' memory for context ("During that time (600-800 msec), that process of seeking episodic context to bind with the presented item fails") or how if (retrieval of) context fails during the item judgment it can then be successfully retrieved during the source judgment**.

Thank you for raising this good point. The purpose and premise of the statements quoted above stem from the BIC model as noted in the manuscript (Eichenbaum et al., 2007; Diana et al., 2008), which proposes that recollection occurs when the episodic context is bound together with the item representation; thus, if that binding of item and context fails, then recollection would presumably not occur. In that example, one could retrieve the context of a prior episode not successfully bound together with its item, similar to as has been described by Whittlesea & Williams (2000).

To the Reviewer's second question, we would suggest that if context was the element of the memory that was familiar, that its retrieval would guide a participant to necessarily report that the item had to be 'old' (e.g. familiar; not because they retrieved item information but because a familiar context would logically suggest a prior occurrence of the item) and this retrieval of context would guide the correct source judgment. We have now taken steps to update the revised manuscript so that these aspects are more clear. We have also added an expanded the Discussion section to include better clarity and elaboration upon those points, particularly expanding upon the prior work of Whittlesea & Williams (2000) as excerpted below:

> "*The idea that there can be different kinds of familiarity for items and for contexts has been previously mentioned by several others (Addante et al., 2012; Montaldi & Mayes, 2010; Whittlesea & Williams, 2000). Whereas prior work established the premise that there can be separable familiarity of items and familiarity of contexts (Montaldi & Mayes, 2010; Whittlesea & Williams, 2000), they nevertheless proposed that each was stemming from the same singular memory process of underlying familiarity. For instance, the Discrepancy Attribution Hypothesis, offered that people can produce either piece of mnemonic information separately (Whittlesea & Williams, 2000), stating: "contextual information can also cause the person to experience a feeling of familiarity: encountering a stimulus in a context that is unusual for that stimulus (e.g., a clerk on the bus) can cause this." However, and importantly, they go on to propose that "the use of contextual information to perform judgments about the past is not qualitatively different from the use of identity" (page 561) (Whittlesea & Williams, 2000). Here, we put forth a fundamentally different view: that production of familiar context does differ qualitatively from the production of familiar item information, and provide converging lines of support that the familiarity of items and familiarity of contexts are evidently operating as separable independent neurocognitive processes*".

Minor.

**R1.a: Figure 1 is overall very clear, but I would recommend swapping the positions of the two judgments in panel A since most readers will read top to bottom and it seems like the source judgment comes first (despite the arrow)**.

The revised manuscript has now been updated accordingly.

**R1.b:. "i5s4, i4s4, and i45su" are not used elsewhere in the text, I recommend replacing with the condition names recollection, item familiarity, context familiarity**.

The revised manuscript has now been updated accordingly.

**R1.c:. When describing reaction times, it is easier for the reader if the 'direction' is consistent, e.g. instead of "recollection responses were faster than responses for both context- and item-familiarity" and "Context familiarity responses for item judgments were also reliably slower than those for item familiarity" both sentences should be written in reference to either faster or slower responses**.

The revised manuscript has now been updated accordingly.

**R1.d: The "four temporal latency windows" should be explicitly referenced in the text**.

The revised manuscript has now been updated accordingly.

**R1.e: How/was semantic relatedness between words controlled**?

We thank the Reviewer for pointing out that the earlier version of the manuscript had omitted details about the stimuli, which was our oversight. We have now updated the revised manuscript to note the details of the stimuli, which includes that the word stimuli were selected from the same source as stimuli used in the original study in 2012, and preserved the same characteristics. Semantic relatedness was not a measure that had been logged for the stimuli, but it is unclear to us how that would have been a factor for which the present results would be attributable to. Word stimuli were selected from the Medical Research Council Psycholinguistics Database (http://www.psych.rl.ac.uk/MRC Psych Db.html). Word stimuli had an average rating of concreteness of 589 (min=400, max=670), image-ability of 580 (min=424, max=667), Kucera–Francis Frequency of 30 (min=3, max=198), and an average number of 4.9 letters in each word (min=3, max=8).

**R1.f:This statement in the discussion sounds circular "This rules strongly against the alternative possibility that somehow context familiarity effects were being driven in some undetectable way by the other standard processes of recollection and/or item-familiarity, and leaves open the core question of what, then, the effects of the condition represent. We contend that it represents the evidently independent memory process of context familiarity." It reads as "This rules strongly against the alternative possibility that somehow context familiarity effects were being driven [by XYZ and instead] it represents the evidently independent memory process of context familiarity**."

Thank you for pointing this out. We agree that this statement didn't make very good sense, so we have now removed it from the revised manuscript.

**R1.g: The discussion subsection heading "Are effects representing implicit memory driven by fluency or guessing?" seems incorrect -- shouldn't it be something like "Are effects representing context familiarity driven by fluency or guessing?**"

The revised manuscript has now been updated accordingly.

**Reviewer #2 (Remarks to the Author):**

**R2: Addante et al present behavioral and electrophysiological (scalp EEG) data addressing the component processes of recognition in episodic memory. Working within the recollection/familiarity process framework, the authors report evidence for a third process they term 'context familiarity.' Behaviorally, response times show that context familiarity is statistically different from recollection and familiarity. Electrophysiologically, the authors report ERP signals that are significantly different across time and space (electrode) for item familiarity, context familiarity, and recollection trials. This study contributes to a longstanding debate about the processes that contribute to recognition in episodic memory, and therefore seeks to address an important question. I have some comments about the interpretation of the context familiarity process, its neural basis, and the degree to which it can be differentiated from other previously studied memory processes**.

> We thank the Reviewer for taking the time to provide a thoughtful, insightful, and helpful review, as well as for both supportive and constructive comments that we believe have led to valuable improvements to the manuscript.

**Major Comments**

**R2.1: The timing of context familiarity: typically, familiarity is thought to be a faster process than recollection, meaning that individuals can make use of the familiarity signal earlier than they can a recollection signal (Eichenbaum et al., 2007). Along with behavioral data, there is prior neural evidence for faster familiarity compared to recollection in the medial temporal lobe. In the ERP data here, the context familiarity signal occurs much later than the 'standard' item familiarity signal (800-1200 ms vs. 400-600 ms). Why, under the authors' model, should context familiarity occur so much later than item familiarity? I am specifically wondering about this in regard to the neural basis of context familiarity: in the Discussion the authors propose that parahippocampal cortex could subserve context familiarity processes (while perirhinal cortex supports item familiarity). Is there something about this neuroanatomical distinction that would predict such timing differences**?

> The Reviewer raises a very good point about the relative timing of the effects. First, regarding the Reviewer's question about the neuroanatomical basis for context familiarity (and its timing) in the parahippocampal cortex: the basis for our ascribing it to the parahippocampal cortex stems directly from that being the role already ascribed to it in existing models of the medial temporal lobe, particularly as described in the Binding in Context (BIC) model (Eichenbaum et al., 2007; Diana et al., 2008). In that regard, we are only echoing what we and others had previously reported as neuroanatomical substrates for item familiarity and context familiarity processing (Montaldi & Mayes, 2010; Addante,

Ranganath, & Yonelinas, 2012; Addante et al., 2012b), and this was consistent with similar fMRI dissociations that have been reported, too (Diana, Yonelinas, and Rangnath, 2013, 2012). Importantly, those regions are quite unlikely to be direct volume conducting to the cortex and then scalp, so it is very unlikely that the subcortical anatomical distinction in the MTL mentioned by the Reviewer could, in itself, support making predictions about timing of memory effects at the scalp (though we note that some intracranial recordings have shown temporal dissociations of basic familiarity and recollection amid the rhinal cortex and hippocampus, respectively (Fernandez et al., 1999, Science), and we are in the process of conducting more specified iEEG studies that are better suited for testing the hypothesis that item- and context-familiarity temporally dissociate in the MTL subregions.

Second, regarding the Reviewer's second question about why, under our proposed model, context familiarity might occur so much later than item familiarity. In our main ERP contrasts of each memory condition, context familiarity manifested effects later than item familiarity(800-1200 msec, 400-600 msec), and behavioral response times for context familiarity judgments were similarly later than those for item familiarity (2710 msec, 2361 msec). [Context familiarity responses were faster, though, than item familiarity during the source memory judgments (2075 msec, 2612 msec)]. These timings of ERP effects are occurring (for all conditions, in generally all studies throughout the literature), substantially before the later behavioral memory judgments, and thus it is quite important to note that the ERP effects have never been thought to represent a direct 1:1 mapping of memory process (as discussed in Addante et al., 2012b) but instead are widely regarded as "the putative neural correlates" of the memory processes as stated and Discussed by Rugg & Curran (2008, TICS Review). That is, the ERPs are *covert* measures of memory-related activity (Luck, 2016), but the *overt* memory is generally thought to occur later in time (when people indicate their memory response). So, there should be caution exerted in assuming too much about the timing of ERPs- what matters most, in our view here, is that they differ at all.

Third, regarding the request to speculate about *why* this timing difference could be? To a certain extent, the present work is not designed to directly answer *why* the times differ, but importantly is reporting the original and novel finding that *they are, in fact, different at all*. One speculative reason is that context information may be more broad than item information: encompassing additional elements/features that require neural pattern completion- and that this could take the extra few hundred milliseconds that we observe here. It will likely take future studies that we hope are inspired from the present findings in order to adjudicate *why* it is happening, but we look forward to that.

Finally, we note again here that the later ERP time interval (800-1200 msec) was not the only time that context familiarity was found to exert an effect: it also differed reliably from item familiarity ERPs during the early epoch of 300-500 msec (Figure 5), too. We underscore that this early finding was quite important to the present work, because it demonstrated that at a fundamental level the two memory conditions are occurring through differential patterns of neural activity very early in the epoch (300-500 msec)- not just later in the epoch.

**R2.2: In the Discussion, the way the authors describe the cognitive basis for context familiarity sounds a lot like a controlled memory search process, which they acknowledge and contribute several paragraphs to rebutting. However, I do not think that this argument is strong enough, and I think the authors need to do more to support it, if they want to claim to have identified a separate context familiarity process. Specifically, there needs to be additional analysis of the data (not just comparison of the current effects to those found in previous studies) to suggest that the context familiarity trials are not simply those where participants are using controlled memory search**.

This is a very interesting point to consider, and we thank the Reviewer for helping us think in this way. It is a substantive question, which overlaps with several other points raised by this and other Reviewers, so the response to it is unpacked here in several sequential lines of reasoning from the range of evidence that has accrued. We believe the Reviewers raised very important points about this, and we have since revised the manuscript to incorporate these points in a lengthy, more substantive discussion in the Supplemental Materials.

First, in regard to the request for additional analyses of the data: we have now updated the manuscript to include an entire independent replication of the context familiarity differences found between item familiarity, drawing upon previously published datasets utilizing the same paradigm. Indeed, these familiarity-based differences between item and context were apparently there in the original 2012 data but the first author (RJA) had, myopically, not thought to look for it (that 2012 study was, instead, premised upon investigating the relationship of this memory response condition to recollection). Evidently, all we had to do was look for it, and we found it, again. These new results are now reported as Experiment 2 in the Supplemental Materials, and it reproduces all of the ERP effects we report in the present findings. Furthermore, we have conducted additional analyses as described in response to *Reviewer 1 [see point R1.*4], which controlled for important potential confounds and still revealed core differences in the memory processes. We hope that this replication of the novel findings here inspires more confidence in the validity of the effects.

Second, in regard to the question about if the trials for context familiarity are ‘simply’ reflecting instances in which participants used a ‘controlled memory search’, we think this requires some thoughtful consideration to unpack. To begin with, in theory, every trial of the study should be inducing participants to engage a ‘controlled memory search’ simply by virtue of the memory question being asked of them. That is, the participants are expected to be searching their memory on every trial for if they remember the item and the source of its encoding condition, then reporting it. So in that sense, the controlled memory search would be a variable that is held constant across all conditions, thus canceling out in our contrasts, and leaving open the question of why this particular memory condition would show the effect when compared to other conditions that also involve the searching of memory do not show the effect.

For instance, in the item familiarity condition (i.e. successful recognition memory hits), participants report that they do not have source memory for the encoding task, and spend longer response times to say so on the source memory task (even when removing the high confidence recognition trials from the analysis- which is a new analyses suggested by Reviewer 1.4, see above). This provides a solid premise to infer that the condition would also include a "controlled search of memory" for the source information that still comes up empty for them, and makes a clear hypothesis that we should see a major negative-going ERP effect on these trials if it reflected the same thing as the LPN (Mecklinger et al., 2016). Yet, that is not what the data shows here, nor in several of the independent replications of the finding across different measures and laboratories, and instead we see that the item familiarity condition differs reliably from the context familiarity condition. Hence, the differences must be attributable to another variable than just a controlled search of memory.

As another example of a line of evidence arguing against a 'controlled memory search' account of the LCN correlates of context familiarity effects, we also refer the readers to the previous findings (Figure 7 in Addante, Rangananth, and Yonelinas, 2012) when the LCN effects for context familiarity were also evident when compared to incorrect source responses that were matched on levels of item & source recognition, which again presents a situation where both conditions should be reflecting a controlled search of memory- yet in which they differ reliably based upon another factor (e.g. context familiarity).

Third, as a final note on this point, a 'controlled memory search' noted by the Reviewer is one of the primary processes that the Late Parietal Negativity (LPN) ERP effect has been attributed towards (Mecklinger et al., 2017, *Neuroscience and Biobehavioral Reviews*; Mecklinger & Johannson, 2006), which the Reviewer here acknowledged we addressed in our Discussion, but asked for more on. A similar point was made by Reviewer 3, wondering if the effects we report here might be reflecting an LPN effect instead of our interpretation. So, here we turn directly to what the authors of the LPN Review paper(s) specifically noted in their most recent, authoritative Review paper on the LPN, where they noted in several places that the LPN was not the same effect as what we report(ed) here (2012) as the LCN:

> (page 631) *"Even though these late negativities have been associated with attempts to reconstruct memory-stored information, processes usually ascribed to the LPN, the non-posterior scalp distributions suggest that it is unlikely the same ERP component as the LPN. The same objection holds for the late negativities reported for young adults in the studies by* ***Addante et al. (2012a),*** *Leynes and Zish (2012), Leynes et al. (2013) and, for amnesia patients, by Addante et al. (2012b). All of these studies report negativities that are larger for*

*studied than unstudied information but due to a lacking posterior scalp distribution* ***these components are unlikely the same ERP component as the LPN****." [emphasis added].*

And (page 634): *"****Addante et al. (2012a)*** *examined ERP correlates of source memory for items that were recognized with high or low confidence. A parietal old/new effect (the ERP correlate of recollection) was obtained for correct source judgments, but only for items recognized with high confidence. Correct source judgments for items that were less confidently recognized* ***elicited a late but broadly distributed negativity, resembling the LPN****. It could be argued that the low confident item memory in this study constitutes a paradigmatic case for a high need for continued evaluation of source-specifying memory information and LPN generation. However, it could also be argued that the low confidence for the item judgments is a direct consequence of the failure to retrieve source specifying information. Irrespective of this interpretational ambiguity,* ***the rather broad scalp distribution and the early onset of the negativity do, however, not necessarily support an LPN interpretation of this effect****" [emphasis added].*

These statements by the LPN authors converge with what was originally stated by Addante, Ranganath, & Yonelinas (2012) about the potential interpretations of the original LCN findings as representing the 'controlled search process' of the LPN:

**"***There are several key distinctions though, which suggest that the ERPs found in the current study are not likely to be reflecting the same type of processing as the LPN. For example, while the negative ERP observed in the current study emerged at anterior sites from 600 to 800 ms, its topography was generally much more widespread, becoming insensitive to anterior/posterior factors from 800 to 1000 ms. Additionally, the LPN is not generally observed in studies such as ours that utilize item judgments followed by source discriminations (Duarte et al., 2004; Trott et al., 1999; Wilding, 2000). Finally, when the LPN has been observed in source memory tasks, it has been found to be either insensitive to source accuracy (Herron, 2007) or to be even larger for inaccurate source judgments (Johansson and Mecklinger, 2003; Wilding, 1999)."*

The present study's findings provided a complete replication of the prior findings that were discussed in the several quotes above, and as such also provided a vindication of the interpretations of the LCN effect not meeting the broad criteria of representing an LPN effect. In fact, we believe that the present findings make it more possible that many of the reported LPN Effects could have instead represented findings that could actually reflect context familiarity (e.g. Strunk et al., 2016, *Neurpsychologia*). We believe that this has been a very important and fruitful discussion about the nature of effects and their potential interpretations, and have thus updated the Supplemental Discussion section to reflect this additional information to better augment the manuscript. We believe we have provided a thorough and exhaustive effort at analyses ruling out such alternatives, but also that these are the kind of future research questions that we hope the present work can inspire to be conducted moving forward.

**R2.3: Related to the previous point, if participants are in fact using controlled memory search on the context familiarity trials, you would expect them to show the pattern of response times that is seen here. For example, participants may find on context familiarity trials that they are unsure about the familiarity of the item. They then engage controlled search using the two sources that were used in the experiment to probe the representation of the item in memory. The time that this process takes would then lead to slower response times to the item familiarity probe–but then, because participants have already probed the source and used it to evaluate the old/new status of the item, they then respond more quickly to the source probe. Can this hypothesis be ruled out?**

The Reviewer makes some excellent points here regarding response times. We first note that in response to Reviewer 1, we have now added an additional finding of response time effects of context familiarity as compared to item familiarity conditions that excluded the high confidence recognition response (*see response to comment R1.4 above*). This contrast found that context familiarity trials were now slower than item familiarity during recognition, and argues against the prediction made in the Reviewer's comment here that a controlled search process would be evidenced by context familiarity being faster during item recognition. In that respect, we can rule out the hypothesis put forward by the suggestion in the comment above via having now directly tested it, and incorporating those findings into the revised manuscript.

Furthermore, see also the same points mentioned above *[in R2.2*] about the nature of a 'controlled search process' being presumed to be occurring on every trial, across all conditions, because the memory test is asking participants to search their memory each time, for over 300 trials. Thus, it's difficult to accept the premise that only in certain times would they "*then engage a controlled search*…". In the present paradigm, participants know that both item and source will be probed sequentially on each trial, so while the Reviewer's proposed account here is logical, upon further consideration it is also likely that the account's proposal that participants *"already probed source and used it to evaluate old/new status of the item*" and "*engage a controlled search process using two sources used in the experiment*" could also be occurring across a range of different trials, whereby our task can only probe item and source sequentially to capture the various different combinations of memory responses we report.

**R2.4: My understanding of the Methods is that the ERP data were analyzed with respect to the onset of the item probe. What about the ERP effects that occur following the onset of the source memory probe? This interval seems like it would be critical for differentiating context familiarity as independent from item familiarity or recollection. All the reported ERP effects occur no later than 1200 ms after stimulus onset, yet the response time averages for recollection/item/context are all greater than 1800 ms. Following the item memory probe, participants then spent 2000 ms on average or more**

**on the context/recollection/item source judgments. So, there is a lot of time spent on these second decisions–is there any evidence that context familiarity is distinct from item familiarity and recollection during this interval**?

Reviewer is correct that ERPs were analyzed with respect to the onset of the item probe. We did not analyze ERP effects that might occur after the onset of the source probe, for several reasons. First, given that it followed differential item recognition motor responses, the baseline period for the ensuring source judgments could become problematic for ERP analyses, and further confounded by inclusion of source memory responses [*see similar response R1.1 to Reviewer 1*]. Second, the purpose of the study was to assess the replicability of the previously reported findings (2012), and thus required consistency to the same analytic approaches- which in this case meant assessing ERPs to the onset of the recognition probe, since that is what the prior study did. Third, interpretations of the ERP effects to the broader literature (especially the much-better known FN400 and LPC) requires comparable analytic parameters- which for the ERP effects of memory involve being time-locked to the onset of the memory probe (not response-locked to a secondary memory probe) [*also noted in R1.1 above*].

In that respect, it is still the time interval of activity following the onset of the memory probe that remains critical for understanding the respective memory processes/activity involved in what goes on to be remembered (and how it does so). The field of memory ERPs has always looked at the FN400 and LPC as the putative neural correlates of the memory processes, and not a direct 1:1 mapping of the actual processes nor their occurrences, as Rugg & Curran (2008) stated in their Review paper (see also discussion Addante, Olichney, et al., 2012; Muller, for similar points). In doing so, the field has understood memory operations for recollection and familiarity by virtue of the underlying physiology associated with the brain's response to a stimulus that leads to differential patterns of memory responses (i.e.: Rugg et al., 1998). We preserve that core foundation in the present investigation because for our goal of investigating the possibility of context familiarity that was raised in the 2012 findings it requires it to be directly contrasted to item familiarity effects as they are traditionally known: time-locked to the onset of a stimulus and not response-locked.

Regarding the Reviewer's point about the response times for the ERPs occurring prior to the response times for the behavioral judgments (see also the *Responses to Reviewers 1.7, and 2.1, for related response information*): first, yes, there is evidence that context familiarity is distinct from item familiarity and recollection during the time interval of the second decision (source memory), in that they exhibit reliably different response times for the second memory judgment. Second, the Reviewer is correct in noting that the timing of the ERP effects all precede the timing of the behavioral responses, but we would merely offer that this is not out of the ordinary, as we noted above that ERPs reflect the covert neural activity that typically precede the overt behavioral responses of memory (Luck, 2014). We agree that it would be interesting for future studies to look at potential ERP activities of memory response later in such epochs, but we leave it to future studies, since we first needed to establish here if/that the different kinds of memory response patterns differ at all (it appears from our converging data here that they do). We hope the current work can inspire those kinds of follow-up investigations.

**Reviewer #3 (Remarks to the Author):**

**R3.0: Addante and colleagues report a new set of analysis of an already published study. The aim of this report was to dissociate the ERP correlates of three dissociable processes in recognition memory. In addition to (item) familiarity and recollection, the authors claim to have identified a third process: context familiarity. The paradigm comprises a source memory task (where source denotes one of two study tasks) in which participants gave confidence judgments (1-5) within the item memory test and within the source test. The authors defined three contrasts of interest: item familiarity (item hit (conf 4 + 5) + source miss); recollection (item hit (conf 5) + source hit); context familiarity (item hit (conf 4) + source hit). With the first two contrasts, they replicated earlier work and found the FN400 effect for item familiarity and the LPC effect for recollection. For context familiarity, they identified a dissociable ERP old/new effect, the LCN effect, between 800 and 1200 msec post-stimulus. I have great sympathy for the research question and think it is theoretically well motivated. However, I have doubts that the current design is suited to isolate context familiarity**.

We thank the Reviewer for taking the time to provide a thoughtful, challenging, and helpful review, as well as for both the supportive and constructive comments that we believe have led to valuable improvements to the manuscript. We have carefully considered the points raised, and they led to very good additions that we have now incorporated into the updated manuscript.

**R3.1: In the beginning of the introduction, I could easily follow the examples given for context familiarity, e.g., a familiar place for which I cannot remember the person I met there. These examples are similar to those usually given for item familiarity (a familiar person for whom I do not remember the place I met her). From this description, it is clear that context familiarity (in the same way as item familiarity) is best identified when the context is familiar but there is NO recollection of the association to the item. Therefore, a context familiarity contrast should include those trials in which a context was familiar but not the item and there was no memory for the association between them. To be clear, I do not think that context familiarity can only occur when there is no familiarity/recollection for the item. However, the absence of memory for the item and especially the association between item and context is a prerequisite to unambiguously identify a neural correlate of context familiarity (as it is usually defined the other way round for item familiarity). In the current paradigm with only two different contexts, context familiarity cannot be isolated trial-wise**.

This is a very good insight and terrific comment from the Reviewer; we will try to unpack it in a few segments noted below.

First, there may be different routes by which to identify an admittedly-difficult construct such as context familiarity, and while the Reviewer has noted one excellent approach (i.e.: "*The absence of memory for the item and especially the association between item and context is a prerequisite to unambiguously identify a neural correlate of context familiarity*), we believe that there may be others, too. The route by which we had arrived at the construct of context familiarity came via a different way. The original observation in 2012 was that we were able to identify instances when people provided accurate source judgments for items recognized with low confidence, which led to assessing the ERPs for such rare instances. We thought it would be associated with an LPC if supported by recollection and/or an FN400 if supported by familiarity; we were surprised when it exhibited neither effect but revealed a wholly new one. As noted in that paper's excerpt below, we developed the idea of context familiarity as a way to understand the observed effects within the framework of existing models (BIC; Eichenbaum, et al., 2007; Diana et al., 2008)):

> "*A second, more speculative account of the late negativity that was related to source memory for low confidence recognition is that it reflects a form of `contextual familiarity'. The background for this idea is that episodic memories may reflect the binding of neural representations of item and context information (Diana, et al., 2007; Eichenbaum, et al., 2007). According to this view, item familiarity may reflect the strength of the item representation (e.g., a word) that supports recognition memory. Context information might include information about the place and time it was encountered, and how it was processed (see Ranganath, 2010 for review). Recollection can be thought of as the recall of context information associated with an item, but it is possible that processing of a studied item could elicit weak activation of the associated context representation even when recollection fails. This type of contextual familiarity signal, in turn, could support source discrimination*"

From that original interpretation, it was the condition of 'low confidence recognition with correct source memory' that became the potential vector for measuring the construct of context familiarity in the present study. The name of 'context familiarity' stemmed logically from the theories made from various memory models (Eichenbaum et al., 2007; Montaldi & Mayes, 2010; Whittlesea & Williams, 2000), but like Shakespeare famously said: "a rose by any other name would smell as sweet"- that is, the name matters less than having identified an important independent construct of memory processing/es.

Second, some points regarding the comment about the need to "*unambiguously dissociate memory processes*". In this study, we sought to assess if the neural activity/effects for this condition could be unambiguously dissociated not only from recollection (to validate the prior findings via replication) but also from item-familiarity (to test predictions made by the 2012 paper, and extend the prior findings in new ways). ***We believe strongly from the present data that it has done just that, and did***

***unambiguously isolate context familiarity in a trial-wise way*** (and across subjects, too; Figure 5B). Indeed, our approach of the present study to isolating differential memory processes was the same way that the dissociable neural correlates for recollection and familiarity responses have been widely taken as a gold standard for their unambiguous dissociability as dual memory processes (Rugg et al., 1998, *Nature*; Rugg & Curran, 2008, *TICS*; Kwon et al., 2023) even though recollection-related responses are widely understood to also be non-exclusive, non process-pure, and contain inherent familiarity (with recollection responses tending to include both FN400 and LPC effects of each process; e.g. Ozubko et al., 2021). As detailed in the manuscript, the ERP activity we reported here for the condition we identify as context familiarity was unambiguously dissociated from that for item familiarity by virtue of Bayes Factor evidence for an absence of well-established activity of item familiarity processing (FN400) and the evidence for an absence of 'association between item and context' (recollection, LPC) as suggested as prerequisite by the Reviewer.

We very much agree with the Reviewer's sentiment that "*context familiarity can [not] only occur when there is no familiarity/recollection for the item*", but we think that there are other effective ways at demonstrating a dissociation than just those in which there is "*an absence of the memory for item information*". Indeed, our particular condition studied in the present investigation may likely be representing that kind of instance of when '*there is no familiarity for the item nor recollection*', in that their item responses could be driven by the familiar context that de facto leads the participant to deduce that the item must have likely been 'old' if there is something else [i.e. context] about it that seems familiar, and we contend that the physiological way that we have dissociated the differential memory response patterns can be an effective way to identify memory processes consistent with the literature, as opposed to starting with memory processes and seeking if they demonstrate differential activity.

Among that good foundation of reasons to have confidence in the findings is the fact that there is an *unambiguous* 3rd way that people are making correct, explicit episodic memory judgments, beyond traditional processes of recollection and item familiarity. We see it repeatedly, over several studies, laboratories, contrasts, and measures. We draw upon the prior literature in 2012 to call it context familiarity, and here we show that its neural correlates are dissociable from those of item familiarity (and recollection, and behaviorally, too). This pattern can not be accounted for by predictions made by single or dual process models of recognition without making post-hoc modifications. These findings are reproducible and robust. A dissenting view is welcome to call it something other than context familiarity, but it is burdened then with providing a viable account that can handle the array of different converging observations thus far. Any alternative account to the present findings' interpretations must be able to accommodate these lines of evidence in a parsimonious way. We respectfully believe that an account such as the LPN (*as discussed in Responses to Reviewers 2.2 above and below*), of this effect simply being "searching memory" is insufficient to account for the totality of the findings, as is an account of it being merely 'reconstructing the past'.

**R3.2: For the context familiarity contrast specified by the authors (hit 4 + source hit), it seems equally plausible that it reflects a delayed recollection (or another search) process instead of context familiarity**.

> The reviewer raises a good and interesting point. We have thought about this possibility a lot since originally reporting these findings in 2012, and believe that there are good reasons why it is likely that the context familiarity condition could not plausibly be reflecting a “Delayed recollection” or “another search process”. For instance, there is little evidence to support the effect reflecting a contribution of recollection: there has been absolutely no ERP Evidence for it across several different studies, nor behavioral indications of recollection, and this is further supported by Bayes Factor evidence of absence. Alternatively, the evidence that has accrued thus far across various studies, measures, and contrasts for context familiarity continues to unambiguously indicate that it differs reliably from recollection trials in both behavioral response times for item recognition, and for source memory judgements, and then also differs reliably and reproducibly in several ERP measures across several experiments.
>
> The context familiarity condition of ‘correct source judgments for low confidence recognition items’ also tends to be low confidence source judgments, which also argues against the premise that even a later process of slow recollection might be driving their trials following a delayed memory search, because if recollection occurred later it could be assumed to have more high confidence correct source judgments (and yet the data doesn’t have that profile). Furthermore, if the effects of the context familiarity condition were reflecting “another search process” then one would expect that the ERPs would not differ from the item familiarity condition in which participants are also searching their memory for source information (i.e. source unknown) and yet respond more slowly to for those judgments- yet those ERPs do differ reliably, and reproducibly (*for more extended detail on similar point see response to Reviewer 2.2 above*). The ‘delayed recollection/search’ account also would have trouble explaining why the context familiarity ERPs differed reliably from the item familiarity ERPs during the early latency of 300-500 msec if it was only representing a delayed process happening later. For these reasons, we do not think that there is evidence available that the effects of context familiarity could plausibly be reflecting a delayed recollection or search process.

**R3.3: I have also some concerns regarding the analysis: If I understand it correctly, source memory performance is calculated by dividing the total number of source correct trials by the total number of all correct and incorrect source memory responses. However, source unknown responses (3) are excluded from the denominator, which in my view is not correct. For the extreme case that a participant chooses a source only in two trials and does so correctly but indicates in all other trials to not know the source. This participant would have a source memory of 1.0 which implies that he had a very good source memory whereas this was not the case. Relatedly, I think chance level should also be defined as 0.5 of the total number of old responses. However, if the**

**authors follow my suggestion, then source memory is not above chance for "unsure old" responses for the whole sample. What does that imply for the interpretation of the results?**

The reviewer raises an excellent point here, as we are sympathetic to it; however, for the reasons detailed below we also believe that it would not be the right measure to use to assess the empirical hypotheses that the present work was guided by. We agree with the reviewer that there is no perfect measure of source memory performance, but for the reasons detailed below we are confident that our measure is the most appropriate for the present paradigm's data. In short, the source memory calculation we used is akin to the standard recognition measure of memory performance (%Hits-%FA) but applied to source memory discrimination, which would require excluding those 'source unknown' responses, since a false alarm in source memory would be a source misattribution (not the lack of source memory). When we look at source memory discriminations between available options of the two sources for participants to choose from, they have a 50% chance or being right/wrong (e.g source correct or source incorrect responses), and this is why we set chance levels to be at 50% in that analysis (as opposed to 33% if assessing it relative to the three response options of correct, incorrect and unknown).

We first address the Reviewer's point about our performance calculation approach that excluded trials in which people reported not remembering any source memory at all (the Reviewer's position was that those trials should instead be included in the calculation). In measuring source memory, like most studies, we wanted to know how accurate people were in _**discriminating between the two sources of information**_ in the experiment. We did not want to know how frequently people remembered correct source information at all, out of all of the possible times that they could have, because, like in real life, this number would presumably be quite low (as the Reviewer calculated)- but artifactually low. This is similar, and applicable, to similar issues in reality monitoring (e.g. clinical studies of auditory hallucinations in schizophrenia patients, for instance), whereby the pertinent question is about understanding how well the source determination is made when a source of the information is available- not instance in which it is not.

The Reviewer's calculation would include all trials of when people had an absence of source memory in a calculation for source memory discrimination, and this would be presenting an entirely different kind of confound. For example, if we all were tested on if we remembered where we learned various items of information in our lives (while already knowing that most of us remember [if we are lucky…] only the information we learned though not the place/person/source of when we learned it). Then consider if the performance measures for our memory measured the few times that we did remember correctly amid _all_ the times when we couldn't remember any source at all (likely most of the time…). Under such calculations, most of us would [falsely] appear to be dense amnesiacs- despite likely having reasonably decent source discrimination capabilities for the times that we could retrieve source memory information.

In the present study, we wanted to know: when people had some memory for source information, how accurate were they in discriminating among those source memories? Therefore, to assess our question, it requires looking at how well people could discriminate between the two possible sources among the times in which they were able to retrieve any source at all- which (contrary to the Reviewer's suggested alternative calculation) necessitates excluding the instance of when they said 'source unknown'. *That is precisely what our calculation did, and that is why it (properly, we contend) excluded the option we provided subjects to respond with of "source unknown".* Our having provided subjects that option to indicate 'source unknown' was an important paradigmatic innovation when we first reported it in 2012, because if not providing it then subjects would be artificially confounding otherwise binary source judgments typically used in studies (correct, incorrect) with instances of both guessing and/or not having any memory at all- that would contaminate those results with major confounds.

Having a lack of source memory is not the same thing as being incorrect about a source memory that one does have- and therein lies the difference between our approaches to calculating source memory. When assessing how people performed those source memory judgments, it was therefore important to only include those times when people had source memory- not the times that they didn't. We believe that the proposed approach in the Reviewers comment would introduce a serious confound and produce an artificial result of floor effects. We are confident that our approach is valid, and it comes from previously published work, which we replicated in entirety in the present findings.

The Reviewer's other, related point was that she suggested that if using her calculation approach of including all of the 'source unknown' trials into the source discrimination calculation, that chance performance should be identified as 50%. We disagree. We believe that this kind of approach also creates another set of confounding issues. For instance, it may not be fair to say that chance should be 50% of the total number of old, as the comment suggested, because this would depend on how each participant differentially distributed their responses, and with the "unknown" option, there is no way chance would be 50%. If including the third source memory response option of 'source unknown' in the calculation, then it could be argued that chance performance among those judgments would be 33%, given that there were 3 source options to have distributed responses to randomly if by chance (correct, incorrect, unknown).

A related problem for the approach suggested by the Reviewer is that in such cases one won't know what the response criterion a subject is adopting when making the 'unknown' response. Some might say 'unknown' only 2 times because they always have some weak feeling of source, whereas another person might say 'unknown' 20 times because they are only going to give a source response if they are sure they would get it right. In that sense, there is not an absolutely correct way of measuring accuracy in this case, as extreme cases can impact outcomes from either approach, as each version of measurement calculation seems to have pros/cons, and each can produce hypothetical edge instances of issues. However, we believe that, for the experimental questions investigated in the present study, our approach is the better one for most closely approximating the accuracy of source memory discrimination between two sources and is more appropriate for excluding confounds and artifacts.

**R3.4: In the 3x2 ANOVA on RTs, for recollection only i5s4 items are included. Why not also i5s5**?

The Reviewer raises a good question. For the initial analyses of behavioral response times, our purpose was to assess replicability and behavioral results of the the preceding 2012 paper that motivated the current investigation, which had originally focused upon the conditions of i4s4, i5s4, and correct rejections (2012, Experiment 1, Neuroimage). The reason for that approach was because it held constant the variable of source memory confidence levels, so that we could be sure that if we didn't see an LPC for the i4s4 condition that it wouldn't be attributable to there being lower confidence in the source judgment than there was in the LPC 'control' condition of 'item5 with source correct'. Later in that study, those findings were independently replicated by its Experiment 2 data (from a 2011 study, Addante et al., *PNAS*), in which a more general measure of i4SC and i5SC was used (each collapsed across both high and low levels of source memory confidence) to verify that the same effects were preserved either way. Thus, we did not include the i5s5 conditions in the RT analysis because we wanted to a) preserve the closest possible analysis for replication, and b) to be rigorous in minimizing any potential confounds from factors of confidence ratings, even though the data has also shown that the same effects are preserved when using the more general measures of the conditions, respectively.

**R3.5: In my eyes, the claim that the behavioral dissociation of the three conditions provides evidence that they reflect distinct cognitive processes is too strong. For example, if we assume a slow recollection process going on, this might lead to slower responses in the item task but to faster responses in the source task as the retrieved source information was just activated at the end of the item task and is therefore more readily available**.

Reviewer makes a good point. For clarity, while oftentimes behavioral dissociations of response times for memory conditions has been taken as evidence reflecting distinct cognitive processes (Yonelinas, 2002), in the present study similar claims that we make are premised not just upon the behavioral dissociations, but upon the converging evidence of observations from the neural data in multiple different measures (hits vs. correct rejections; item familiarity vs context familiarity; each condition of recollection, item familiarity, and context familiarity vs correct rejections; etc.), and as reproduced among several different independent studies.

As to the second part about assuming a slow recollection process going on (*see detailed response to Reviewer 1.7a abov*e), we would note that it is a heavy assumption to make, because we have scant evidence to suggest that the memory condition context familiarity is being supported by any indications of recollection-based processing. That is, it differs reliably from recollection trials in behavioral response times for item recognition, and for source memory judgements, and then also differs reliably and reproducibly in several ERP measures as well. That the source judgments for low confidence recognition also tend to be low confidence judgments (2012) also argue against the premise that even a later process of slow recollection might be driving their trials, because if recollection occurred later it could be assumed to have more high confidence correct source judgments (and it doesn't have that profile). That being said, we have also now updated the Discussion section (Supplemental Materials) to also directly address further details about the relative timing of the responses and the possibility of fast and slow recollection, which we hope provides better balance and support to the interpretations of our findings.

**R3.6: It was not clear to me how the item familiarity time window was defined. In the first set of analyses, it was 400-600 msec whereas it was 300-500 msec in later analyses**.

This is a good question, and we apologize for the prior manuscript being a bit confusing on this front.

For the first ERP analysis of the paper (Hits vs CR), the time window of 400-600 msec was used because the prior findings from this data (Muller et al., 2021) and this contrast (Addante, Ranganath, and Yonelinas, 2012) identified the FN400 effect from 400-600 msec, so this was used  to assess the extent to which the present data set could establish a basic replication of the prior findings of ERP effects.We maintained that time interval for other contrasts in which we sought to assess replicability of prior findings reported for that time when compared to correct rejections.

However, the 300-500 msec time interval was used for a different kind of analysis that directly contrasted what we hypothesized was two different kinds of familiarity processes. This analysis had never been done before, and critically, it was based upon the conventional theory of an FN400 effect as the putative neural correlate of item familiarity (Rugg & Curran, 2008, TICS) that the broader literature has traditionally found from 300-500 msec- so we wanted our foundational analysis to provide a direct test of that traditional view. Thus, we endeavored the novel analyses in the present investigation of the extent to which the two familiarity conditions of item- and context- might be found to be dissociable, using the time interval of 300 to 500 msec because of how common that had been in other historical studies of the FN400 (Rugg et al., 1998, Nature; for Review see Rugg & Curran, 2008).

As stated in the Results section for the analysis directly assessing the two conditions of familiarity (item, context), "*we first wanted to know if the conditions varied during the conventional time window reported for familiarity-based processing of the FN400 (300-500 msec*)". We considered an otherwise negligible difference since the FN400 is reported among slightly different latencies across other studies (Ozubko et al., 2021), and one which could also ensure that effects could be interpreted relative to both the preceding study that motivated it as well as the broader literature of earlier historical findings, too, and were glad to see common results among both approaches across experiments..

**R3.7: It is not clear to me why the between subjects analysis was carried out only for the first two time windows but not the third context familiarity time window**.

The original reasoning for this had been because we were testing the hypothesis of if context familiarity is different from item familiarity during the traditional time intervals of familiarity and recollection. We agree it would be useful to continue that analysis to the later time interval, and have now updated the revised manuscript to include that interval.

**R3.8: I am not convinced that the effect which is claimed to be an LCN effect is not just an LPN effect. In their first argument, the authors say that in other studies the LPN was larger for incorrect source judgments or invariant to source memory, whereas in the current study, the effect is larger for correct source judgments. However, I do not think that the studies are comparable as response times were considerably faster in the other studies. Therefore, the process underlying the effect (LPN or LCN) might have contributed more to source judgments in the current study than in the other studies**.

This is a good question about the interpretation of the results as an LPN, and was similarly raised by each of the other two reviewers above. The issue of the interpretation of the ERP effects of context familiarity reflecting an LPC has been addressed in earlier responses to Reviewers above (*see 2.2, 1.9, 1.7, 1.1*). Specifically responsive to this particular comment, it is unclear what is meant by the comment here: In the present study, it is not just that the effect is larger for correct source judgments: it is that it is *only present and observable at all for correct source judgments*; and furthermore, only for *some* correct source judgments (low confidence recognition, 4's) and not other source judgments (high confidence recognition, 5's). It is also not clear to us what is meant by the relation of response times to ERP effects in the comment, and how that would explain any differential ERP results of source memory within or across studies- though we would be glad to respond better if provided more detail.

As to whether or not the observed LCN effect is or isn't just an LPN effect (*see previous Responses to Reviewers 2.2, 1.9, 1.7, 1.1*), the authoritative Review paper about the LPN effect (Mecklinger et al., 2016) has in fact itself argued that the present findings (2012) were *not* the same thing as our LCN and should not be confused as such.

> (page 631) *"Even though these late negativities have been associated with attempts to reconstruct memory-stored information, processes usually ascribed to the LPN, the non-posterior scalp distributions suggest that it is unlikely the same ERP component as the LPN. The same objection holds for the late negativities reported for young adults in the* ***studies by Addante et al. (2012a****),*

*Leynes and Zish (2012), Leynes et al. (2013) and, for amnesia patients, by Addante et al. (2012b). All of these studies report negativities that are larger for studied than unstudied information but due to a lacking posterior scalp distribution t**hese components are unlikely the same ERP component as the LPN.**" [emphasis added]*

And (page 634): *"**Addante et al. (2012a**) examined ERP correlates of source memory for items that were recognized with high or low confidence. A parietal old/new effect (the ERP correlate of recollection) was obtained for correct source judgments, but only for items recognized with high confidence. Correct source judgments for items that were less confidently recognized elicited a late but broadly distributed negativity, **resembling the LPN.** It could be argued that the low confident item memory in this study constitutes a paradigmatic case for a high need for continued evaluation of source-specifying memory information and LPN generation. However, it could also be argued that the low confidence for the item judgments is a direct consequence of the failure to retrieve source specifying information. Irrespective of this interpretational ambiguity, **the rather broad scalp distribution and the early onset of the negativity do, however, not necessarily support an LPN interpretation of this effect**". [emphasis added].*

**R3.9: In their second argument, they say that an LPN should have been observed also in the item familiarity and the recollection condition. However, if participants could make a correct item judgment on the basis of familiarity or recollection, why should they (necessarily) search for source information**?

If participants had made a correct item judgment on the basis of item familiarity, and could not recollect the source of the event, then presumably they would be engaging in a controlled search process of their memory to see if they could find that source information. That is why one could expect to see an LPN observed in the item familiarity condition: because they made an accurate item recognition but then reported that they did not remember the source. In fact, they took longer to report that source judgment of 'unknown', suggesting that they were indeed searching more for that information unsuccessfully. That seems like a condition of a 'controlled search', yet it exhibited no evidence of an LPN. Accordingly, if recollecting, they'd also presumably be searching for memory information, too (successfully, this time), and one could surmise that could evince an LPN, too.

**R3.10: The third argument of the topographical dissociation is also not very strong as the LPN can indeed vary in its topography depending on task and domain-specific factors (Mecklinger et al., 2016, Neuroscience and Behavioral Reviews). The same holds presumably also for the timing.**

The Reviewer raises an interesting point. The point that the Reviewer is referring to from the manuscript (of the topographic and temporal differences between where the LCN effects were observed to manifest in the present study, as compared to where the LPN effects have been reported as observed in other studies), was one particular line of evidence among several different lines of converging evidence that addressed the bigger question of whether or not the observed LCN effects might be the same as LPN effects that others have observed in different studies about memory search. We thank the Reviewer for raising this point, and appreciate the opportunity to address it with additional points beyond those already made in responses above to *Comments (R1.1, 1.7, 1.9, 2.2, 3.8, and 3.9*).

The Reviewer's point here claimed that the Late Parietal Negativity ERP effect (LPN) "***can indeed vary in its topography…and timing***". But 'topography' and 'timing' are both core, defining features of the Late Parietal Negativity effect- without which the ERP effect could only be described as a mere "Negativity". If it can vary in its timing and topography, then there is nothing late nor parietal about the Late Parietal Negativity- That is not very helpful as a descriptor, then, of the neural correlates of memory operations identified with an LPN. If the field were to adopt such an approach, then the "LPN" would thus be merely a generic negative-going wave happening at some place on the scalp at some time during memory (note: we do not think it is so; but that is the limitation of the proposed characterization of it in this case). We would completely understand (and expect) some natural variability in the LPN effect as is the case for nearly all neural correlates of memory, but would also find it more compelling of an account if only one of its three features were found to be variable in a particular measure, not two-thirds of its defining features.

The LPN is defined quite nebulously, operationalized broadly as a merely a "reconstructing the past" or a "controlled search process" (Mecklinger et al., 2016, 2006), which, colloquially, only means "trying to remember", which should occur on every memory retrieval trial to one extent or another. Being told here that it can vary across any temporal or topographical extent only makes it more nebulous and less specific towards being a useful cipher of memory process operations. Due to the poor operational definitions available to the LPN, it is just as possible that it could instead be reflecting the contribution of at least some context familiarity being retrieved in the absence of cohesive recollection or item familiarity (e.g. Strunk et al., 2016, *Neuropsychologia*). If the LPN is to be useful as an ERP effect towards understanding memory, then we ought to use it as linked to at least some of its defining features, not just as a generic negative ERP effect (many other ERP effects are also negative-going in polarity, too, but reflect very different psychological constructs), and that should be falsifiable in saying the instances when it is, or would not be, occurring. We can not just attribute all negative-going ERP effects of memory as an LPN, no more than we can attribute all positive-going ERP effects of memory as an LPC.

Since the Reviewer's Comment here specifically mentioned the aspects of topography and timing, ***in fact, the authors of the Review paper cited by the Reviewer's comment (Mecklinger et al., 2016, Neuroscience and Biobehavioral Reviews) wrote***

***specifically in that Review Paper about how the present effects of the LCN are actually *not* likely to be reflecting an LPN- particularly based upon factors such as topography and timing***. For instance, Mecklinger et al., 2016 wrote the following statements to this effect:

> (page 631) *"Even though these late negativities have been associated with attempts to reconstruct memory-stored information, processes usually ascribed to the LPN,* ***the non-posterior scalp distributions suggest that it is unlikely the same ERP component as the LPN****. The same objection holds for the late negativities reported for young adults in the* ***studies by Addante et al. (2012a),*** *Leynes and Zish (2012), Leynes et al. (2013) and, for amnesia patients, by Addante et al. (2012b). All of these studies report negativities that are larger for studied than unstudied information but* ***due to a lacking posterior scalp distribution these components are unlikely the same ERP component as the LPN.****" [emphasis added]*
>
> And (page 634): *"****Addante et al. (2012a) examined ERP correlates of source memory for items that were recognized with high or low confidence****. A parietal old/new effect (the ERP correlate of recollection) was obtained for correct source judgments, but only for items recognized with high confidence. Correct source judgments for items that were less confidently recognized elicited a late but broadly distributed negativity, resembling the LPN. It could be argued that the low confident item memory in this study constitutes a paradigmatic case for a high need for continued evaluation of source-specifying memory information and LPN generation. However, it could also be argued that the low confidence for the item judgments is a direct consequence of the failure to retrieve source specifying information. Irrespective of this interpretational ambiguity, t****he rather broad scalp distribution and the early onset of the negativity do, however, not necessarily support an LPN interpretation of this effect****". [emphasis added].*

Furthermore, other lines of evidence we have provided weighing against the LPN account of our findings comes from testing a direct prediction made about the LPN. The Mecklinger et al LPN Review (2016) made the specific prediction that the LPN should be present "*not necessarily tied to successful memory retrieval but is also observed during misses…"* (pg. 635). We did not find that prediction held in our data, as the present findings directly dissociated ERP effects of misses from the ERP correlates of context familiarity, and this thus adds to the line of evidence weighing against an interpretation of the effects as an LPN.

Any alternative account of the present findings needs to be able to explain having found these effects repeatedly across several different studies, measures, and decades. It needs to be able to explain by what memory process people are arriving at these curious combinations of responses, as currently the pattern of reproducible results can not be accounted for by either single or dual process models of recognition. We have gone to extraordinary lengths testing many different control analyses in attempts to falsify our interpretations. We welcome any other lines of evidence, derived from testable and falsifiable hypotheses. Until that time, we report what we have observed, and look forward to future studies inspired by these findings.

**Minor points**:

**R3.a: Where participants compensated for their participation?**

Good question. We forgot to include this in the earlier version of the manuscript, as this information had been noted in the original paper reporting the data set by Muller, Siranni, & Addante (2021, European Journal of Neuroscience). We have now added it to the Supplemental Materials, which states: "*Participants were recruited through a combination of methods including advertisements placed around CSUSB or through the schoolwide research pool SONA. Participants recruited through advertisements were paid $10 an hour for sessions that lasted approximately 2 hr*".

**R3.b: What was the range of the years participants had been speaking Englis**h?

The Method section states: "*Most participants spoke English as their first language (N = 47), and the 15 whom indicated speaking a different language first had been speaking English for an average of 16.73 years (SD = 4.74)*". Unfortunately, we can not find the background logs of the data for the min/max range of the years that participants had been speaking English, as that data was collected nearly five years ago and both of the researchers involved in collecting the data for that project (AM, Masters Thesis; RJA, Principal Investigator) have since moved from California to the University of Arizona and Florida Institute of Technology, respectively, and the location of that data was lost in amid the moves (which occurred during the Covid-19 pandemic). We have looked extensively for it, and though we can not find it now, it may potentially emerge later. Importantly, we are confident that language was not a barrier to the study nor a viable account of the results.

**R3.c: With respect to power of the study: I understand that there was no a priori power analysis as the study was designed for another purpose. However, it would be nice to read some words about the sensitivity of the sample to detect the effect sizes relevant in the study**.

The Reviewer raises a great point. We have now included a power analysis in the revised manuscript's Supplemental Materials, where it describes the results provided below, which we believe adds further strength to the findings.

The sample of data of the present study demonstrated sensitivity to detect the effects of context familiarity originally discover by the first study conducted in 2012, by virtue of it having reproduced those findings of differences between context familiarity and recollection in a sample (N = 54) that was more than double the original sample size (N

= 25). However, it remains unclear what the precise statistical power of the study was to detect the full range of potential differences. Therefore, a post hoc power analysis was conducted using G*power version 3.1.9.7 (Faul et al., 2007) in order to estimate the sensitivity of the sample to detect the effect sizes reported in the present findings.

For the investigation's finding of replicability of the 2012 findings (ERPs for context familiarity vs correct rejections at Cz from 800-1200 msec), using a significance criterion of a = .05, and the observed effect size (.486) in the in the available subjects (N = 27), the power was .681. For the main finding dissociating item- and context-familiarity ERPs within-subjects(N = 11, Cohen's d = .762), it had a power value of .626. The between-subjects effects for the same contrast (N = 9, N = 17, effect size of 1.14) had a power of .755. Behaviorally, the main effect of item recognition response time differences for item- and context-familiarity conditions (N = 38, a = .05, Cohen's d = .481) had a power of .823. When high confidence recognition responses were excluded from the item familiarity condition in the same contrast, there was a power of .530; whereas the ERP correlate of the same control analysis (Experiment 2, Supplemental Materials; N = 12, effect size of .569) had a power of .436.

Overall, these power analyses indicated that the study was somewhat underpowered to detect some of the effects which were observed, yet we still found consistent reliable results, which our additional studies demonstrated were independently replicable. Thus, while the present study was fairly sufficient in its sensitivity to detect the effects of interest, the results of the power analyses suggest that there may likely be more, or stronger, effects existing to be identified in future studies that are better powered. Together, this suggests that we are 'leaving money on the table', so to speak, and that there may very well be far more to discover about context familiarity effects in future research studies designed to be better-powered to detect the full extent of effects.

**R3.d: I do not think that the Procedure should be described in the supplemental materials (SM). This also holds for those parts of the Discussion which are marked as SM**.

This is an exceptionally good point, which we agree with in totality. However, the journal has very specific limitations on word count (~5500) that would be exceeded if these sections were added to the main text. If the Editor were to create an exception of permission to exceed the parameters for word count/length, then we would be more than happy to add what we, too, feel are very important sections to the main text.

**R3.e: I would also like to read some more details about the procedure: What was the timing of the trial procedures? Were trials terminated by response? Was the source memory question presented in each trial or only for old items or when an old response was given?**

These are good questions. Since the present study was a secondary re-analysis of a previously-published data set, we had cited it in the Method section for brevity (Muller et al., 2021, EJN). We have now updated the revised manuscript to have this information included in the Supplemental Materials.  Trials were terminated by subject response. Source memory questions appeared for every trial. As noted in the Muller et al (2012) study, every 10th trial there was also a metacognitive judgment made by participants as well. The figure of the procedure provided below is an excerpt from the original report by Muller, Sirianni, & Addante (2021) and provides additional detail of the procedure, and full details are also written in the publication linked to below if interested (https://onlinelibrary.wiley.com/doi/full/10.1111/ejn.14935).

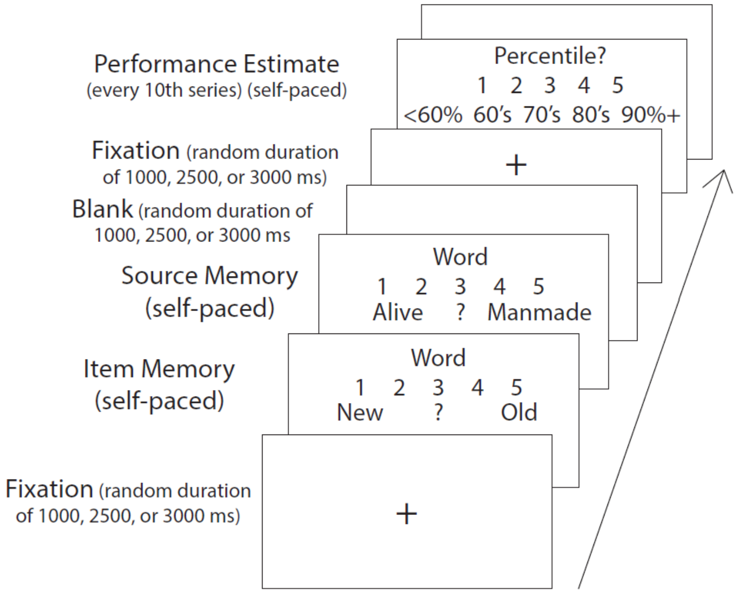


**R3.f: During the recognition test, participants saw almost double the number of old items compared to new items. Might an induced bias to respond "old" be a problem for the interpretation of the results? Please also relate this to other studies investigating ERP old/new effects**.

In regard to the proportion of targets and lures, the proportion of targets to lures in the present study was designed to match the same proportion used in the preceding study that motivated the current work (Addante, Ranganath, & Yonelinas, 2012).  The reason that the study used different numbers of trial types for old and new items in this experimental design is because this approach is nearly always used by studies to equate the number of each class of items (e.g., ~100 trials of old items from source1, ~100 trials of old items from source2 and ~100 trials of new items). We are not aware of any studies that would indicate that this type of proportion of targets and lures would alter accuracy, but rather it just increases a bias to say old to both old and new items (e.g., Rattcliff et al., 1992). Ian Dobbins and Mike Miller have done several studies of extreme varying of the proportion of lures, showing it is really hard to get subjects to change their criterion even when you'd think they really would do it. The worst case scenario is that people might have a slightly higher response bias to say old, but that would likely only water down the observed effects, in that it'd be watering down the pool of "old" trials but the present findings had no trouble detecting differences between old and new items, and the results for old and new items are very consistent with the other findings in the literature. Overall, we do not see a reason to assume any substantive impact of target/lure proportions on the pattern of results the present study focuses upon.

**R3.g: I would highly recommend sharing data and scripts in a publicly accessible format**.

We are committed to following the journal's policies on sharing data, and to provide the data and scripts in accord with the policies of the journal. We are working on hosting the data online to facilitate that process amid the very large sets of complex data.

**R3.h: For the case of exploratory analysis, Type I error correction for multiple comparisons needs to be applied**.

This has now been done for all of the exploratory analyses. Of note is that all of the behavioral results remained unchanged after application of strict Bonferroni corrections.

**R3.i: Please provide a justification for the choice of the electrode sites analyzed**.

The results and methods sections note that the choice of electrode sites for analyses were based upon a priori hypotheses derived from the same sites reported in prior studies that we sought to replicate in the present work. For instance, in the Results section describing the analysis of hits vs correct rejections, it states: "...*applied here to three different locations and latencies selected based upon prior findings* [1,37]*…For the first analysis, we used a basic contrast of measuring ERPs for hits vs. correct rejections in every subject (N=54) focusing on same sites as the previous reports of context familiarity* [1,37]." Later, when describing the results section contrast ERPs for context familiarity vs item familiarity, it states: "*we first wanted to know if the conditions varied during the conventional time window reported for familiarity-based processing of the FN400 (300-500 msec,* [42,43]*, focusing on site Cz as the site of maximal activation for these in the current and prior study* [1]*.*". When performing the direct test between ERPs for context familiarity and recollection, the Results section stated: "*A 3x3 repeated measures ANOVA for factors of memory condition (recollection, context familiarity), time interval (400-600ms, 600-900ms, and 800 1200 msec), and site (P3, Cz) was performed based upon the factors identified in the preceding analyses and prior reports dissociating context familiarity from recollection-based processing* [4]*….These findings were then explored with planned t-tests to characterize the nature of the interaction effects at the sites of the maximal activations, respectively (P3 for recollection, Cz for context familiarity)*."

**R3.j: In the Discussion (section 4.2), familiarity for pictures and faces is considered as instances of context familiarity. I do not agree. Of course, pictures and faces can serve**

**as context in some instances, but this holds also for words or concepts. However, in most experimental investigations, pictures of objects or faces are presented as items**.

The reviewer makes a good point here that there may be varying opinions on how to interpret certain data on memory for faces, pictures, words, and concepts. We have updated that section of the Discussion to soften this consideration and specifically noted the Reviewer's concern as noted below:

> "*This patient was deficient in familiarity for items (words) but showed intact abilities of familiarity for pictures and faces [84-87], which we consider here as broadly representing kinds of context in episodic events (though which may also be treated as an item in certain studies with other contexts), and this could be seen as performance consistent with the patient's intact parahippocampus service of context familiarity*"

**R3.k: In 4.3.2, towards the end of the first paragraph: change 'become' to 'becomes'.**

This has been updated accordingly.

**R3.l: Figure 2: Please align the order of the description of the condition to the labeling of the x-axis. The coloring of the violin plots is very light**.

The figure caption and violin plots have been updated accordingly.
